# Panmap: Scalable phylogeny-guided alignment, genotyping, and placement on pangenomes

**DOI:** 10.64898/2026.03.29.711974

**Authors:** Alexander M. Kramer, Alan Zhang, Nicolas Ayala, Bianca de Sanctis, Lily Karim, Angie S. Hinrichs, Sumit Walia, Yatish Turakhia, Russell Corbett-Detig

## Abstract

Pangenomes capture population-level variation but remain computationally challenging at scale. We present Panmap, a tool that leverages evolutionary structure to place, align, and genotype sequencing reads against mutation-annotated pangenomes containing up to millions of genomes. Panmap introduces a phylogenetically compressed k-mer index that stores only sequence differences along branches, enabling efficient comparison of reads to both sampled genomes and inferred ancestors. This approach reduces index size by up to 600-fold and construction time by over three orders of magnitude relative to existing tools. Panmap places a 100× coverage SARS-CoV-2 sample onto 20,000 genomes in 0.4 seconds and onto 8 million genomes in under two minutes. Furthermore, it enables accurate haplotype identification and abundance estimation in metagenomic samples and sensitive placement of ancient environmental DNA without prior alignment. Our approach makes large-scale pangenomes directly amenable to read mapping, genome assembly, alignment-free phylogenetic placement, and metagenomic analysis.

## Main

Pangenome approaches have emerged as a powerful alternative to single-reference methods, jointly leveraging many reference genomes to capture greater genetic diversity and improve variant identification. Because they encode known variants within populations, species, or clades (Marschall et al. 2018), reference pangenomes enable more sensitive and accurate analyses than single-reference approaches (Jana Ebler et al. 2020) and have become essential in fields such as human genomics, infectious disease surveillance, and agriculture (Sirén et al. 2021; Zhou et al. 2022; Rice et al. 2023).

Most pangenome methods represent variation as sequence graphs, encoding genomes as paths through networks of shared and divergent sequences (Liao et al. 2023). Tools like VG (Erik Garrison et al. 2018), Progressive Cactus (Joel Armstrong et al. 2020), and Minigraph (Li et al. 2020) exemplify this framework. However, graph-based approaches face significant scalability challenges. They often require extensive memory and time for read alignment and variant identification, especially for large microbial pangenomes that may contain thousands or millions of whole genomes. In addition, sequence graphs typically represent variation structurally but do not explicitly encode the evolutionary history relating sequences, limiting opportunities to leverage ancestral sequence information during analysis.

An alternative paradigm, used by tools like UShER and *tskit*, represents genomic variation as genealogical structures, encoding variants as mutations along evolutionary branches (Kelleher et al. 2018; McBroome et al. 2021; Yatish Turakhia et al. 2021). The Pangenome Mutation-Annotated Network (PanMAN) format uses this approach for lossless pangenome representation, capturing point mutations, indels, and structural variants as mutation-annotated genealogies. These genealogies may be single trees for minimally recombining species, or phylogenetic networks for more complex evolutionary histories (Walia et al. 2026). By encoding ancestral sequences at internal nodes, PanMAN achieves superior compression while providing a richer reference structure that includes both sampled diversity and inferred ancestral sequences at the internal nodes in the network. However, fully leveraging such genealogically structured pangenomes for common tasks like read alignment, phylogenetic placement, and metagenomic inference requires new methods to efficiently index and utilize mutation-annotated phylogenetic networks as a reference object.

Here we present Panmap, a tool that indexes *k*-mers in PanMANs to perform phylogenetic placement, read alignment, genotyping, and mixed sample haplotype deconvolution from sequencing reads. We apply it to pathogen genome assembly, wastewater metagenomic surveillance, and phylogenetic placement of ancient environmental DNA, demonstrating equivalent or better performance while scaling to reference pangenomes intractable for existing tools.

## Results

### Panmap overview

Panmap builds a compact k-mer index from the sequence content and phylogenetic structure of a PanMAN, supporting placement, alignment, genotyping, consensus assembly, and metagenomic deconvolution from sequencing reads. It provides two processing modes (Figure 1). In single-sample mode, *k*-mer seeds (see Methods) from all reads are scored jointly across the phylogeny to find a close reference haplotype, which may be a sampled genome or an inferred ancestor. Panmap can then optionally align reads to that sequence and produce a consensus assembly with genotype calls. In metagenomic mode, reads are scored independently, yielding per-read node assignments and haplotype abundance estimates (see Methods).

**Figure 1.**
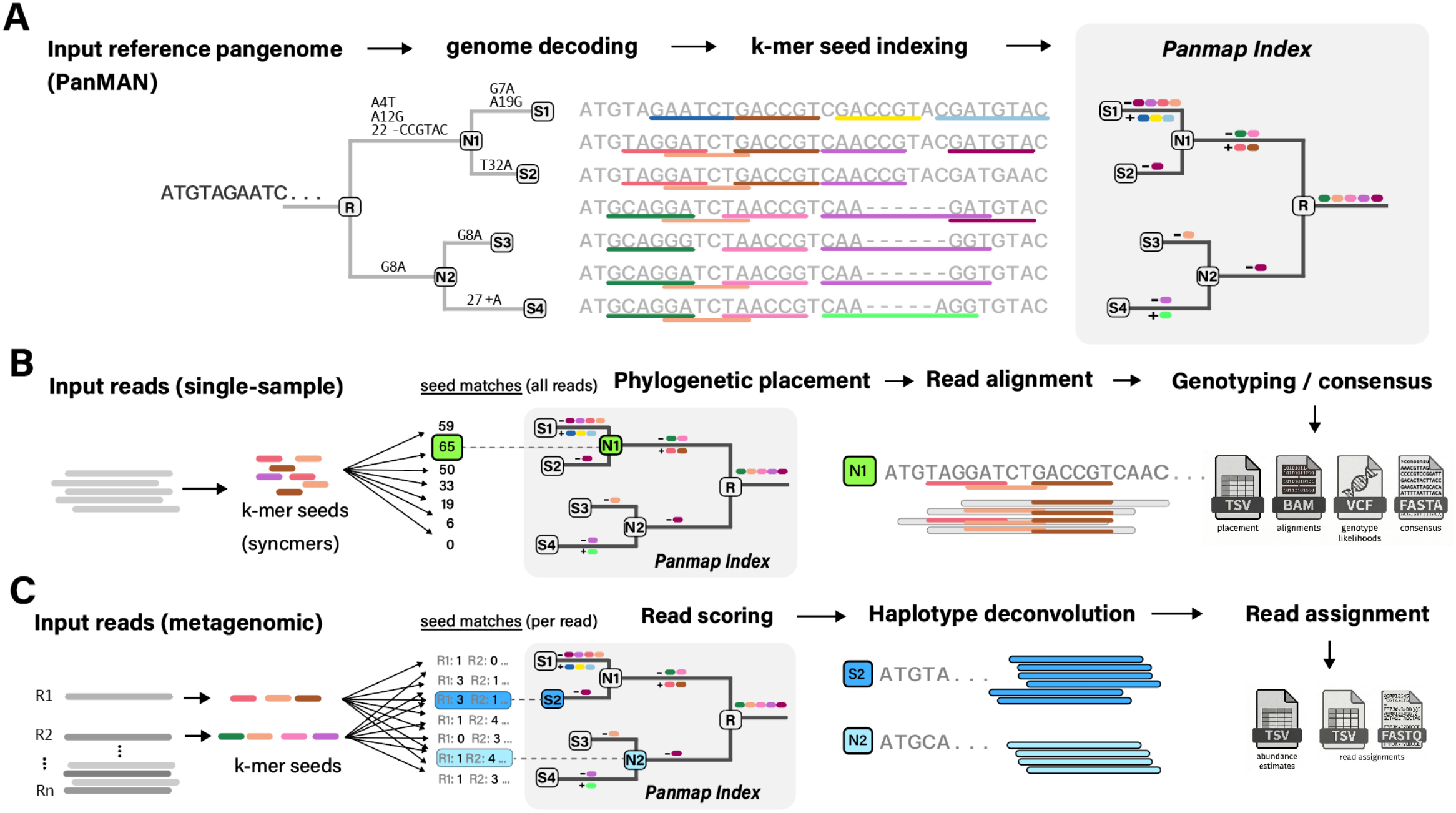
Overview of Panmap’s indexing and analysis modes. (**A**) Pangenome indexing. Panmap takes a reference pangenome in PanMAN format (a mutation-annotated phylogeny encoding SNPs, insertions, and deletions along each branch), decodes the full genome sequence at each node, extracts *k*-mer seeds, and stores only the seed differences between parent and child nodes in a phylogenetically compressed index. (**B**) Single-sample mode. Input reads are decomposed into *k*-mer seeds, and Panmap scores all seeds collectively against every node in the index during a single tree traversal to identify the best-matching reference (here, node N1 with 65 seed matches). Raw counts are shown, though Panmap uses a containment metric in practice; see Methods). Reads are then aligned to the placed node’s genome sequence (decoded from the PanMAN), and Panmap produces a placement report (TSV), read alignments (BAM), genotype likelihoods (VCF), and a consensus assembly (FASTA). Each stage is optional: for example, Panmap can report only the phylogenetic placement without proceeding to alignment or assembly. (**C**) Metagenomic mode. Each read is scored independently against the index, and per-read scores are used to identify the reference haplotypes present in a mixed sample (here, S2 and N2). Reads are deconvolved by competitive assignment to their best-matching haplotypes, and Panmap estimates relative abundances and outputs abundance estimates (TSV) and per-read assignments (FASTQ).

### Phylogenetic compression yields indexes up to 600× smaller than alternatives

Panmap’s efficiency results from its phylogenetically compressed *k*-mer index. Because closely related genomes share most k-mer seeds, representing genomes in a phylogeny allows Panmap to store only the seed differences along each branch rather than storing seeds independently for every genome. Panmap traverses the entire reference PanMAN phylogeny once and encodes only the seed differences between parent and child nodes. These delta encodings capture changes in k-mer seed presence along each branch, exploiting the similarity among closely related genomes for dramatic compression and extremely efficient computation of the index.

Panmap’s index outperforms alternatives in construction time, index size, and memory required. We compared the Panmap index to IPK/EPIK (Nikolai Romashchenko et al. 2023), which builds phylo-*k*-mer databases for phylogenetic placement, and VG Giraffe (Erik Garrison et al. 2018; Sirén et al. 2021), which indexes pangenome graphs for read alignment (Table 1, Table S2). For 4,000 RSV genomes, Panmap’s 5.7 MB index built in 4 seconds, versus VG Giraffe’s 3.5 GB (614× larger) in 6 hours (5,580× slower) and IPK/EPIK’s 2.1 GB (368× larger) in 3 hours (1,005× slower). For 20,000 SARS-CoV-2 genomes, Panmap’s 7.3 MB index was 614× smaller than VG Giraffe’s 4.5 GB and built in 3 seconds versus 7.5 hours. For *M. tuberculosis,* Panmap indexed the 400-genome pangenome about 520× faster than VG, but the compression advantage was less pronounced (240 MB vs 500 MB, 2.1× smaller), likely because the *M. tuberculosis* genomes included in the PanMAN are not as closely related as for the other pathogens considered. IPK/EPIK failed to index both the SARS-CoV-2 and *M. tuberculosis* pangenomes. We performed all of the above experiments with one thread each, though Panmap also supports multithreaded indexing and placement (Figure S1). These results imply that Panmap will scale favorably as data sets continue to grow in size.

**Table 1.**
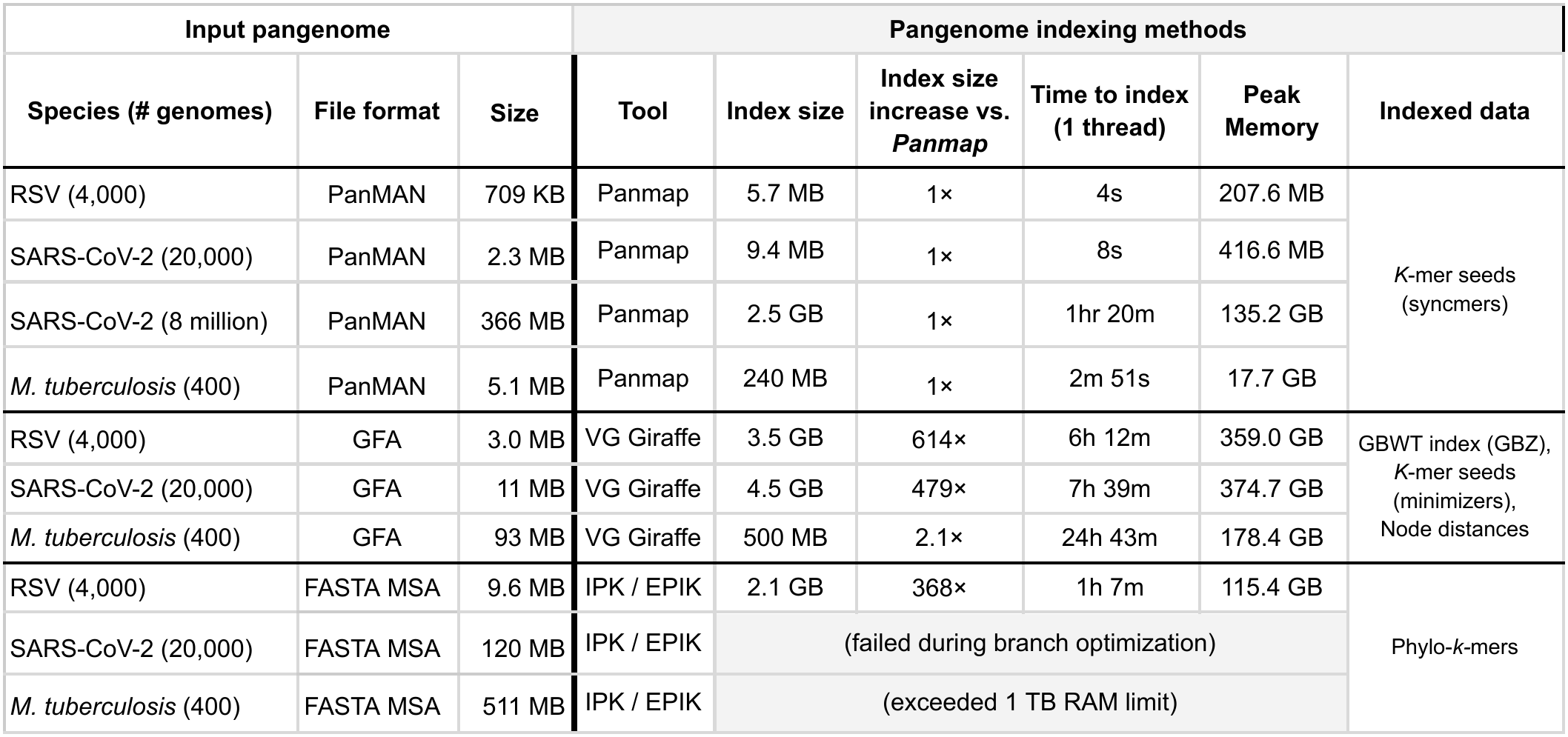
For three pathogen species, we indexed pangenomes with Panmap, VG Giraffe, and IPK/EPIK. We measured the construction time, memory usage, and size after GZIP of the indexes. The Panmap index records *k*-mer seed differences across all genomes and is used for phylogenetic placement and abundance estimation. For read alignment to a pangenome graph, VG Giraffe indexes *k*-mers (minimizers), a graph Burrows-Wheeler transform (GBWT), and node distances. IPK builds the phylo-*k*-mer index used by EPIK for phylogenetic placement. All programs were run with one CPU core on a dedicated computing cluster node and allowed up to 1 TB of runtime memory. We built all Panmap indexes in single-sample mode with default parameters (k=19, s=8, l=3) for all species except *M. tuberculosis*, for which we used k=51, s=8, l=1). Metagenomic mode’s index is nearly identical but also includes seed orientations, which incurs a small size overhead (Table S1).

| **Input pangenome** |  |  | **Pangenome indexing methods** |  |  |  |  |  |
|---|---|---|---|---|---|---|---|---|
| **Species (# genomes)** | **File format** | **Size** | **Tool** | **Index size** | **Index size increase vs. *Panmap*** | **Time to index (1 thread)** | **Peak Memory** | **Indexed data** |
| RSV (4,000) | PanMAN | 709 KB | Panmap | 5.7 MB | 1× | 4s | 207.6 MB | *K*-mer seeds<br>(syncmers) |
| SARS-CoV-2 (20,000) | PanMAN | 2.3 MB | Panmap | 9.4 MB | 1× | 8s | 416.6 MB |  |
| SARS-CoV-2 (8 million) | PanMAN | 366 MB | Panmap | 2.5 GB | 1× | 1hr 20m | 135.2 GB |  |
| *M. tuberculosis* (400) | PanMAN | 5.1 MB | Panmap | 240 MB | 1× | 2m 51s | 17.7 GB |  |
| RSV (4,000) | GFA | 3.0 MB | VG Giraffe | 3.5 GB | 614× | 6h 12m | 359.0 GB | GBWT index (GBZ),<br>*K*-mer seeds<br>(minimizers),<br>Node distances |
| SARS-CoV-2 (20,000) | GFA | 11 MB | VG Giraffe | 4.5 GB | 479× | 7h 39m | 374.7 GB |  |
| *M. tuberculosis* (400) | GFA | 93 MB | VG Giraffe | 500 MB | 2.1× | 24h 43m | 178.4 GB |  |
| RSV (4,000) | FASTA MSA | 9.6 MB | IPK / EPIK | 2.1 GB | 368× | 1h 7m | 115.4 GB | Phylo-*k*-mers |
| SARS-CoV-2 (20,000) | FASTA MSA | 120 MB | IPK / EPIK | (failed during branch optimization) |  |  |  |  |
| *M. tuberculosis* (400) | FASTA MSA | 511 MB | IPK / EPIK | (exceeded 1 TB RAM limit) |  |  |  |  |

### Panmap enables phylogenetic placement directly from sequencing reads

Phylogenetic placement positions query sequences within existing reference phylogenies, enabling sample identification and phylogenetic profiling without de novo tree reconstruction (Lucas Czech et al. 2022). Most existing approaches require some form of preprocessing before placement. Alignment-based methods, such as EPA-ng (Pierre Barbera et al. 2019) and pplacer (Matsen et al. 2010), require queries to be pre-aligned to a reference multiple sequence alignment. EPIK (Nikolai Romashchenko et al. 2023) enables alignment-free placement using phylo-k-mers but still requires assembled query sequences. UShER (Yatish Turakhia et al. 2021) achieves rapid maximum-parsimony placement on mutation-annotated trees but requires VCF input derived from prior variant calling. For ancient DNA, pathPhynder (Martiniano et al. 2022) places samples onto phylogenies from pre-aligned BAM files, while euka (Vogel et al. 2023) and Soibean (Vogel et al. 2024) use VG-format pangenome graphs for taxonomic detection and species-level identification from ancient environmental DNA, but are restricted to curated mitochondrial references for select taxa. Because all these methods depend on a prior alignment, assembly, or variant calling step, they cannot extract phylogenetic signal from reads that fail to map. Panmap overcomes this limitation by performing alignment-free phylogenetic placement directly from sequencing reads. This allows Panmap to use reads that would otherwise be discarded, increasing sensitivity in samples with low endogenous content or highly degraded DNA.

### Panmap achieves high placement accuracy across coverage levels

Phylogenetic placement using Panmap’s single-sample mode is fast and sensitive for simulated and real sequencing data, remaining accurate even for samples with very low coverage depth (Figure 2). For simulation experiments, we selected 70 leaf nodes at random in each PanMAN, applied random SNPs and indels at two mutation rates to generate new descendant genomes, simulated short reads based on each novel genome, then placed the collection of reads for that sample with Panmap. For real data experiments, we chose up to 100 PanMAN leaf nodes per species, downloaded corresponding FASTQ data from the Sequence Read Archive, then removed each chosen node from the PanMAN before placing the reads with Panmap. For both experiments, we quantified placement accuracy as the difference in parsimony score between Panmap’s placement and the sample’s original position in the reference phylogeny before removal (placed minus original). A difference of zero indicates exact recovery of the original placement; positive values indicate Panmap placed the sample less parsimoniously, while negative values reflect a more parsimonious placement. For easier interpretation, we adjust each final score to max(0, score) so that a score of zero indicates an equivalent or better placement by Panmap.

**Figure 2.**
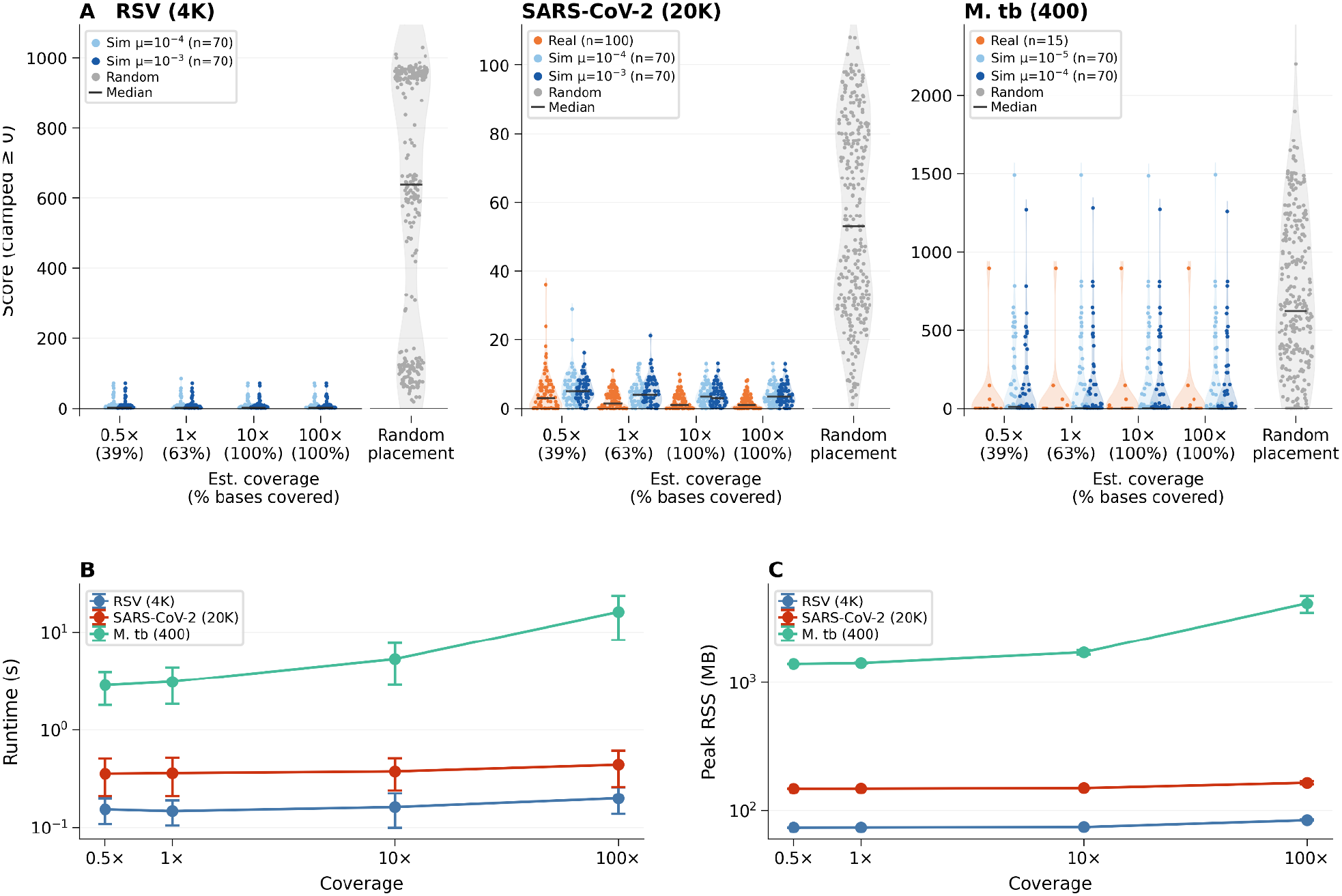
Phylogenetic placement accuracy and performance of Panmap on simulated and real sequencing data. (**A**) Placement accuracy. For each of RSV (4,000-genome PanMAN), SARS-CoV-2 (20,000-genome PanMAN), and M. tuberculosis (400-genome PanMAN), we selected 70 leaf nodes, applied random mutations at two species-specific mutation rates to generate descendant genomes, simulated 150 bp paired-end reads at four estimated coverage levels (0.5×, 1×, 10×, 100×), removed each node from the tree, and placed the reads with Panmap (70 unique samples × 2 mutation rates × 4 coverages = 560 placements per species). We performed the same experiments with real data. Violin plots show the distribution of parent-relative placement scores; a score of zero indicates placement at least as parsimonious as the original position. The gray violins show the distributions of random placements for comparison. Under Poisson sampling, estimated coverage levels of 0.5×, 1×, 10×, and 100× correspond to approximately 39%, 63%, ∼100%, and ∼100% of the genome covered by at least one read, respectively. (B) and (C) show runtime (wall-clock seconds) and peak memory usage as a function of coverage, colored by species.

For SARS-CoV-2, RSV, and M. tuberculosis, Panmap achieved accurate placement across all coverage levels. On simulated SARS-CoV-2 data (n=140, two mutation rates), placement accuracy was stable across coverages, with a median score of 3–5 mutations from the true phylogenetic position. For RSV, simulated placement (n=140) achieved a median score of 2–3 across all coverages, and real RSV data (n=20) had a median score of 4–5. Phylogenetic placement of individual SARS-CoV-2 and RSV samples completed in 0.3–0.7 seconds across all coverage levels, using 148–183 MB of RAM. For the 400-genome M. tuberculosis PanMAN, placement on real (n=15) and simulated data (n=140) achieved a median score of 0–2 mutations.

Panmap achieves accurate placement even with very short reads. On simulated 100× coverage data, RSV placement accuracy was stable across all read lengths down to 10 bp (Figure S2, left). For SARS-CoV-2, where the larger pangenome presents greater ambiguity, linked syncmers (l > 1) substantially improved placement at all read lengths (Figure S2, right). This demonstrates that chaining syncmers into linked seeds (Panmap defaults to l=3) can provide additional specificity for accurate placement on larger, more diverse pangenomes, even when individual reads are shorter than typical Illumina output.

We additionally benchmarked placement onto a PanMAN containing 8 million SARS-CoV-2 genomes. A single sample at 100× coverage requires ∼108 seconds (Figure S3), with more than half of that time spent loading and initializing the index. For large PanMANs, we therefore recommend processing samples in batch mode (--batch) to amortize this overhead. Using 8 threads, Panmap placed a batch of 70 SARS-CoV-2 samples at 100× coverage in ∼8.5 minutes (∼8 samples per minute), with a peak RAM usage of ∼51 GB.

### Phylogenetic placement enables scalable pangenome-guided read alignment

Read alignment to reference sequences is a foundational task in genomic analysis (Reinert et al. 2015; Kim et al. 2020; Alser et al. 2021). In single-sample mode, after *k*-mer based phylogenetic placement, Panmap aligns all input reads to the placement taxon’s genome with the Minimap2 library (Li et al. 2018). The default alignment mode is fast and achieves high mapping rates in most cases. We also provide an optional more sensitive alignment mode that uses classic ancient DNA parameters for bwa-aln (Li and Durbin 2009), which performs well when the input sequences are extremely short (<70bp) or highly damaged, as is typical of ancient DNA (Oliva et al. 2021).

Panmap’s single-sample mode uses phylogenetic placement to identify an optimal reference haplotype, then aligns all reads to that single sequence. For applications where a full graph representation is unnecessary, this approach substantially accelerates pangenome-based read alignment. While pangenome graphs can accommodate tens or hundreds of large genomes (e.g., human), they struggle with thousands or millions of samples, even for species with small genomes (Walia et al. 2026).

We compared Panmap to VG Giraffe for read mapping on large pathogen pangenomes and found graph-based mapping computationally challenging at this scale (Table S2). For example, after a 7h 39m indexing step, VG Giraffe took 26 minutes and 21 GB of memory to map 25 150bp reads onto a graph of 20,000 SARS-CoV-2 genomes using 25 threads. We also evaluated VG Giraffe’s haplotype sampling mode (Sirén et al. 2024), which improves mapping speed by restricting alignment to a set of 4 synthetic haplotypes (default setting) but still requires the 7h 39m indexing step. Using 1 thread, VG Giraffe completed haplotype sampling and alignment of 19,934 simulated SARS-CoV-2 reads (∼100× coverage) in 1m 27s, with peak memory usage of 470.9 MB. Using the 20,000-genome PanMAN, Panmap completed the same task in 1.42 seconds and 135 MB of memory, after a 3 second indexing step.

### Panmap assembles higher quality genomes than single-reference methods

Genome assembly with Panmap is more accurate than single-reference-guided methods for RSV, SARS-CoV-2, and *M. tuberculosis*, especially at very low depths of coverage (Figure 3). We evaluated Panmap’s accuracy using real sequencing reads from public SRA datasets with reference pangenomes containing 4,000 RSV, 20,000 SARS-CoV-2, and 400 *M. tuberculosis* genomes. For each sample, we performed leave-one-out evaluation: the sample’s known genome was removed from the pangenome before placement and assembly. Ground truth sequences were obtained directly from the PanMAN leaf nodes (the original GenBank submissions). We subsampled reads to coverages ranging from 0.5× to 100× and assembled each sample with Panmap and separately with a field standard single-reference consensus assembly workflow for each species: BWA-MEM + iVar consensus (Li 2013; Grubaugh et al. 2019) for RSV and SARS-CoV-2, and Clockwork (Hunt et al. 2022) for *M. tuberculosis.* We measured accuracy as the fraction of the ground truth genome (excluding 150 bp at each end) correctly reconstructed by each method. At 0.5× coverage, Panmap correctly genotyped a median of 91% of the RSV genome compared to 15% for BWA + iVar, which can only call bases at positions with read support. At 100× coverage, Panmap reaches 97% genotyping accuracy while BWA + iVar plateaus at 82%, because the standard RSV-A (NC_038235.1) and RSV-B (NC_001781.1) references are too divergent from contemporary circulating strains to achieve complete read mapping. For SARS-CoV-2, both methods converge near 100% at 100× coverage, but Panmap maintains >99% accuracy even at 0.5× depth where BWA + iVar covers only ∼30% of the genome. For M. tuberculosis, both Panmap and Clockwork achieve >99.5% median accuracy at all coverage levels, reflecting the species’ lower genetic diversity (more samples are similar to the standard reference genome). However, Panmap achieves slightly better accuracy (median >99.8%) in a fraction of the runtime. Panmap’s advantage stems from pangenome-guided placement: by selecting the closest reference from hundreds or thousands of genomes, reads map more completely, and uncovered positions inherit bases from a near-identical reference rather than being left as ambiguous.

**Figure 3.**
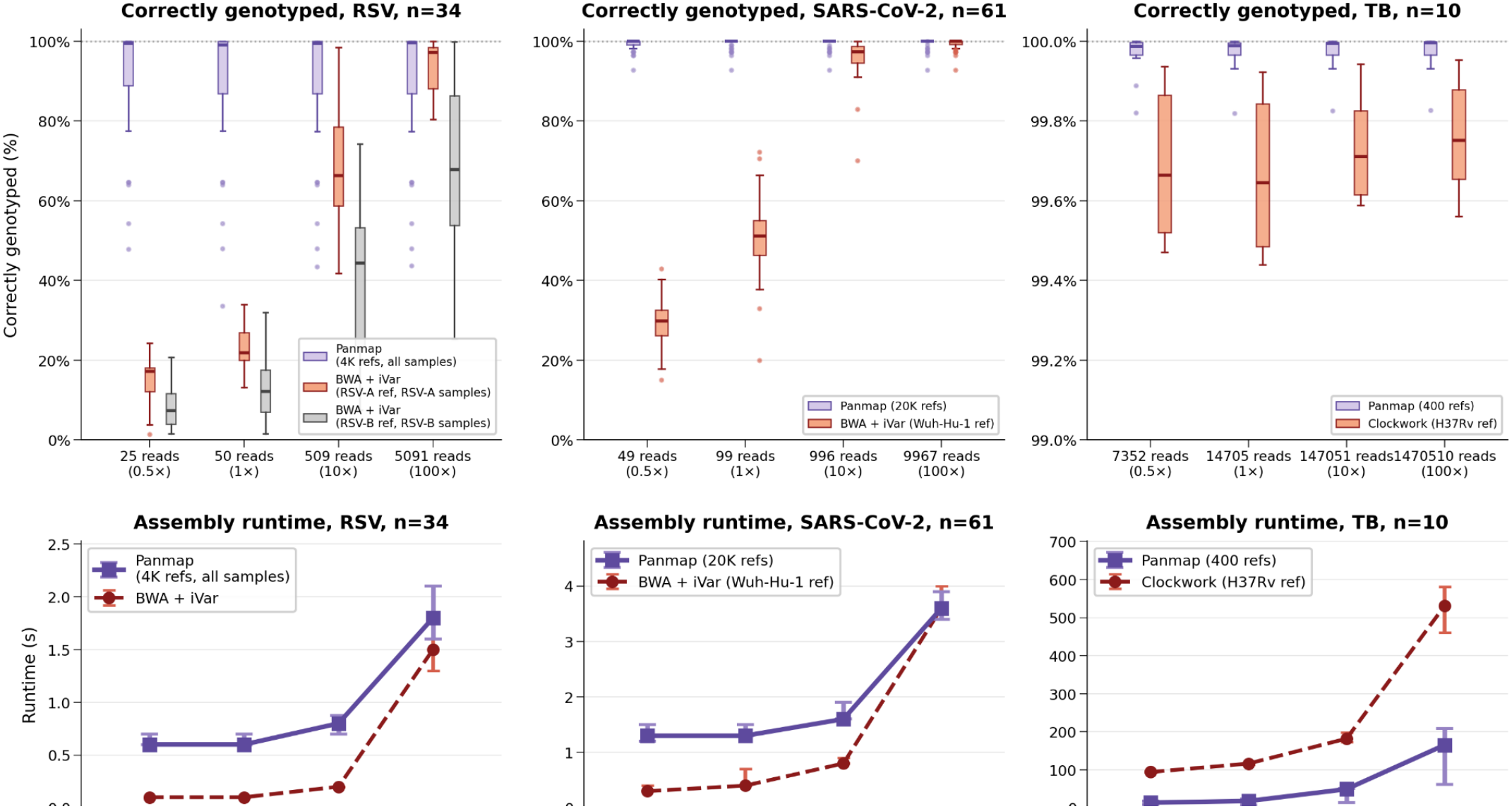
Consensus genome assembly accuracy and performance of Panmap compared to field-standard single-reference workflows on real sequencing data. (A) Consensus assembly accuracy: the percentage of the ground truth genome (excluding 150 bp at each end) correctly reconstructed by each method for SARS-CoV-2 (20,000-genome PanMAN vs. BWA+iVar with NC_045512.2), RSV (4,000-genome PanMAN vs. BWA+iVar with LR699737.1), and M. tuberculosis (400-genome PanMAN vs. Clockwork with NC_000962.3). We subsampled real sequencing reads to four coverage levels and compared each assembly to a high-coverage de novo ground-truth assembly (see Methods) using minimap2. Error rates are event-based: each SNP counts as one error, and each contiguous insertion or deletion counts as one error regardless of length. Box plots show the distribution of per-assembly error rates; lower is better. (B) Runtime comparison between Panmap and the reference-based pipeline for each species.

### Panmap resolves haplotype mixtures across diverse viral species

In metagenomic mode, Panmap accurately estimates haplotype abundances in simulated and real SARS-CoV-2 mixture samples, even when haplotypes carry variants absent from the reference tree. We evaluated metagenomic Panmap using the published SARS-CoV-2 PanMAN containing 20,000 SARS-CoV-2 sequences spanning 1,000 Pango lineages, as described in Walia et al. We simulated read mixtures with 1,000× coverage from up to 10 sampled haplotypes, each haplotype containing up to 20 introduced SNPs. We evaluated Panmap on the simulated samples and compared estimated haplotype abundances to the truth with a 3 bp difference tolerance (see methods) (Figure 4A). Abundance estimates closely matched their simulated proportions across all mutation levels, with root mean square error (RMSE) values of 0.009, 0.011, 0.011, 0.020, and 0.036 for samples containing 0, 1, 5, 10, and 20 SNPs per haplotype, respectively. As expected, increasing mutation levels led to more haplotypes not in the truth set to be estimated to be present (“others” category), with cumulative abundances of 0.27%, 0.60%, 0.95%, 2.03%, and 3.68% across the 40 replicates at each SNP level, reflecting the added noise introduced by unaccounted mutations. Despite this trend, false positives remained low in most scenarios, and true haplotypes were accurately estimated even in samples containing 200 total unaccounted mutations, implying that our method will be applicable for populations that are less densely sampled than major human pathogens. Simulations conducted on an RSV PanMAN with 4,000 samples and an HIV PanMAN with 20,000 samples yielded concordant results (Figure S4). For HIV, we demonstrated that our method can be applied to more divergent species that present challenges for single-reference based approaches, such as Freyja, WEPP, and ref. Pipes et al. (Karthikeyan et al. 2022; Pipes et al. 2022; Gangwar et al. 2025) We compared Panmap specifically to WEPP as it has been shown to be the most accurate and generalizable among single-reference based tools (Karthikeyan et al. 2022; Pipes et al. 2022; Gangwar et al. 2025). As WEPP requires an UShER-format mutation-annotated tree (MAT), we constructed an HIV MAT following the *viral_usher* pipeline (Hinrichs) using the same 20,000 samples as the published HIV PanMAN and using K03455.1 as the root. Our method achieved substantially lower weighted haplotype and peak distances than WEPP (Figure S5). While WEPP relies on a single root reference and is susceptible to information loss specifically through alignment failure of highly divergent reads, our approach utilizes the entire pangenomic reference.

**Figure 4.**
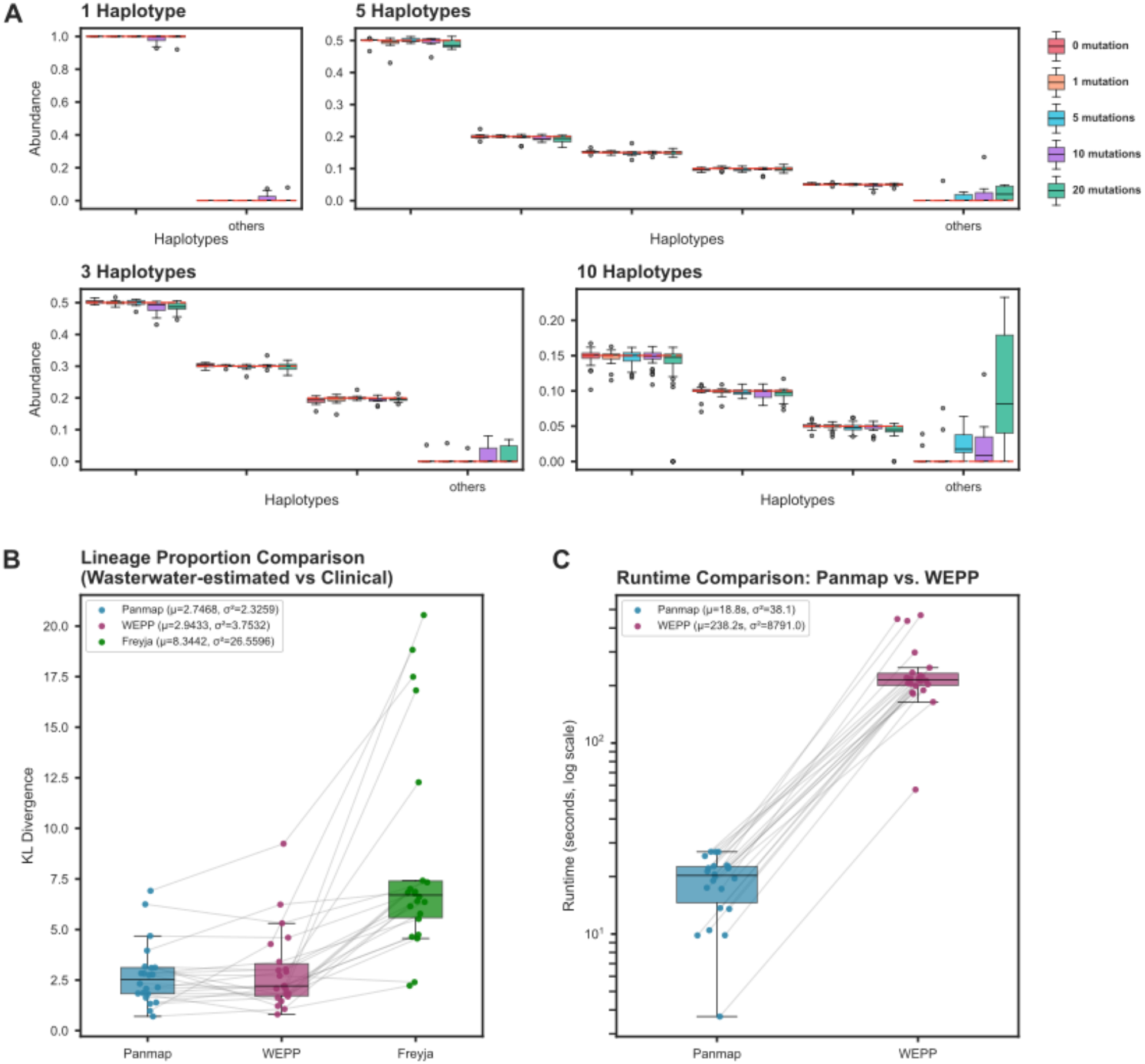
Evaluation of abundance estimation in simulated and wastewater SARS-CoV-2 mixture samples. (**A**) Estimated haplotype abundance in simulated SARS-CoV-2 mixture samples with 1, 3, 5, or 10 haplotypes, introducing 0, 1, 5, 10, or 20 single-nucleotide polymorphisms (SNPs) per haplotype. Ten replicates were simulated per composition and mutation level. Red dashed lines denote true simulated abundances; the ’others’ category represents accumulated false-positive haplotype abundance. (*10-haplotype samples: 4 haplotypes at 15%, 2 at 10%, 4 at 5% abundance) **(B)** Kullback-Leibler (KL) divergences between lineage proportions estimated by Panmap, WEPP, and Freyja from 22 wastewater samples, compared to one-week-delayed clinically observed lineage proportions. **(C)** Runtime comparison between Panmap and WEPP in seconds. Gray lines connect results from the same wastewater sample

### Panmap outperforms existing tools on real wastewater surveillance data

We next evaluated metagenomic Panmap’s performance on 22 real SARS-CoV-2 wastewater samples collected at Point Loma between November 27, 2021, and February 7, 2022 (Karthikeyan et al. 2022). We evaluated our estimated lineage proportions by comparing them to those derived from clinical sequences in San Diego with a 7-day temporal shift as a proxy for ground truth (following Karthikeyan *et al*. and Gangwar *et al*.). This window was selected for its high epidemic surveillance efforts and high sequence submission rates. We used the same published SARS-CoV-2 PanMAN described above for Panmap, and a subsampled SARS-CoV-2 UShER-format MAT containing the same 20,000 sequences as the PanMAN for WEPP. Compared with WEPP and Freyja, Panmap achieved higher accuracy than Freyja and comparable accuracy to WEPP (Figures 4B, S6, S7), while demonstrating a >10X speedup in runtime (Figure 4C).

### Application to ancient DNA phylogenetic analysis

Panmap’s metagenomic mode demonstrates utility for ancient environmental DNA (eDNA) analysis. We constructed a vertebrate mitochondrial genome pangenome containing 15,655 sequences representing 7,893 unique species and indexed it with Panmap (see Methods). To assign eDNA reads to specific taxa, we implemented a competitive mapping approach in Panmap that filters low-confidence mappings, uneven coverage, and phylogenetically ambiguous reads matching multiple distantly related taxa (see Methods). This method is similar to LCA-based approaches commonly used in environmental and ancient DNA studies that map reads against large multi-terabyte reference databases (Wang et al. 2021; Kjær et al. 2022; Wang et al. 2022; De Sanctis et al. 2025).

With these implementations, Panmap filters and assigns reads and read clusters concordant with published results. We applied this approach to two recently published ancient sedimentary DNA datasets: circumpolar ancient permafrost DNA (Wang et al. 2021; Miller and Simpson 2022) and 2-million-year-old ancient sediment DNA (Kjær et al. 2022). Although our phylogenetic placement evaluated all vertebrates, we focused on the Elephantidae family for both studies, as genera within this family were found in both previous studies, and are of high ecological interest (Wang et al. 2021; Miller and Simpson 2022).

For the Wang et al. (2021) data, we obtained adapter-trimmed reads and placement results directly from the authors and performed quality control on the FASTQ files before applying our method (see methods). In accordance with the original paper’s findings, we identified a clear read assignment hotspot in the Elephantidae clade, specifically at the *Mammuthus* nodes (shown across all Wang et al 2021 samples in Figure 5A). Panmap processed ∼9 billion reads in the original 651 samples in 8 minutes using 16 threads, and assigned a total of 4,098 reads to nodes within the Elephantidae family, more reads than we assigned to any other families. This is nearly five times more than the 846 total mammoth reads identified by the traditional competitive mapping approach which was implemented in Wang et al (2021). There are several reasons for this, but two important ones are that our approach does not explicitly require reads to be aligned, or indeed be alignable, to reference genomes and that only nodes with maximal placement score to a read are considered for the taxa LCA filter.

**Figure 5.**
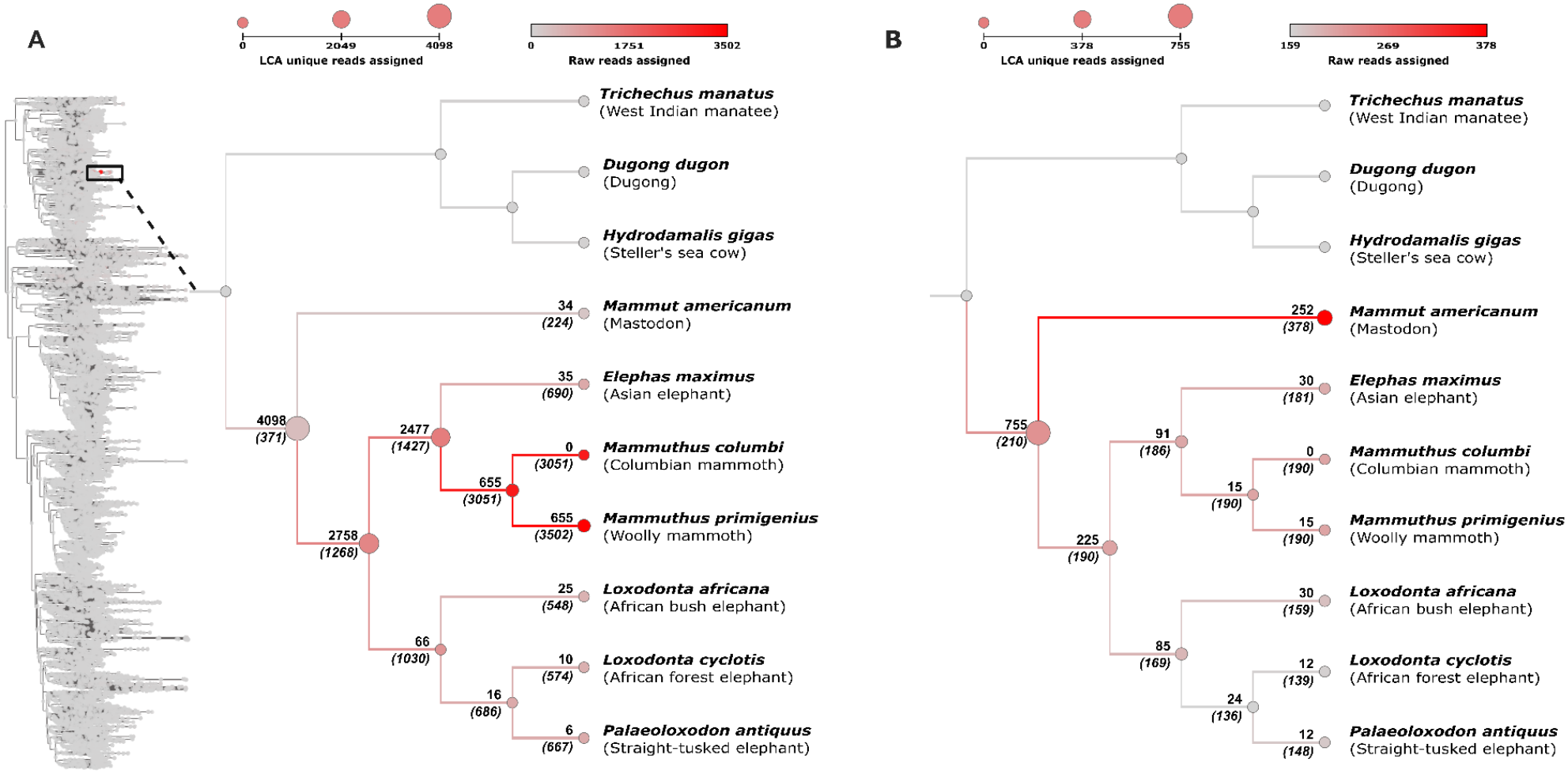
Number of reads assigned to nodes for two datasets processed by Panmap. Note that reads may be assigned to more than one node within a family. Numbers above branches indicate LCA counts, or the total number of reads assigned *only* to that node or any node below it, i.e. not including any reads which are also assigned elsewhere. Numbers in parentheses below branches indicate the count of raw reads assigned directly to the node (which may also be assigned elsewhere). Node color scales with raw read count, and node size scales with LCA read count. **(A)** Read assignments for the Wang et al. dataset. The full vertebrate mitochondrial PanMAN is shown on the left, with nodes colored by raw read assignment count; the highlighted Elephantidae clade is expanded on the right. **(B)** Read assignments for the Kjær et al. dataset, restricted to the Elephantidae clade.

These Panmap-assigned reads were further validated to be *Elephantidae* using full-read alignment when possible, BLAST (Altschul et al. 1990), average sequence divergence, and postmortem damage plots (see methods, Figure S8, Table S6). ∼93% of the assigned reads were alignable to the *Mammuthus primigenius* mitochondrial reference genome and ∼93% of the alignable reads had exclusive BLAST hits to Elephantidae mitochondria. Finally, out of the 4,098 Elephantidae reads identified from our vertebrate-wide PanMAN, we placed the 3,803 reads that are alignable to the *M. primigenius* mitochondrial genome to a *Mammuthus*-specific PanMAN containing 78 Mammuthus mitochondrial genomes, for each Wang et al 2021 sample separately (see Methods). This allowed direct comparison to the original per-sample pathPhynder (Martiniano et al. 2022) placement results. Panmap achieved on average more than 2 branches deeper placement than pathPhynder, yielding more informative placements overall (Figure S9, Table S5).

For the 2-million-year-old sample from Kjaer et al., we downloaded the published raw data, performed quality control, and applied our method as above by placing reads from all samples combined into our vertebrate PanMAN. Our placement shows a concordant cluster of reads assigned to the mastodon node within the Elephantidae and Mammutidae families (Figure 5B). The original Kjaer et al. study noted that the mastodon placement was supported by only two single-nucleotide polymorphisms (SNPs), and this was later confirmed in further work (Vogel et al. 2024). Our analysis provides further evidence supporting this placement.

## Discussion

Panmap addresses a central limitation of existing pangenome tools by enabling phylogenetic placement, genotyping, and assembly against reference pangenomes containing tens of thousands of genomes, a scale at which graph-based and alignment-based approaches become impractical. By incorporating evolutionary information, Panmap improves the resolution and efficiency of studying new samples, achieving better accuracy than single reference-guided genotyping methods for *M. tuberculosis*, respiratory syncytial virus (RSV), and SARS-CoV-2. The accuracy gap widens at low coverage, where a poorly chosen reference introduces systematic errors that sparse read data cannot correct. We expect this advantage to grow as pangenomes continue to become larger and more densely sampled. Panmap can also place and align reads to inferred ancestral sequences at internal nodes, providing suitable references even for samples that diverge from all observed genomes in the pangenome.

Panmap consistently produced smaller indexes than similar methods that index pangenomes. Panmap’s indexes for the SARS-CoV-2 and RSV pangenomes were hundreds or thousands of times smaller than VG Giraffe’s graph-based indexes. However, Panmap’s index was only about 30% smaller than VG Giraffe’s for *Mycobacterium tuberculosis*. One reason for this is that the pangenome has fewer, though larger, genomes, making Panmap’s compression advantage less apparent than in the viral pangenomes comprising thousands of samples. Another reason is that the pangenome has a more complex (Table S4) or simply longer evolutionary history, reducing its phylogenetic compressibility. The compression advantage of Panmap’s *k*-mer index is likely greatest when the overall sequence diversity is minimized and the phylogenetic relatedness is maximized in the reference pangenome, and more evident as the number of individual genomes increases.

Panmap’s metagenomic mode broadens its utility to mixed-sample haplotype deconvolution from bulk sequencing data, addressing a long-standing bottleneck in population and microbial genomics.. In simulated SARS-CoV-2 mixtures, abundance estimates remained accurate even with constituent haplotypes carrying up to 20 unobserved single-nucleotide polymorphisms (SNPs), demonstrating robustness against incomplete reference pangenome representation. This robustness is crucial for surveillance of under-sampled pathogen populations, where reference phylogenies often lag behind circulating diversity. Concordant results across RSV and HIV pangenomes further support this generalizability, particularly for HIV where Panmap handles the elevated sequence divergence that causes single reference-based methods, such as WEPP, to lose accuracy. On real wastewater data, Panmap achieved KL divergence comparable to WEPP in estimating clinically derived lineage proportions (Karthikeyan et al. 2022; Gangwar et al. 2025), while running 10X faster than WEPP. This favorable accuracy-speed trade-off supports its utility for high-throughput wastewater-based epidemiology.

The ancient eDNA application demonstrates that Panmap’s metagenomic framework generalizes beyond within-species haplotype deconvolution to cross-species taxonomic assignment. By competitively mapping reads against a vertebrate mitochondrial pangenome of nearly 16,000 sequences, Panmap recovered substantially more high-confidence Elephantidae reads than the commonly-used two-step ancient environmental DNA competitive mapping and phylogenetic placement approach, while producing placements concordant with published findings (Wang et al. 2021; Kjær et al. 2022). The ability to score and place reads in a single framework, without requiring a separate alignment step or a multi-terabyte reference database, suggests that pangenome-based competitive mapping may offer a practical alternative to LCA-based workflows for degraded and low-complexity ancient DNA.

Panmap’s phylogenetic approach to pangenomic analysis offers advantages over both single-reference and graph-based methods: it scales to millions of genomes, produces highly compressed indexes, and leverages evolutionary relationships to improve both placement and genotyping accuracy. For metagenomic applications, the same phylogenetic framework enables haplotype deconvolution and abundance estimation with reduced reference size and computational cost. These capabilities make Panmap particularly well-suited for real-time pathogen surveillance, where rapid turnaround and scalability are essential, as well as for retrospective analyses of ancient and environmental DNA where sample quality and reference completeness are limiting factors.

## Methods

Panmap accepts two inputs: a reference pangenome in PanMAN format (Walia et al. 2026) and sequencing reads in FASTQ format. The PanMAN encodes both a phylogeny and genomic variation, including point mutations, insertions, deletions, and structural variants. PanMANs may be constructed from multiple sequence alignments using panmanUtils (https://turakhia.ucsd.edu/panman). Panmap supports single or paired-end reads, and is optimized for short reads.

Panmap has two modes. Single-sample mode assumes all reads originate from one haplotype; it jointly scores reads to identify an optimal reference, aligns reads to that reference, and produces a genotyped consensus assembly. Metagenomic mode handles mixed samples containing multiple haplotypes; it scores each read independently against every node in the phylogeny and then either infers the constituent haplotypes and their relative abundances from the score distribution, or applies competitive mapping filters for eDNA read assignment. The following sections describe components shared by both modes, followed by mode-specific methods and experiment details.

### Reference PanMAN structure and genome decoding

A PanMAN pangenome comprises an ancestral consensus sequence, a phylogeny, and mutations annotated along branches of the tree. Each node in the phylogeny represents a full genome or haplotype: observed sequences at leaf nodes and inferred ancestral sequences at internal nodes. Any genome in the pangenome can be recovered by tracing its lineage from the root and applying the branch-labeled mutations at each step. Panmap leverages this structure by performing a single depth-first traversal of the phylogeny, maintaining one genome sequence in memory that is incrementally mutated at each node. This traversal-based approach enables rapid indexing of pangenomes containing many thousands of sequences.

### Indexing k-mer seeds

We compactly index the positions and hashed sequences of all seeds (linked syncmers) in the reference pangenome by compressing them according to phylogenetic structure. Because similar sequences are expected to share many seeds, we use the phylogenetic tree to annotate seeds only at the node where they first appear. To build our seed index, we traverse the tree once in depth-first order, maintaining a list of seed starting positions that is updated at each node’s haplotype. The index is stored on disk as a depth-first list of nodes and a list of “mutations” for each node that encodes insertions or deletions of hashed seeds relative to the parent node’s seed list. For metagenomic mode, the additionally stores seed orientations, which adds a small overhead (Table S1).

### Indexing optimizations

We employ several additional strategies to make seed indexing faster and compact. During traversal, we expect each taxon to have nearly the same list of seeds as its parent, except for a small number of changes near mutations to the nucleotide sequence. Thus, we only recompute seeds within *k* bases (the length of a syncmer) upstream of each nucleotide mutation at a given taxon, retaining the rest of the seeds from the parent. The indexing algorithm is multithreaded by partitioning the depth-first traversal into equal segments that are processed independently and then merged. To write compressed index data to disk and decode it during placement, we use Cap’n Proto (https://github.com/capnproto/capnproto) to serialize the index followed by a Zstandard (https://github.com/facebook/zstd) compression layer (compression level 7 by default).

### Seed sketches of genome sequences

For sample placement and read alignment, we consider a subset of *k*-mers, called seeds, from each genome or haplotype in a PanMAN. Retained seeds for each sequence are chosen under a syncmer scheme (Edgar 2021) so that the similarity between haplotypes can be approximated by the similarity of their seed sketches. Syncmer seeds are parameterized by two values, k and s, selecting as a seed any k-mer whose lexicographically smallest s-mer begins at either the start or end of the k-mer. In practice, we order k-mers and s-mers by a hashed integer value rather than their lexicographic ordering. User-supplied values of k and s, or default values k=19 and s=8, determine the syncmer parameterization and may be tuned to affect the density and length of sampled seeds. Following the minimizer-based approach of (Ekim et al. 2023) we also define *l-*linked syncmers: ordered tuples of *l* consecutive syncmers treated as a single matching unit. Linked syncmers represent longer genomic regions than individual syncmers while remaining robust to local variation. Panmap uses 3-linked syncmers (l=3) by default.

### Mutation spectrum prior for genotype likelihoods

We compute a mutation spectrum prior for a reference pangenome by performing a depth-first traversal of its phylogeny, improving genotyping accuracy by incorporating evolutionary context. During traversal, we count substitutions between each parent and child node, recording their mutation types (e.g., A→C, G→T) in a 4×4 substitution matrix. Substitutions are counted only in aligned regions present in both parent and child. After traversal, we normalize each count by the total observations of each reference base to obtain substitution probabilities. The mutation spectrum is scaled to an expected branch length (user-specified, or 0.0011 by default, matching minimap2’s mutation rate).

### Genotype likelihoods and consensus assembly

Panmap computes initial genotype likelihoods internally using bcftools mpileup and bcftools call (--ploidy 1 -cA), then applies the mutation spectrum prior for nucleotide substitutions. The posterior probability of genotype g given data D is:

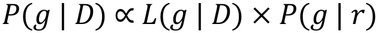

where *L*(*g* | *D*) is the genotype likelihood from bcftools and *P*(*g* | *r*) is the prior probability that reference base r mutates to variant base g, derived from the mutation spectrum. For indels, we use unmodified posterior probabilities from bcftools. The consensus sequence is called from the final posterior genotype probabilities.

### Phylogenetic placement in Single-sample mode

In single-sample mode, Panmap identifies an optimal reference haplotype by collectively comparing input reads to every sequence in the pangenome. Rather than scoring each read individually (as in metagenomic mode), single-sample Panmap collects k-mer seeds from all reads and compares them to the indexed seeds of each node. A single depth-first traversal of the phylogeny assigns each node a similarity score approximating its sequence identity to the input sample. During traversal, we maintain a running list of seeds at the current node by applying indexed changes to the parent’s seed list, avoiding recomputation. The highest-scoring node, whether a sampled genome at a leaf or an inferred ancestral haplotype at an internal node, is selected as the reference for downstream alignment and genotyping.

Our default similarity metric is a log-scaled containment index. For each node i, we compute:

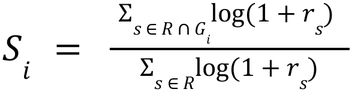

where *R* is the set of unique seeds extracted from all reads, *G_i_* is the set of seeds present in genome *i*, and *r_s_* is the frequency of seed *s* across all reads. This metric measures the fraction of read seed content contained in each genome, with log-scaling so that high-coverage seeds do not dominate the score.

We compute this metric dynamically across all nodes by incrementally updating the numerator using only the seed changes between parent and child nodes stored in the index. Panmap can alternately use several other scoring metrics, including log-scaled cosine similarity and raw seed matches, though containment performed best in our experiments (Figure S10).

### Read alignment in Single-sample mode

After a suitable reference is chosen, in single-sample mode all input reads are aligned to this sequence using a seed-chain-align approach. Alignment seeds are by default recomputed with smaller *k*-mer size, or optionally, seeds identified during placement may be passed directly to the chaining and alignment functions of the Minimap2 library (Li et al. 2018). Panmap also supports a bwa-aln (Li and Durbin 2009) mode that improves mapping sensitivity for very short or damaged reads, such as those from ancient DNA samples (Figure 3). Because bwa-aln outperforms other alignment algorithms for ancient DNA (Oliva et al. 2021), we recommend this mode for such samples.

### Read scoring in Metagenomic mode

For mixed samples containing multiple haplotypes, Panmap’s metagenomic mode scores each read independently against every node in the phylogeny, rather than jointly scoring all reads. Read scores can be either the raw count of seed hits or a pseudo-chaining score (raw count by default). From the query reads, Panmap generates a seed-to-read hash map, where each key is a seed hash and each value is the list of read indices containing that seed. This enables rapid retrieval and updating of reads affected by a seed change during tree traversal. A child node effectively inherits all read scores unaffected by seed changes from its parent.

We implemented several strategies to further optimize computational efficiency during read scoring:

**Read score annotations:** To avoid storing read scores for every read against every node, Panmap only records read scores when they are updated at a node. The complete read scores for any node can be reconstructed by traversing from the root to that node and sequentially applying the recorded score changes.

**Query de-duplication:** Sequencing datasets often contain many identical reads, particularly for amplicon protocols. Panmap collapses exact duplicate reads into unique sequences with associated counts before further processing. Reads are then sketched into seeds, and reads sharing identical seed sketches are further collapsed. This two-stage deduplication substantially reduces the number of reads to be placed and scored.

**Seed pruning and tree condensation:** Only seeds that are present in both the query and the reference tree contribute to read scoring; therefore, query-only or reference-only seeds can be discarded with no impact. Panmap discards query-only seeds from the query and prunes seed changes of reference-only seeds from the tree. After reference-only seeds are pruned, Panmap further condenses the tree by merging nodes that have no seed mutations post-pruning. This greatly reduces wasteful iterations and lookups, as well as memory required for seed storage.

### Abundance estimation in Metagenomic mode

We implemented an expectation–maximization (EM) algorithm accelerated by SQUAREM (Du and Varadhan 2020) internally to estimate the haplotypes present and their proportions (following Pipes et al. 2022). To reduce the search space for the EM algorithm, we select a subset of the N most probable haplotypes (default N=1000) by identifying the nodes with the highest seed overlap coefficients (OC) with the query, where the overlap coefficient between a reference R and a query Q is:

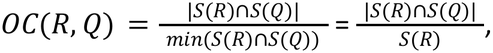

where S(R) is the set of unique seeds sketched from the reference, and S(Q) is the set of unique seeds sketched from all query reads in the sample. A probability matrix of size (Q, N), where p_qn_ is the probability of query q being generated from the genome at node n, is estimated by

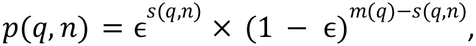

where ɛ is the error rate (default: 0.005), *s*(*q*, *n*) is the score of query q against node n, and *m*(*q*) is the total number of seeds sketched from query q (i.e. the maximum possible score query q can have).

An abundance vector of size N is initialized with uniform proportions for probable haplotypes. After a round of EM converges, haplotypes with an estimated proportion less than a threshold (default to 0.5%) are discarded. An abundance vector for the remaining haplotype is initialized with uniform proportions, and another round of EM is performed. This repeats until all haplotypes are above the threshold proportion after an EM convergence.

### Experimental setup and data preparation

To evaluate the performance and accuracy of Panmap, we conducted a series of experiments using both real and simulated genomic datasets. We used reference PanMANs of *M. tuberculosis,* SARS-CoV-2, HIV, and RSV genomes, as well as a vertebrate and *Mammuthus* mitochondrial genomes. We evaluated Panmap on the pathogen pangenomes with real and simulated sequencing data, and used real ancient DNA reads for the *Elephantidae* pangenome.

**Virus and bacteria genomes**. For *M. tuberculosis,* SARS-CoV-2, HIV, and RSV, we downloaded reference pangenomes in PanMAN format from https://zenodo.org/records/17781629, comprising 400, 20K, 20K, and 4K genomes, respectively. We also downloaded the 8M-genome SARS-CoV-2 PanMAN from the same archive.

**Vertebrate mitochondrial genomes.** We obtained all vertebrate mitochondrial genomes from RefSeq. We generated the MSA and phylogenetic tree of these genomes using *TWILIGHT* and *DIPPER*, respectively, which were then passed on to *panmanUtils* to generate the PanMAN.

**Mammuthus mitochondrial genomes**. We obtained 78 *Mammuthus* spp. mitochondrial genomes and their inferred phylogeny from ref. (Martiniano et al. 2022). We then aligned the 78 sequences using TWILIGHT, and used the resulting multiple sequence alignment, together with the Martiniano et al. Newick tree as a guide, to construct a PanMAN.

### Panmap index evaluation

We evaluated Panmap’s index in comparison to VG Giraffe and EPIK, two other tools that index *k-*mers in pangenomes. EPIK requires a database of phylo-*k*-mers (Nikolai Romashchenko et al. 2023) built with the related tool IPK from a phylogenetic tree and a multiple sequence alignment. For read alignment to a pangenome graph in GFA format, VG Giraffe indexes *k-*mer minimizers, a graph Burrows-Wheeler transform, and node distances (Sirén et al. 2021; Sirén and Paten 2022). We provided each tool one CPU thread and measured the runtime, memory usage, and total size after GZIP of all index files needed to run the respective tool. All experiments were performed on a reserved node in a computing cluster, and we determined runtime and memory usage from the “elapsed (wall clock) time” and “maximum resident set size” reported by GNU *time* with the argument “-v”.

To build indexes with VG Giraffe, we first extracted Graphical Fragment Assembly (GFA) files from each reference PanMAN pangenome, then indexed the graph with *vg autoindex --workflow giraffe*, which produces a GBZ file (indexed graph Burrows-Wheeler transform), a *.dist* file (node distances in the graph), and a *.min* file (indexed *k*-mer minimizers).

We used IPK to build indexes for EPIK, beginning with a FASTA format multiple genome alignment and a phylogenetic tree, both extracted from the reference PanMAN pangenomes. We first optimized branch lengths of the starting phylogeny by running IQ-TREE with the “-te” argument, which performs maximum-likelihood optimization on a fixed tree topology. We used the GTR+G model for branch optimization, then supplied the best model parameters estimated by IQ-TREE to IPK with the “--ar-config” option for each species. We ran IPK with “-m GTR”, “-k 8”, and “--use-unrooted”, choosing a *k*-mer size of 8 based on the IPK documentation’s recommendation for large trees. IPK can use either RAxML-ng or PhyML for ancestral state reconstruction, and we first tried the default RAxML-ng mode for all species. For 20,000 SARS-CoV-2 genomes, IPK in RAxML-ng mode failed due to an error in branch length optimization, so we tried again with a different tree containing no branch lengths, which was unsuccessful for the same reason. For 400 *M. tuberculosis* genomes, IPK failed in RaxML-ng mode due to exceeding 1 TB RAM usage. We tried again with the PhyML mode, which supports datasets up to 400 sequences, though this run did not complete within 3 days.

### Phylogenetic placement evaluation, Single-sample mode

We evaluated Panmap’s phylogenetic placement on both simulated and real sequencing data. For simulations, we selected 70 leaf nodes at random from each reference PanMAN, applied random SNPs and indels (indel fraction 0.1) at two per-site mutation rates (μ = 10⁻⁴ and 10⁻³ for RSV and SARS-CoV-2; μ = 10⁻⁵ and 10⁻⁴ for M. tuberculosis) to generate novel descendant genomes, then simulated 150 bp paired-end reads at four estimated coverage levels (0.5×, 1×, 10×, and 100×), yielding 560 placements per species. For real data, we retrieved assembled genomes from GenBank and corresponding raw reads from the Sequence Read Archive, filtering for an estimated coverage depth of at least 500× and no more than five N characters in the assembly for the viral species; for M. tuberculosis, we selected complete genome assemblies with paired-end Illumina data available in SRA. We aligned each sample’s reads back to its assembly and discarded samples with more than 5 variants identified by bcftools call. The final real dataset comprised 51 SARS-CoV-2, 25 RSV, and 10 M. tuberculosis samples (Table S3).

For simulated samples, we placed reads directly onto the unmodified PanMAN and compared Panmap’s placement to the true parent of the simulated descendant. For real samples, we used a leave-one-out approach: we temporarily removed the sample’s leaf from the PanMAN, subsampled its reads to the target coverage depth, and placed them with Panmap. In both cases, we quantified placement accuracy as the difference in parsimony score between Panmap’s placed node and the sample’s expected position (placed minus original), adjusted to max(0, score) so that zero indicates an equivalent or better placement. Parsimony costs were computed by aligning the sample’s reference assembly to each candidate node with minimap2 and counting SNPs and indels.

### Haplotype abundance evaluation with simulated data

To evaluate Panmap’s ability to estimate haplotype proportions in simulated mixture samples, we randomly selected sets of 1, 3, 5, and 10 haplotypes from a reference PanMAN and simulated read mixtures for each haplotype level. To assess the effect of variants unrecorded in the tree, we also introduced 0, 1, 5, 10, and 20 single-nucleotide polymorphisms (SNPs) to each haplotype in a sample, generating 10 replicates for each combination of haplotype number and mutation level (200 total replicates). 150-bp paired-end reads at 1000× average depth were simulated using *InSilicoSeq* and its pre-built NovaSeq error model. Given our objective is estimating major cluster abundance rather than exact sequences, we defined “acceptable resolution” for lineage tracking as haplotypes within <=3 bp of the true sequence.

This evaluation was performed for SARS-CoV-2, RSV, and HIV, using panMANs published by Walia et al. (https://zenodo.org/records/17781629).

### Genome assembly evaluation with real data

We evaluated consensus genome assembly using Panmap and commonly used reference-based pipelines on real sequencing data. For SARS-CoV-2 and RSV, we compared against BWA-MEM + iVar consensus, using the NC_045512.2 reference for SARS-CoV-2, the RSV-A reference (NC_038235.1) for RSV-A samples, or the RSV-B reference (NC_001781.1) for RSV-B samples (determined by competitive mapping of the sample’s reads to each reference before evaluation). For M. tuberculosis, we compared against Clockwork variant_call_one_sample(Trimmomatic → minimap2 → Cortex + bcftools → Minos adjudication; NC_000962.3 reference). Each method produced a consensus FASTA, which we compared to the ground truth consensus FASTA. We performed quality control filtering and trimming on all reads with fastp prior to evaluation, and trimmed primers for SARS-CoV-2 amplicon protocols with iVar.

We constructed ground truth genomes for 25 RSV genomes, 51 SARS-CoV-2 genomes, and 10 M. tuberculosis genomes. Because assembly quality of viral data is variable, for SARS-CoV-2 and RSV we assembled ground truths from the full QC’d read sets (high coverage, >500×) using the Broad Institute assemble_denovo pipeline (SPAdes de novo assembly → scaffold against reference → read-back refinement). For M. tuberculosis, we used sequences stored in the PanMAN directly (losslessly identical to a published Genbank assembly). We subsampled each dataset to ∼0.5×, 1×, 10×, and 100× estimated coverage based on genome length. For each assembled consensus by Panmap, BWA+iVar, and Clockwork, we compared each assembly to its ground truth using minimap2 asm20 alignment (--cs), quantifying SNP and indel errors and genome coverage. The final real dataset comprised 74 SARS-CoV-2, 25 RSV, and 10 M. tuberculosis samples.

### SARS-CoV-2 lineage abundance estimation from wastewater data

We evaluated Panmap’s capability to estimate lineage abundance using 22 SARS-CoV-2 wastewater samples, collected by Karthikeyan et al. at Point Loma (San Diego, California) between November 27, 2021 and February 7, 2022. We chose this period due to high local epidemic surveillance and sequencing submission activity. For Panmap, we used a SARS-CoV-2 PanMAN comprising 20,000 samples and 1,000 Pango lineages (https://zenodo.org/records/17781629), with the most recent sample submitted on January 5, 2024. For comparison with WEPP, we used the next available UShER-format Mutation-Annotated Tree (MAT) on the UCSC Genome Browser (Casper et al. 2026) after this date (<u>public-2024-02-01.all.masked.pb.gz</u>).

We preprocessed the wastewater reads by aligning to the Wuhan-1 SARS-CoV-2 reference (NC_045512.2) using minimap2 (minimap -a --sam-hit-only –MD -2 -x sr) (Li et al. 2018; Li 2021) and trimming the primer sequences using ivar (ivar trim -e -q 1 -m 80 -x 3) (Grubaugh et al.

2019).

For Panmap, we converted the trimmed bam files to FASTQ format and removed trimmed regions using samtools. As the wastewater data were generated using tiled amplicons, metagenomic Panmap removes reads whose seed relative frequencies are less than 0.01 compared to the total number of reads associated with the same amplicon to mitigate spurious PCR and sequencing errors.

We ran WEPP and Freyja with the default settings and commands as described in their documentation, following the same mapping and trimming procedures described above.

### Ancient environmental DNA placement

We obtained the adaptor-trimmed circumpolar ancient permafrost DNA dataset from ref. (Wang et al. 2021) and preprocessed the reads using *fastp* (Chen 2023) and *sga (Simpson and Durbin 2012)* to deduplicate exact substrings, trim poly Gs, and remove reads with low quality, low complexity, and short length. We used the following specific commands and parameters:

`fastp -V -D -A --dup_calc_accuracy 5 -g -x --poly_g_min_len 5 --poly_x_min_len`

`5 -q 30 -e 25 -l 30 -y -p -w 8`

`sga preprocess --dust-threshold=30 -m`

`sga index --algorithm=ropebwt; sga filter --no-kmer-check`

We implemented several competitive mapping-based filters for eDNA taxonomic assignment. Let *r* denote a sequencing read and *S*(*r*) the set of unique seeds sketched from *r* , with |*S*(*r*)| denoting the total seed count.

**Low seed-hit filter.** For each read *r*, let σ(*r*) denote its maximum alignment score against the reference tree. The seed-hit ratio is defined as:

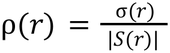

Reads for which ρ(*r*) < τ*_m_* (default τ*_m_* = 0.7) are discarded as low mappability.

**Phylogenetic ambiguity filter.** Let *T*(*r*, *l*) denote the set of taxa to which read *r* is assigned at taxonomic rank *l*. A read is discarded as phylogenetically ambiguous if:

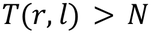

where *N* is a user-defined threshold (default *N*=1) and *l* defaults to the family rank.

**Evenness of coverage filter.** For a reference node *v*, let *R*(*v*) denote the set of reads assigned to *v*, and let *S*(*v*) be the set of seeds sketched from the reference sequence of *v*, with |*S*(*v*)| the total reference seed count. Define ℎ(*s*) as the number of hits to seed *s* ∈ *S*(*v*) from reads in *R*(*v*), and let *D*(*v*) denote the mean number of hits per seed:

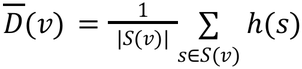

The observed seed coverage is:

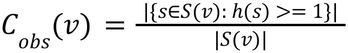

Under a Poisson model with rate *D*(*v*), the expected seed coverage is:

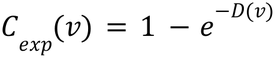

Node *v* is discarded if the coverage ratio 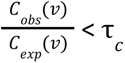, where τ*_c_* defaults to 0.2. Reads assigned to any discarded node are subsequently removed as well.

### Panmap-assigned reads validation from Wang et al. dataset

We validated Panmap-assigned reads by first aligning them to the *Mammuthus primigenius* mitochondrial reference genome (GenBank: DQ188829.2) on the PanMAN using bwa-aln (Li and Durbin 2009) with parameters tuned for ancient DNA (bwa-aln -t 8 -l 1024 -q 15 -n 0.02) <u>(Martiniano et al. 2022)</u>. We then performed nucleotide BLAST (Altschul et al. 1990) on all the alignable reads using the core_nt database (blastn -evalue 0.05 -max_target_seqs 1000). For a sequence, BLAST hits were retained only if their bit scores fell within the top 10% of the maximum score observed for the corresponding sequence, following Slon et al. (Slon et al. 2017). Reads were considered to have exclusively Elephantidae mitochondrial hits if their retained matches were only hits to mitochondrial genomes of Elephantidae species.

### Per-sample placement on Mammuthus PanMAN

We placed each sample of Panmap-assigned reads from the Wang et al. data onto a *Mammuthus* species PanMAN. Samples were processed using haplotype deconvolution as described above. Although most samples returned a single ambiguous haplotype group (i.e. haplotypes indistinguishable from the given data), for samples estimated to contain more than one haplotype group, the most abundant group was selected. Each sample is then placed at the last common ancestor of all haplotypes within the selected group.

## Acknowledgements

We thank Pranav Gangwar for guidance on wastewater surveillance analyses and Yucheng Wang for providing circumpolar Arctic ancient DNA data and placement results. We also thank Anthony Hu and Cassidy Sullivan for their helpful comments on the manuscript.

The authors acknowledge funding from R35GM128932 (to RCD) and the Presidents postdoctoral fellowship program (PPFP to BDS).

