## Supplementary Information for "Panmap: Scalable phylogeny-guided alignment, genotyping, and placement on pangenomes"

**Supplementary Figures and Tables:**

**Figure S1:**
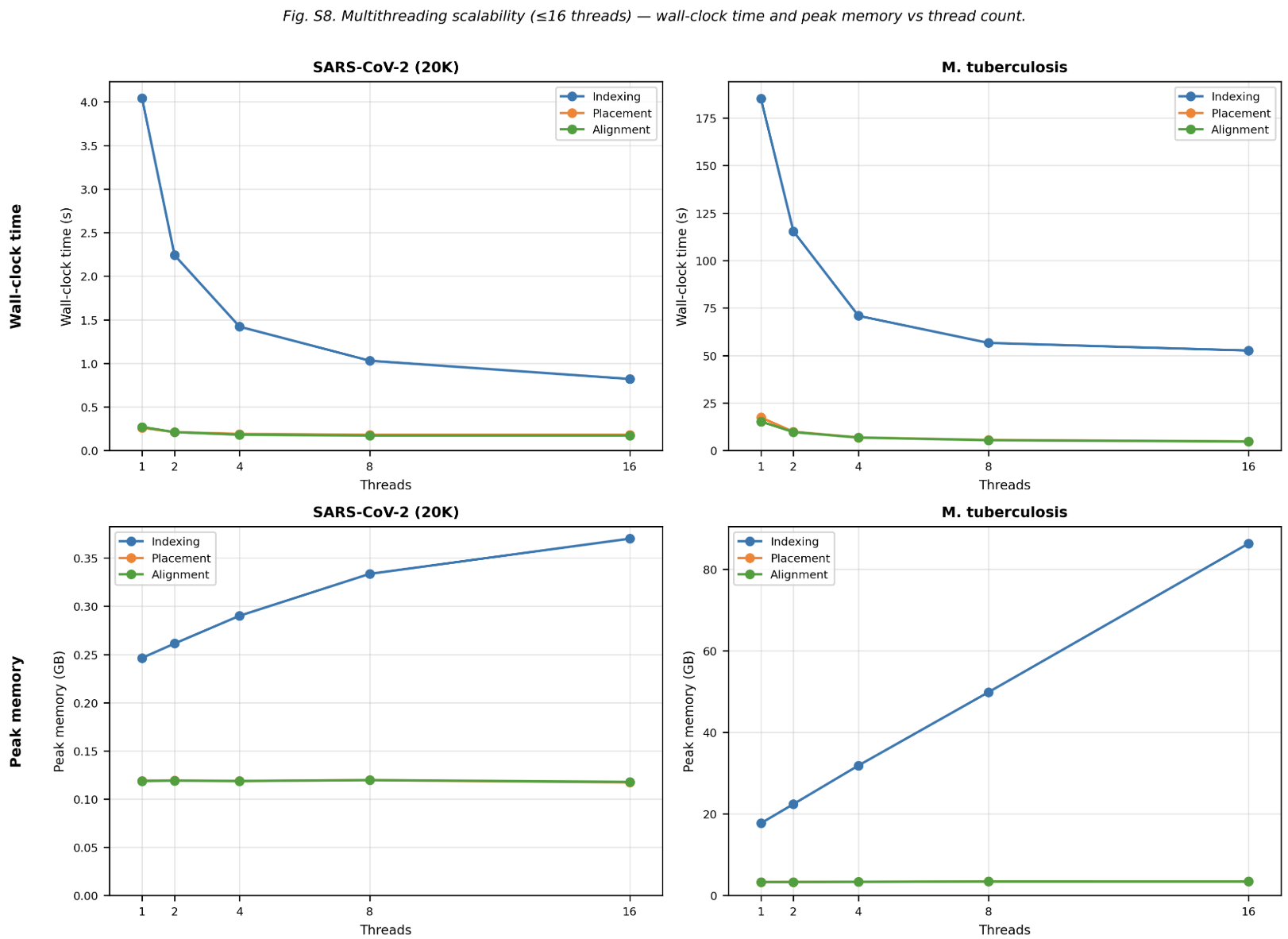

**Supplementary Figure 1. Multithreading scalability of Panmap.** Wall-clock time and peak memory usage vs. thread count for Panmap's three stages (indexing, placement, and alignment) on SARS-CoV-2 (20,000-genome PanMAN, left) and M. tuberculosis (400-genome PanMAN, right). Index construction benefits from multithreading: for SARS-CoV-2, wall time drops from ~4s (1 thread) to ~2.3s (2 threads) to ~1.5s (4 threads) to <1s (8 threads); for M. tuberculosis, from ~175s to ~110s to ~75s to ~55s over the same progression. Returns diminish beyond 8 threads, with 8-to-16 yielding only marginal improvement. Placement and alignment runtimes are effectively constant across thread counts. Peak indexing memory grows with thread count, particularly for M. tuberculosis (~18 GB at 1 thread to ~80 GB at 16), while placement and alignment memory remains flat. Based on these profiles, 4–8 threads offers the best trade-off between speed and memory for index construction.

**Figure S2:**
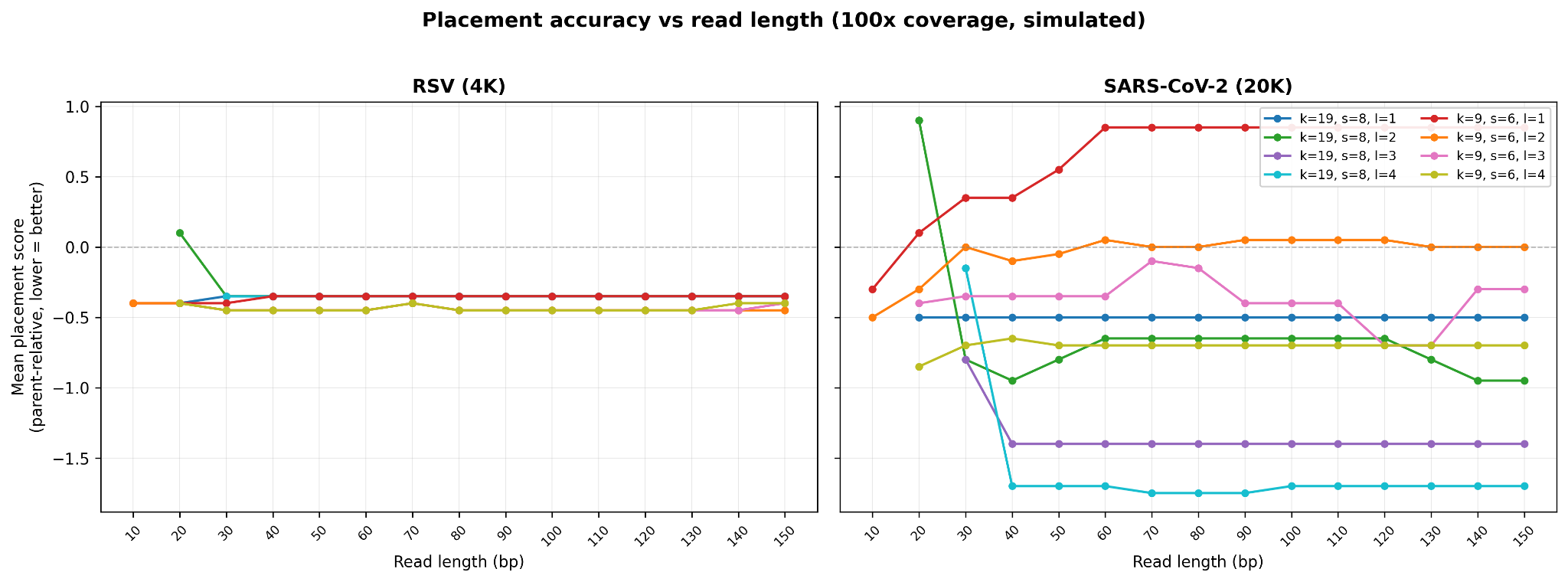

**Supplementary Figure 2. Placement accuracy vs. read length for simulated 100× coverage data**. Mean parent-relative placement score (lower is better; zero indicates placement equivalent to the true position) is shown for RSV (4,000-genome PanMAN, left) and SARS-CoV-2 (20,000-genome PanMAN, right). Each line represents a different combination of k-mer size (k) and linked syncmer chain length (l). For RSV, all configurations achieve stable, near-optimal placement across all read lengths down to ~20 bp. For SARS-CoV-2, linked syncmers (l ≥ 2) maintain substantially better accuracy at read lengths >40 bp. This demonstrates that chaining syncmers into linked seeds provides additional specificity for placement on larger, more diverse pangenomes, and that Panmap can accurately place reads far shorter than standard Illumina output.

**Figure S3:**

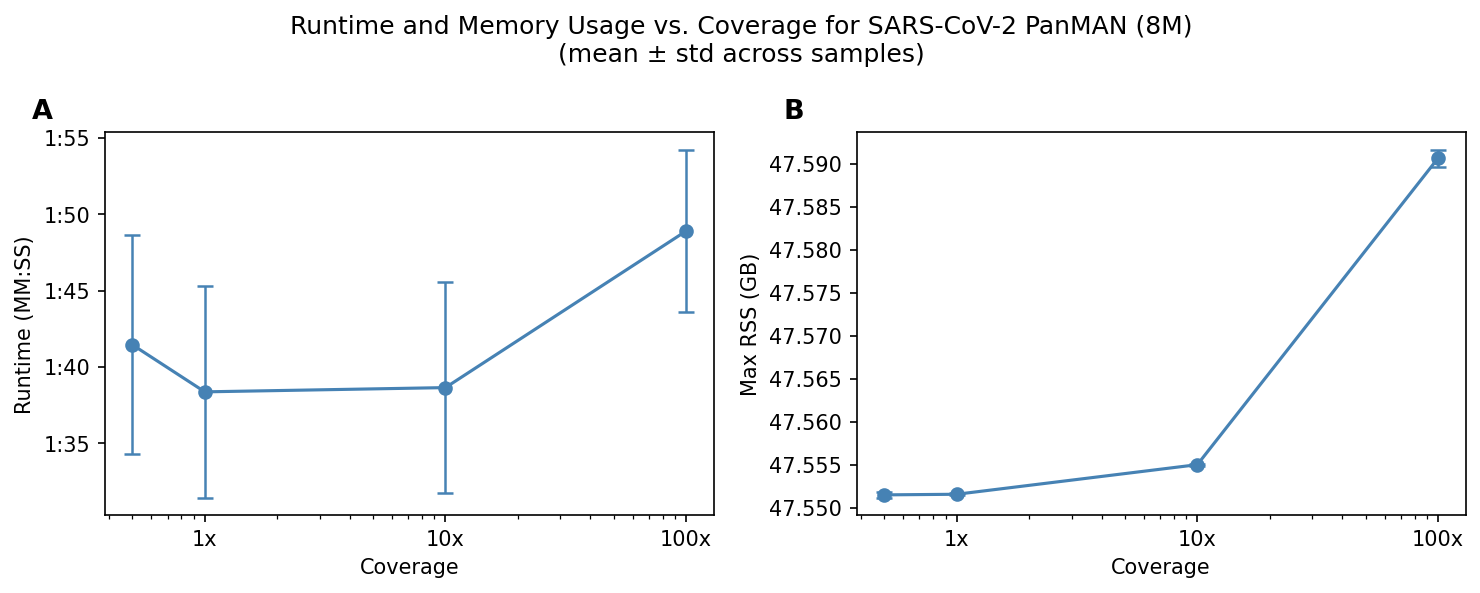

**Supplementary Figure 3. Computational resources for placing samples onto a PanMAN with 8 million genomes.** (**A**) Average runtime for placing a single SARS-CoV-2 sample with 0.5X, 1X, 10X, and 100X coverage. (**B**) Average max RSS usage for placing a single SARS-CoV-2 sample.

**Figure S4:**

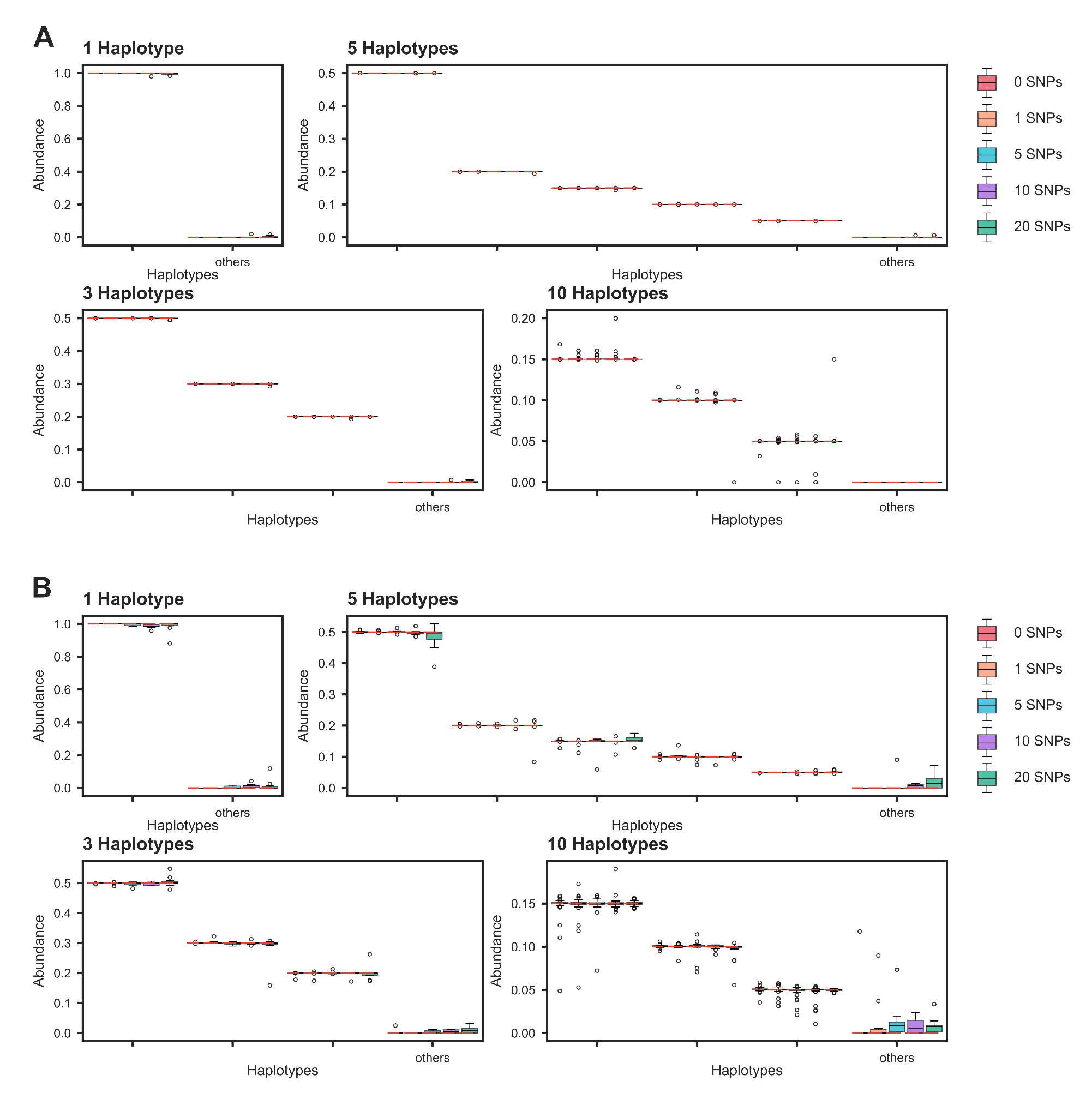

**Supplementary Figure 4. Evaluation of haplotype abundance estimation in simulated HIV (A) and RSV (B) mixture samples.** Estimated haplotype abundance in simulated HIV and RSV mixture samples with 1, 3, 5, or 10 haplotypes, introducing 0, 1, 5, 10, or 20 single-nucleotide polymorphisms (SNPs) per haplotype. Ten replicates were simulated per composition and mutation level. Red dashed lines denote true simulated abundances; the 'others' category represents accumulated false-positive haplotype abundance. (*10-haplotype samples: 4 haplotypes at 15%, 2 at 10%, 4 at 5% abundance).

**Figure S5:**

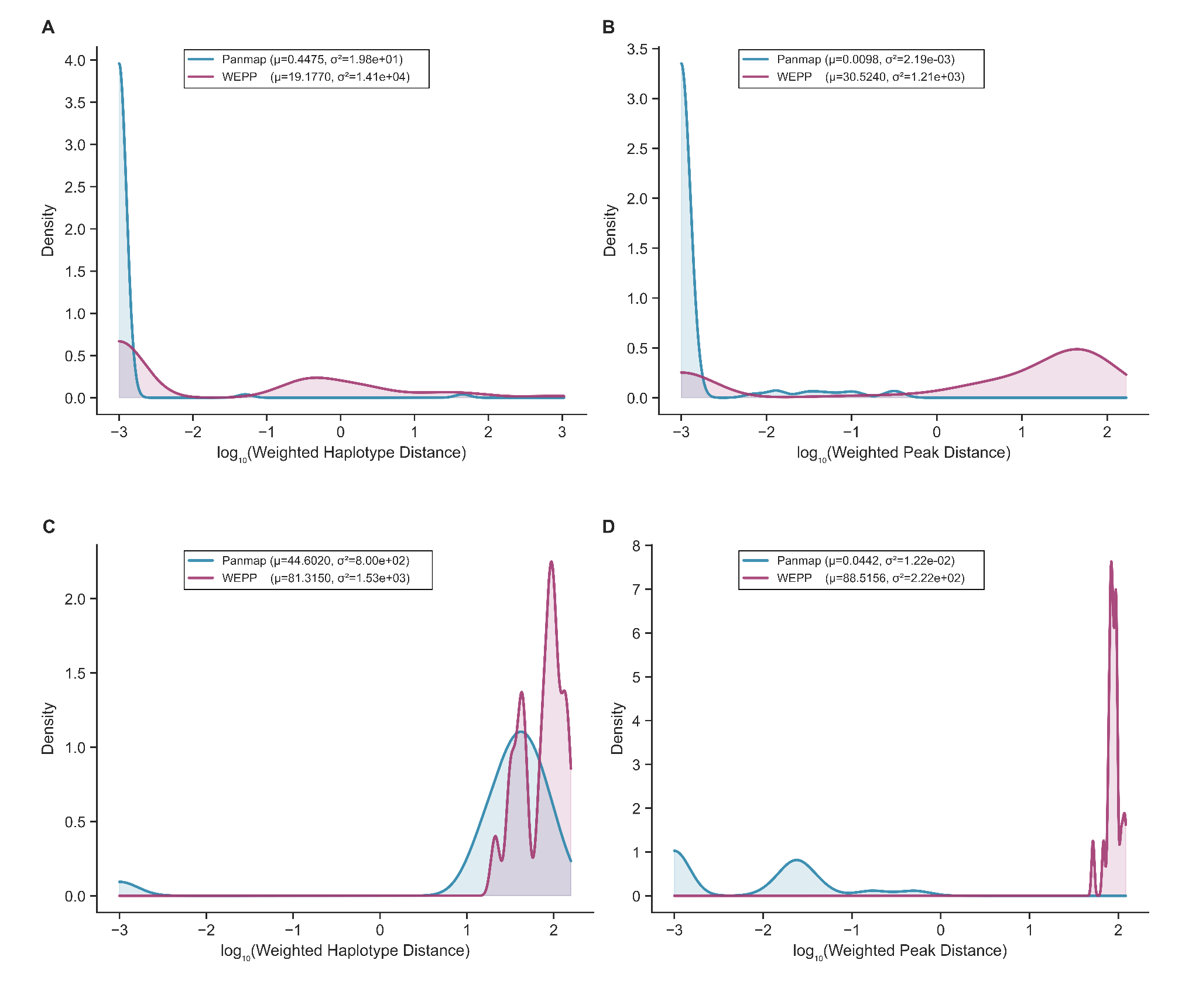

**Supplementary Figure 5. Comparison of haplotype abundance estimation accuracy between Panmap and WEPP.** Density plots show Weighted Haplotype Distance (WHD) and Weighted Peak Distance (WPD) for both methods. WHD and WPD quantify sensitivity and precision of haplotype inference, respectively, with lower values indicating greater agreement with true haplotype compositions (Gangwar et al.). (**A, B**) Samples containing 1, 2, 3, 5, or 10 haplotypes (20 replicates per mixture level; plots aggregate all replicates). (**C, D**) Samples containing 50 haplotypes under limit-of-detection testing (20 replicates total).

**Figure S6:**

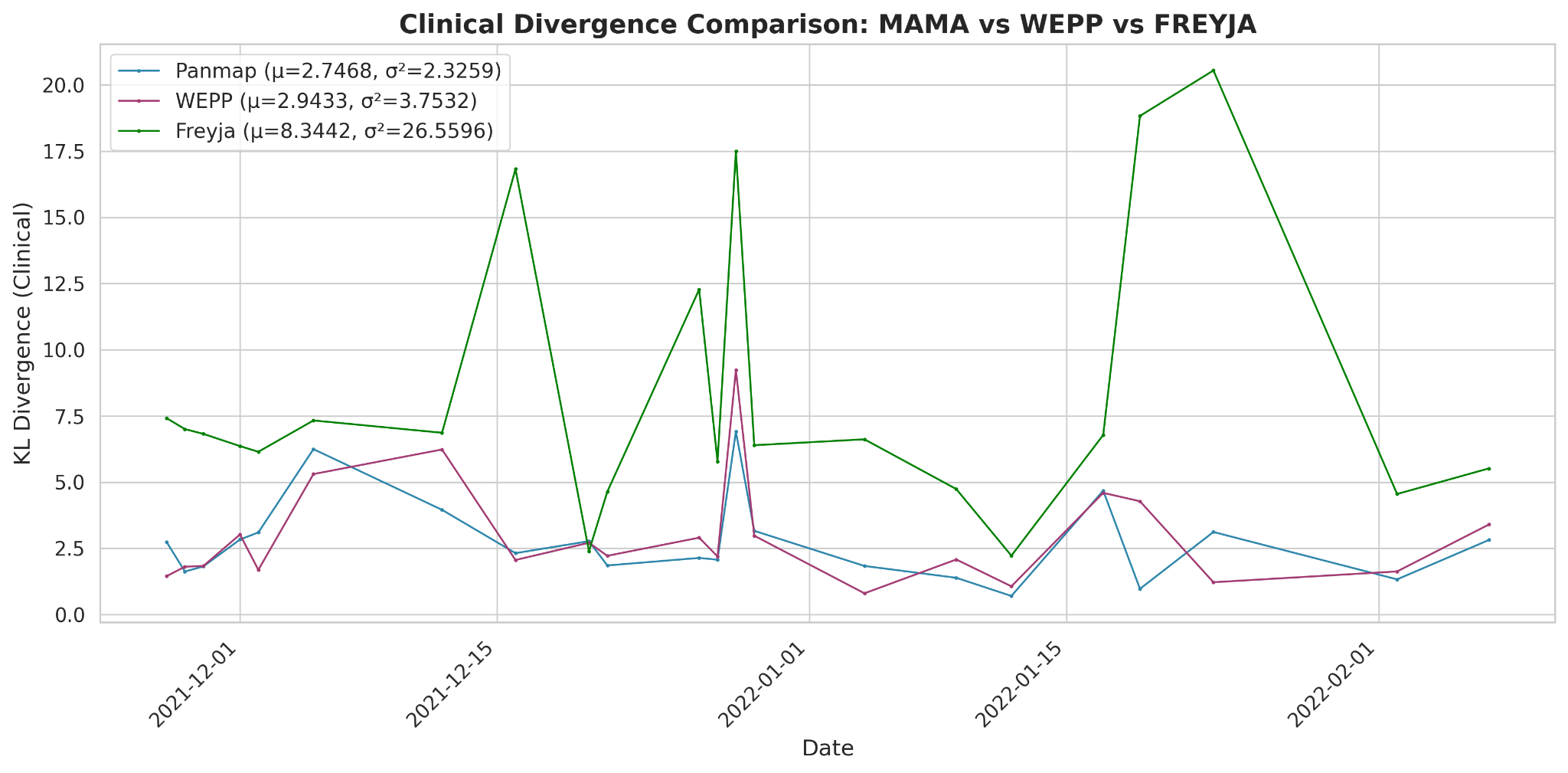

**Supplementary Figure 6. Comparison of SARS-CoV-2 lineage abundance estimation between Panmap, WEPP, and Freyja.** Time series of KL divergences between wastewater estimated lineage proportions and one-week-delayed clinically observed lineage proportions.

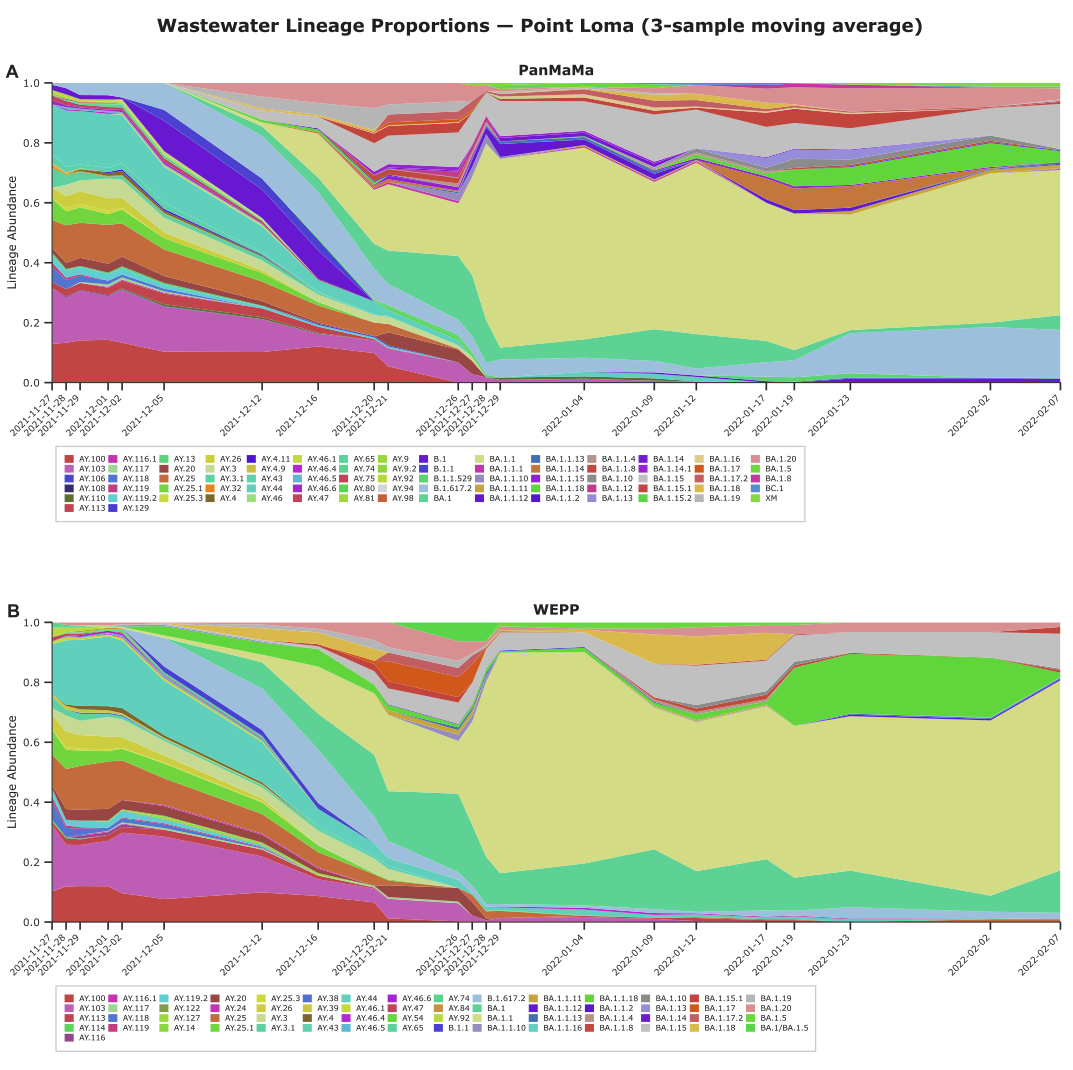
**Figure S7:**

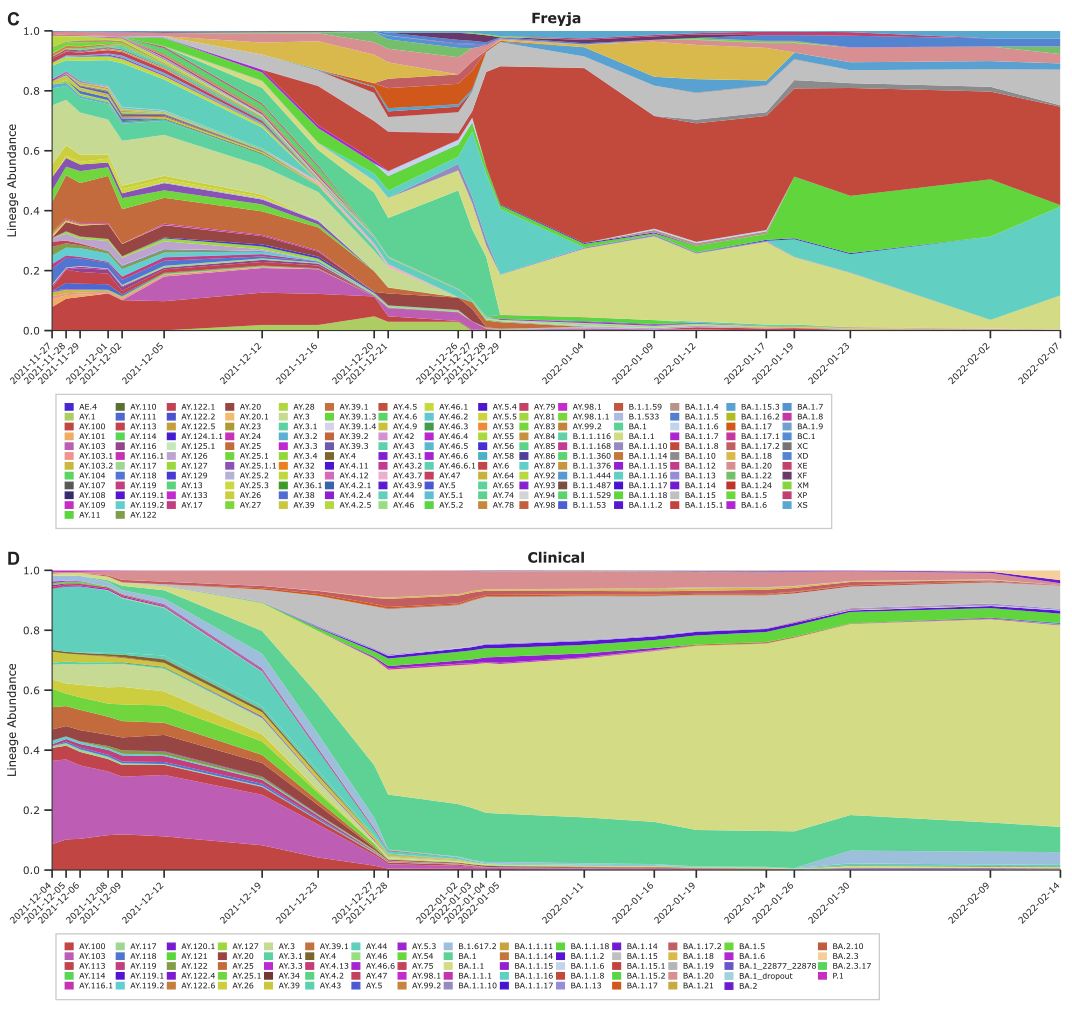

**Supplementary Figure 7. Wastewater estimated and clinically derived SARS-CoV-2 lineage proportions.** Wastewater estimated proportions by Panmap (**A**), WEPP (**B**) and Freyja (**C**) with 3-sample moving average. (**D**) Wastewater proportions derived from sequence submission.

**Figure S8:**

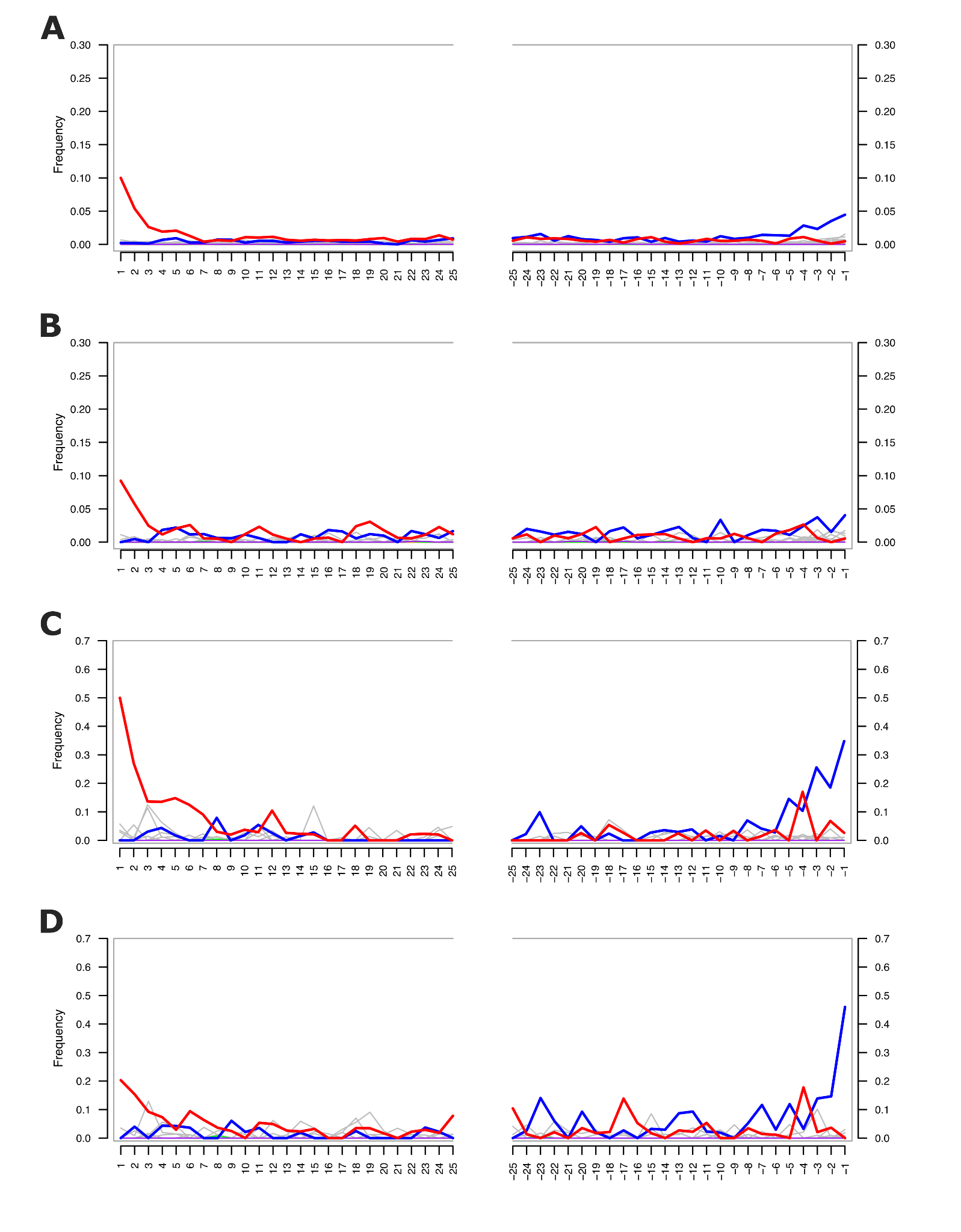

**Supplementary Figure 8. Damage plots of assigned reads generated using mapDamage** (Supp. Reference 1). Red lines indicate C-to-T substitution frequencies, blue lines G-to-A substitution frequencies, and gray lines all other substitutions.(**A**) Panmap-assigned reads from Wang et al. samples mapped to a *Mammuthus primigenius* reference genome (GenBank: DQ188829.2). (**B**) Competitive mapping–assigned reads from Wang et al. samples mapped to the same reference. (**C**) Panmap-assigned reads from Kjær et al. samples mapped to a *Mammut americanum* reference genome (GenBank: KY364233.1). (**D**) Competitive mapping–assigned reads from Kjær et al. samples mapped to the same reference.

**Figure S9:**

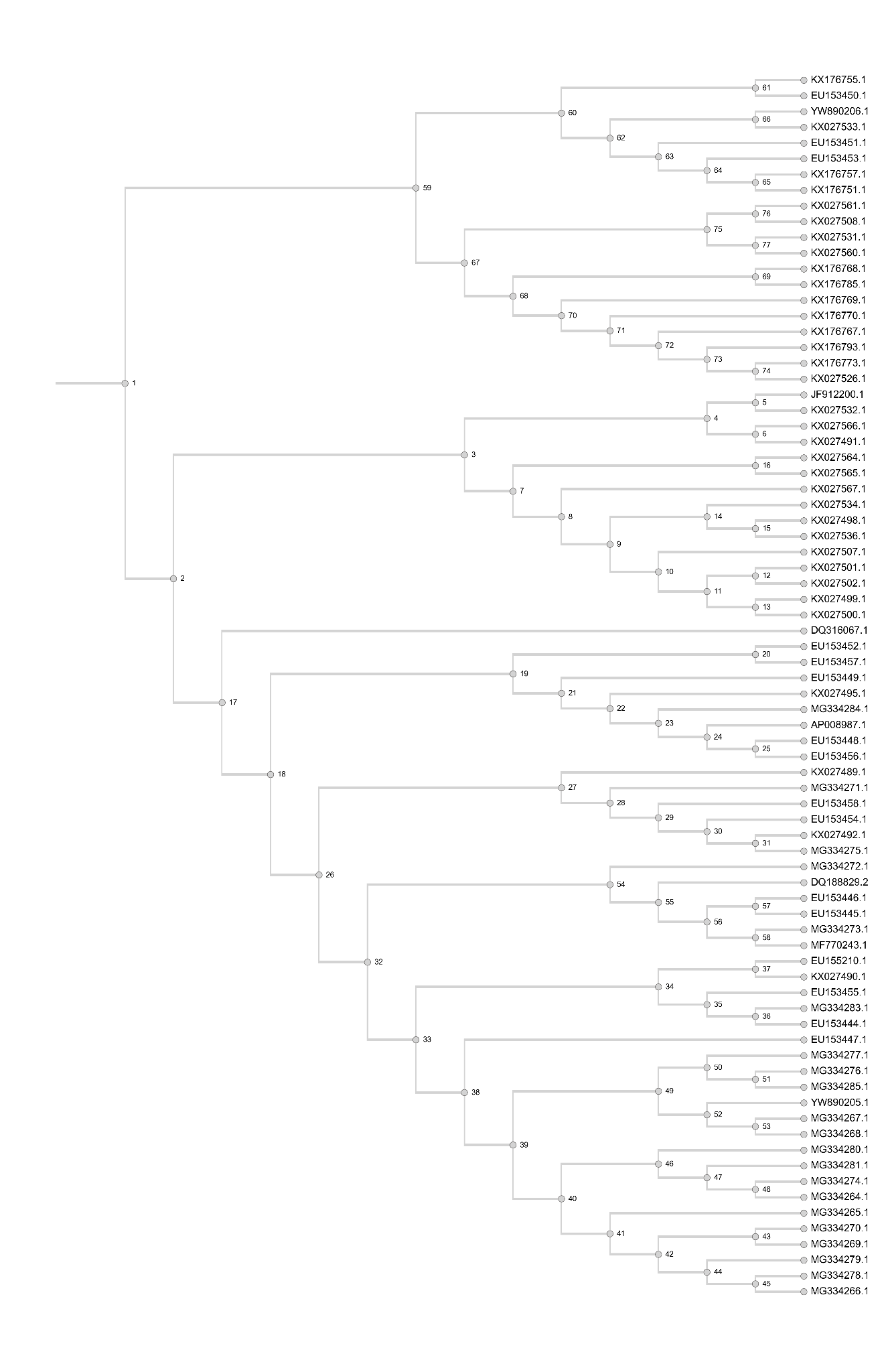

**Supplementary Figure 9. Mammuthus mitochondrial phylogeny.** Tree topology of the mammoth mitochondrial genomes. Referenced by supplementary table 4.

**Figure S10:**

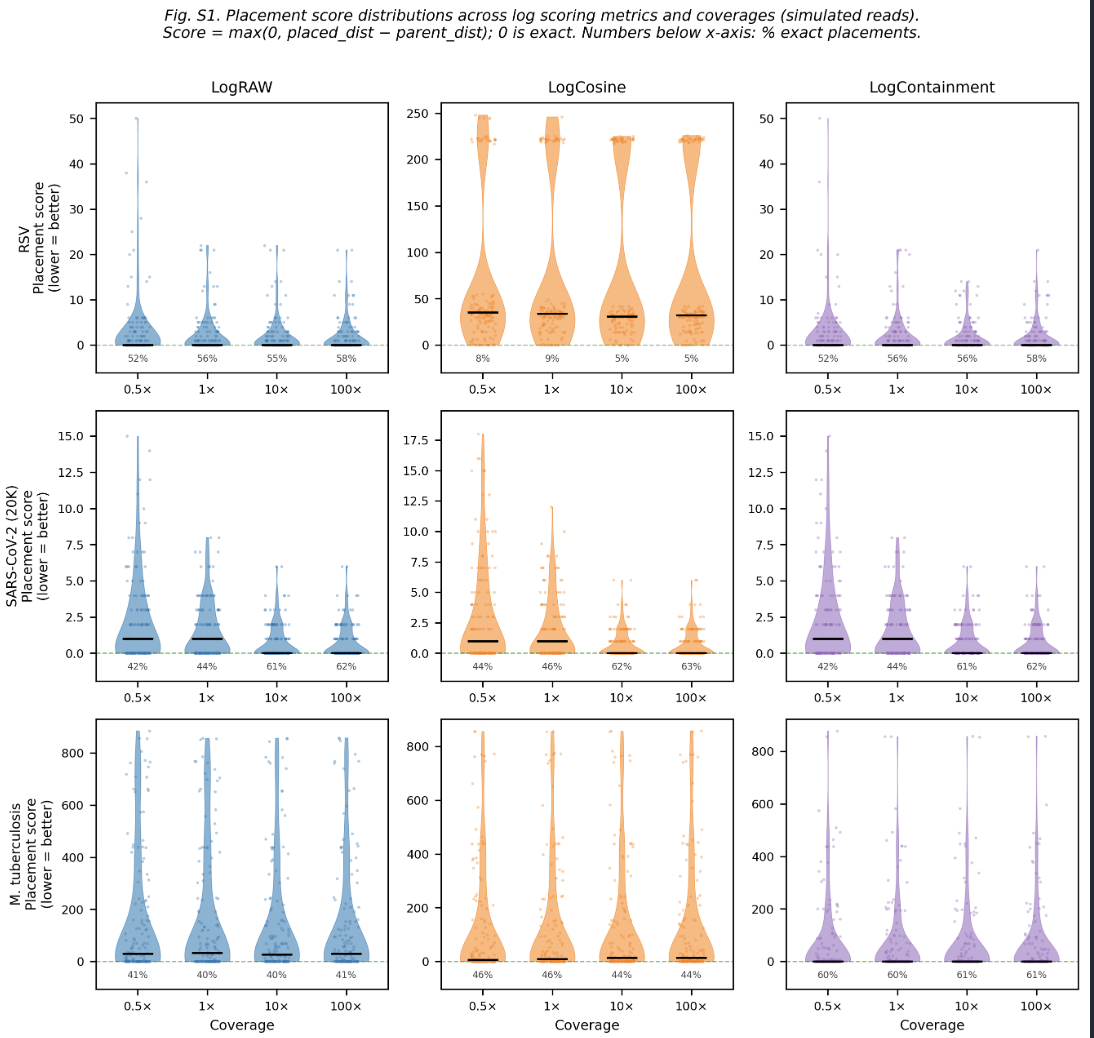

**Supplementary Figure 10. Panmap single-sample scoring metric evaluation.** Comparison of three placement scoring metrics on simulated data across RSV (4,000 genomes), SARS-CoV-2 (20,000 genomes), and M. tuberculosis (400 genomes) at four coverage levels. Violin plots show the distribution of parent-relative placement scores (lower is better; zero indicates placement at least as parsimonious as the true position); black bars indicate medians; percentages show exact-match rates (score = 0). Log containment consistently outperforms the other two metrics. These results motivated our choice of Log containment as Panmap's default placement metric.

**Table S1: Metagenomic mode performance**

| **Input pangenome** | | | | **Pangenome indexing methods** | | | | |
| --- | --- | --- | --- | --- | --- | --- | --- | --- |
| **Species** | **# Genomes** | **File format** | **Size (GZIP)** | **Tool** | **Index size (GZIP)** | **Time to index (1 thread)** | **Peak Memory (RAM) usage** | **Indexed Data** |
| RSV | 4,000 | PanMAN | 709 KB | Panmap, meta | 6.8 MB | 2.2s | 199 MB | *K-mer seeds  (syncmers) and orientations* |
| SARS-CoV-2 | 20,000 | PanMAN | 1.2 MB | Panmap, meta | 7.6 MB | 2.5s | 96 MB |  |
| SARS-CoV-2 | 8,000,000 | PanMAN | 366 MB | Panmap, meta | 2.7 GB | 37min | 68.8 GB |  |
| HIV | 20,000 | PanMAN | 13 MB | Panmap, meta | 156 MB | 56.9s | 12.7 GB |  |
| * Using parameters k=19, s=8, l=3 for all indices above | | | |  |  |  |  |  |

**Table S2: VG Giraffe read mapping performance**

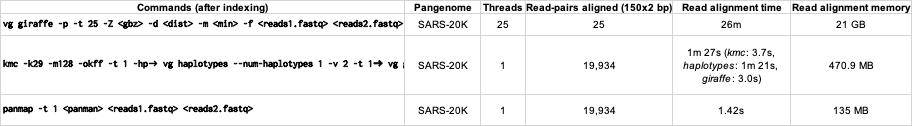

**Table S3: SRA Data used for RSV, SARS-CoV-2, *M. tuberculosis***

| **Species** | **Sample ID** | **run** | **model** | **layout** | **avglen** | **coverage** | **primer_bed** |
| --- | --- | --- | --- | --- | --- | --- | --- |
| RSV | MW033958.1 | SRR12904843 | Illumina | PAIRED | 130 | 1717451x |  |
| RSV | MF001047.1 | SRR24117175 | Illumina | PAIRED | 55 | 17588x |  |
| RSV | MG027860.1 | SRR5408822 | Illumina | PAIRED | 101 | 72553x |  |
| RSV | MG027862.1 | SRR5408827 | Illumina | PAIRED | 101 | 47889x |  |
| RSV | MG027861.1 | SRR5408828 | Illumina | PAIRED | 101 | 79496x |  |
| RSV | MG027859.1 | SRR5408840 | Illumina | PAIRED | 101 | 105501x |  |
| RSV | KY684758.1 | SRR5872102 | NA | NA | 0 | NA |  |
| RSV | KU950563.1 | SRR7822801 | Illumina | PAIRED | 86 | 3876x |  |
| RSV | KU950480.1 | SRR7822931 | Illumina | PAIRED | 252 | 6731x |  |
| RSV | KU950507.1 | SRR7823087 | Illumina | PAIRED | 188 | 2481x |  |
| RSV | KU950587.1 | SRR7823239 | Illumina | PAIRED | 192 | 2635x |  |
| RSV | KU950575.1 | SRR7823396 | Illumina | PAIRED | 229 | 8793x |  |
| RSV | KU950468.1 | SRR7823404 | Illumina | PAIRED | 259 | 5038x |  |
| RSV | KU950649.1 | SRR7823421 | Illumina | PAIRED | 236 | 5118x |  |
| RSV | KU950611.1 | SRR7823951 | Illumina | PAIRED | 183 | 1652x |  |
| RSV | KU950600.1 | SRR7823954 | Illumina | PAIRED | 200 | 8085x |  |
| RSV | KU950540.1 | SRR7823960 | Illumina | PAIRED | 198 | 7222x |  |
| RSV | KU950660.1 | SRR7823987 | Illumina | PAIRED | 171 | 4063x |  |
| RSV | KU950455.1 | SRR7824286 | Illumina | PAIRED | 169 | 1596x |  |
| RSV | KU950685.1 | SRR7824293 | Illumina | PAIRED | 186 | 5379x |  |
| RSV | KU950637.1 | SRR7824339 | Illumina | PAIRED | 183 | 4100x |  |
| RSV | KU950494.1 | SRR7824362 | Illumina | PAIRED | 172 | 6095x |  |
| RSV | KU950529.1 | SRR7824427 | Illumina | PAIRED | 250 | 2500x |  |
| RSV | KU950624.1 | SRR7824493 | Illumina | PAIRED | 253 | 4476x |  |
| RSV | KU950672.1 | SRR7824526 | Illumina | PAIRED | 123 | 2160x |  |
| RSV | KU950552.1 | SRR7824541 | Illumina | PAIRED | 205 | 3413x |  |
| RSV | SRR7824542 | SRR7824542 | Illumina | PAIRED | 250 | 3694x |  |
| RSV | KU950518.1 | SRR7824543 | Illumina | PAIRED | 200 | 1915x |  |
| RSV | SRR7824544 | SRR7824544 | Illumina | PAIRED | 172 | 4219x |  |
| RSV | SRR7824545 | SRR7824545 | Illumina | PAIRED | 206 | 1020x |  |
| RSV | SRR7824547 | SRR7824547 | Illumina | PAIRED | 145 | 5648x |  |
| RSV | SRR7824549 | SRR7824549 | Illumina | PAIRED | 240 | 3052x |  |
| RSV | SRR7824550 | SRR7824550 | Illumina | PAIRED | 251 | 3062x |  |
| RSV | SRR7824798 | SRR7824798 | Illumina | PAIRED | 247 | 3695x |  |
| RSV | SRR7824800 | SRR7824800 | Illumina | PAIRED | 259 | 1534x |  |
| RSV | SRR7824801 | SRR7824801 | Illumina | PAIRED | 189 | 6200x |  |
| RSV | SRR7824802 | SRR7824802 | Illumina | PAIRED | 257 | 5599x |  |
| RSV | SRR7824804 | SRR7824804 | Illumina | PAIRED | 202 | 4603x |  |
| RSV | SRR7824805 | SRR7824805 | Illumina | PAIRED | 271 | 4206x |  |
| RSV | SRR7824806 | SRR7824806 | Illumina | SINGLE | 205 | 50x |  |
| SARS-CoV-2 | USA/UT-UPHL-220121617557/2022\|OM459422.1\|2022-01-16 | SRR17834738 | Illumina | SINGLE | 75 | 42164x | artic_v4.1 |
| SARS-CoV-2 | USA/MA-CDCBI-CRSP_LUFUYT375FXWPR4H/2022\|OM954744.1\|2022-02-28 | SRR18281377 | Illumina | PAIRED | 75 | 8558x | artic_v4.1 |
| SARS-CoV-2 | USA/UT-UPHL-220312378164/2022\|ON082337.1\|2022-01-28 | SRR18368071 | Illumina | SINGLE | 74 | 45637x | artic_v4.1 |
| SARS-CoV-2 | USA/UT-UPHL-220326541922/2022\|ON188351.1\|2022-01-22 | SRR18656913 | Illumina | SINGLE | 74 | 50521x | artic_v4.1 |
| SARS-CoV-2 | USA/VT-Broad-CRSP_BSBAZO4APUX33JZW/2022\|ON321120.1\|2022-04-03 | SRR18886923 | Illumina | PAIRED | 75 | 9243x | artic_v4.1 |
| SARS-CoV-2 | USA/IL-CDC-ASC210750095/2022\|OM911396.1\|2022-02-21 | SRR18929960 | Illumina | SINGLE | 150 | 19498x | artic_v4.1 |
| SARS-CoV-2 | USA/MN-MDH-23670/2022\|ON364448.1\|2022-04-07 | SRR18940544 | Illumina | SINGLE | 480 | 8124x | artic_v4.1 |
| SARS-CoV-2 | USA/FL-CDC-STM-V8QF4J5MV/2022\|OM528050.1\|2022-01-19 | SRR19069645 | Illumina | PAIRED | 120 | 6557x | artic_v4.1 |
| SARS-CoV-2 | USA/FL-CDC-STM-T6A8Z5VCH/2022\|ON294457.1\|2022-04-03 | SRR19105275 | Illumina | PAIRED | 151 | 9580x | artic_v4.1 |
| SARS-CoV-2 | USA/MN-CDC-STM-K5BSUWM2C/2022\|ON232050.1\|2022-03-29 | SRR19106138 | Illumina | PAIRED | 151 | 104242x | artic_v4.1 |
| SARS-CoV-2 | USA/NY-CDC-STM-KR7KCW8TF/2022\|ON210390.1\|2022-03-28 | SRR19113011 | NA | NA | 0 | NA | artic_v4.1 |
| SARS-CoV-2 | USA/CO-CDPHE-2103200142/2022\|ON666200.1\|2022-05-04 | SRR19453763 | Illumina | SINGLE | 467 | 2006x | artic_v4.1 |
| SARS-CoV-2 | USA/CO-CDPHE-2103223795/2022\|ON667174.1\|2022-05-10 | SRR19454413 | Illumina | SINGLE | 72 | 4611x | artic_v4.1 |
| SARS-CoV-2 | USA/TG1162879/2021\|ON517574.1\|2021-12-30 | SRR19466585 | Illumina | PAIRED | 146 | 4646x | artic_v4.1 |
| SARS-CoV-2 | USA/AL-CDC-MMB14648001/2022\|ON018124.1\|2022-03-04 | SRR19690721 | Illumina | SINGLE | 177 | 11600x | artic_v4.1 |
| SARS-CoV-2 | USA/CO-CDC-MMB14765192/2022\|ON242429.1\|2022-03-17 | SRR19692782 | Illumina | SINGLE | 171 | 6957x | artic_v4.1 |
| SARS-CoV-2 | USA/MO-CDC-QDX36965205/2022\|ON698767.1\|2022-05-10 | SRR19693862 | Illumina | PAIRED | 251 | 8533x | artic_v4.1 |
| SARS-CoV-2 | USA/IN-CDC-STM-DEQAFX6DJ/2022\|ON689495.1\|2022-05-13 | SRR19705377 | Illumina | PAIRED | 132 | 38257x | artic_v4.1 |
| SARS-CoV-2 | USA/CA-CDC-QDX37346317/2022\|ON744262.1\|2022-05-22 | SRR19720358 | Illumina | PAIRED | 251 | 23533x | artic_v4.1 |
| SARS-CoV-2 | USA/FL-CDC-LC0706152/2022\|ON821724.1\|2022-06-10 | SRR19796720 | ONT | SINGLE | 729 | 126x | artic_v4.1 |
| SARS-CoV-2 | USA/NC-CDC-MMB14248708/2022\|OM806905.1\|2022-02-08 | SRR19831890 | Illumina | SINGLE | 193 | 5808x | artic_v4.1 |
| SARS-CoV-2 | USA/CA-CDC-LC0703578/2022\|ON824163.1\|2022-06-03 | SRR19839995 | ONT | SINGLE | 726 | 116x | artic_v4.1 |
| SARS-CoV-2 | USA/CO-CDC-MMB14065750/2022\|OM846676.1\|2022-01-31 | SRR19842654 | Illumina | SINGLE | 198 | 8341x | artic_v4.1 |
| SARS-CoV-2 | USA/PA-BOL-M22095157/2022\|ON875980.1\|2022-06-17 | SRR19902572 | Illumina | PAIRED | 100 | 2294x | artic_v4.1 |
| SARS-CoV-2 | USA/TX-CDC-QDX37663357/2022\|ON828662.1\|2022-06-03 | SRR19946282 | Illumina | PAIRED | 251 | 10800x | artic_v4.1 |
| SARS-CoV-2 | USA/TX-CDC-LC0708391/2022\|ON837325.1\|2022-06-06 | SRR19947961 | ONT | SINGLE | 733 | 303x | artic_v4.1 |
| SARS-CoV-2 | USA/MI-CDC-QDX37912287/2022\|ON867733.1\|2022-06-09 | SRR19959413 | Illumina | PAIRED | 249 | 14004x | artic_v4.1 |
| SARS-CoV-2 | USA/NV-CDC-QDX38007139/2022\|ON905841.1\|2022-06-09 | SRR20118704 | Illumina | PAIRED | 250 | 15657x | artic_v4.1 |
| SARS-CoV-2 | USA/CA-CDC-QDX38007692/2022\|ON906540.1\|2022-06-12 | SRR20119416 | Illumina | PAIRED | 250 | 22643x | artic_v4.1 |
| SARS-CoV-2 | USA/OH-CDC-LC0720653/2022\|ON894608.1\|2022-06-10 | SRR20122995 | ONT | SINGLE | 726 | 493x | artic_v4.1 |
| SARS-CoV-2 | USA/NJ-CDC-QDX37054611/2022\|ON925775.1\|2022-05-16 | SRR20124299 | Illumina | PAIRED | 250 | 11217x | artic_v4.1 |
| SARS-CoV-2 | USA/IL-CDC-LC0749472/2022\|ON975722.1\|2022-06-26 | SRR20222422 | ONT | SINGLE | 727 | 334x | artic_v4.1 |
| SARS-CoV-2 | USA/CA-CDC-QDX38473921/2022\|OP009025.1\|2022-06-27 | SRR20305299 | Illumina | PAIRED | 246 | 9423x | artic_v4.1 |
| SARS-CoV-2 | USA/CA-CDC-STM-V5FZQ5EH7/2022\|ON546851.1\|2022-05-07 | SRR20343333 | Illumina | PAIRED | 151 | 222x | artic_v4.1 |
| SARS-CoV-2 | USA/WA-CDC-QDX35917447/2022\|ON382673.1\|2022-04-12 | SRR20344359 | Illumina | PAIRED | 248 | 19447x | artic_v4.1 |
| SARS-CoV-2 | USA/GA-CDC-LC0464293/2022\|OM360598.1\|2022-01-04 | SRR20368218 | ONT | SINGLE | 731 | 108x | artic_v4.1 |
| SARS-CoV-2 | USA/TX-CDC-STM-QXCMG2NM5/2022\|OP025786.1\|2022-07-09 | SRR20376169 | Illumina | PAIRED | 151 | 86871x | artic_v4.1 |
| SARS-CoV-2 | USA/NJ-CDC-LC0589887/2022\|ON415160.1\|2022-04-24 | SRR20381292 | ONT | SINGLE | 725 | 177x | artic_v4.1 |
| SARS-CoV-2 | USA/NJ-CDC-LC0657992/2022\|ON659908.1\|2022-05-22 | SRR20384316 | ONT | SINGLE | 722 | 128x | artic_v4.1 |
| SARS-CoV-2 | USA/OR-CDC-LC0659687/2022\|ON657128.1\|2022-05-18 | SRR20386790 | ONT | SINGLE | 724 | 99x | artic_v4.1 |
| SARS-CoV-2 | USA/NC-CDC-QDX33832252/2022\|OM843825.1\|2022-02-12 | SRR20391345 | Illumina | PAIRED | 251 | 17381x | artic_v4.1 |
| SARS-CoV-2 | USA/FL-CDC-QDX34290221/2022\|OM998480.1\|2022-02-23 | SRR20393485 | Illumina | PAIRED | 251 | 26838x | artic_v4.1 |
| SARS-CoV-2 | USA/RI-CDC-LC0613487/2022\|ON538070.1\|2022-05-04 | SRR20411044 | ONT | SINGLE | 726 | 297x | artic_v4.1 |
| SARS-CoV-2 | USA/CA-CDC-LC0601597/2022\|ON466320.1\|2022-04-25 | SRR20431711 | ONT | SINGLE | 739 | 191x | artic_v4.1 |
| SARS-CoV-2 | USA/IL-CDC-LC0609004/2022\|ON499532.1\|2022-04-30 | SRR20440468 | ONT | SINGLE | 738 | 167x | artic_v4.1 |
| SARS-CoV-2 | USA/CA-CDC-LC0606441/2022\|ON499645.1\|2022-04-30 | SRR20441961 | ONT | SINGLE | 710 | 161x | artic_v4.1 |
| SARS-CoV-2 | USA/AK-CDC-LC0569589/2022\|ON232854.1\|2022-03-28 | SRR20444097 | ONT | SINGLE | 732 | 807x | artic_v4.1 |
| SARS-CoV-2 | USA/CA-CDC-LC0573891/2022\|ON297362.1\|2022-04-10 | SRR20451428 | ONT | SINGLE | 711 | 227x | artic_v4.1 |
| SARS-CoV-2 | USA/CT-CDC-LC0572468/2022\|ON295798.1\|2022-04-06 | SRR20452975 | ONT | SINGLE | 667 | 419x | artic_v4.1 |
| SARS-CoV-2 | USA/CT-CDC-LC0575831/2022\|ON297310.1\|2022-04-10 | SRR20453183 | ONT | SINGLE | 732 | 212x | artic_v4.1 |
| SARS-CoV-2 | USA/TX-CDC-QDX35823923/2022\|ON336360.1\|2022-04-10 | SRR20505101 | Illumina | PAIRED | 249 | 75332x | artic_v4.1 |
| SARS-CoV-2 | USA/TX-CDC-QDX35823919/2022\|ON336345.1\|2022-04-10 | SRR20505104 | Illumina | PAIRED | 250 | 8996x | artic_v4.1 |
| SARS-CoV-2 | USA/NJ-CDC-QDX35823774/2022\|ON336067.1\|2022-04-10 | SRR20505166 | Illumina | PAIRED | 250 | 8357x | artic_v4.1 |
| SARS-CoV-2 | USA/NM-CDC-QDX35734163/2022\|ON334769.1\|2022-04-05 | SRR20506322 | Illumina | PAIRED | 251 | 41229x | artic_v4.1 |
| SARS-CoV-2 | USA/MD-CDC-LC0511880/2022\|OM627630.1\|2022-01-27 | SRR20511034 | ONT | SINGLE | 733 | 538x | artic_v4.1 |
| SARS-CoV-2 | USA/DE-CDC-LC0511879/2022\|OM627617.1\|2022-01-27 | SRR20511035 | ONT | SINGLE | 729 | 487x | artic_v4.1 |
| SARS-CoV-2 | USA/FL-CDC-QDX34933928/2022\|ON111985.1\|2022-03-15 | SRR20518533 | Illumina | PAIRED | 250 | 10296x | artic_v4.1 |
| SARS-CoV-2 | USA/NJ-CDC-QDX35420258/2022\|ON270704.1\|2022-03-30 | SRR20536617 | Illumina | PAIRED | 251 | 9638x | artic_v4.1 |
| SARS-CoV-2 | USA/NY-CDC-QDX35868502/2022\|ON363545.1\|2022-04-11 | SRR20537235 | Illumina | PAIRED | 250 | 17531x | artic_v4.1 |
| SARS-CoV-2 | USA/PA-CDC-QDX35733785/2022\|ON362676.1\|2022-04-07 | SRR20537676 | Illumina | PAIRED | 249 | 6349x | artic_v4.1 |
| SARS-CoV-2 | USA/CA-CDC-QDX35917541/2022\|ON363764.1\|2022-04-12 | SRR20537734 | Illumina | PAIRED | 61 | 872x | artic_v4.1 |
| SARS-CoV-2 | USA/FL-CDC-QDX35624181/2022\|ON311591.1\|2022-04-05 | SRR20565507 | Illumina | PAIRED | 250 | 13909x | artic_v4.1 |
| SARS-CoV-2 | USA/IL-CDC-QDX35624703/2022\|ON311625.1\|2022-04-05 | SRR20565694 | Illumina | PAIRED | 250 | 8373x | artic_v4.1 |
| SARS-CoV-2 | USA/MD-CDC-QDX36111259/2022\|ON452162.1\|2022-04-19 | SRR20568925 | Illumina | PAIRED | 251 | 12821x | artic_v4.1 |
| SARS-CoV-2 | USA/OH-CDC-QDX36112176/2022\|ON452104.1\|2022-04-19 | SRR20569961 | Illumina | PAIRED | 250 | 7092x | artic_v4.1 |
| SARS-CoV-2 | USA/PA-CDC-QDX35974479/2022\|ON458293.1\|2022-04-14 | SRR20572197 | Illumina | PAIRED | 251 | 9618x | artic_v4.1 |
| SARS-CoV-2 | USA/PA-CDC-QDX36315904/2022\|ON490095.1\|2022-04-25 | SRR20575074 | Illumina | PAIRED | 250 | 11886x | artic_v4.1 |
| SARS-CoV-2 | USA/IL-CDC-STM-KXZXT8XKR/2022\|ON655554.1\|2022-05-21 | SRR20581721 | Illumina | PAIRED | 146 | 6331x | artic_v4.1 |
| SARS-CoV-2 | USA/FL-CDC-LC0789043/2022\|OP110610.1\|2022-07-18 | SRR20740244 | ONT | SINGLE | 722 | 481x | artic_v4.1 |
| SARS-CoV-2 | USA/PR-CDC-ASC210853121/2022\|ON309432.1\|2022-04-07 | SRR20774109 | Illumina | PAIRED | 151 | 104088x | artic_v4.1 |
| SARS-CoV-2 | USA/PR-CDC-ASC210853120/2022\|ON309433.1\|2022-04-07 | SRR20774110 | NA | NA | 0 | NA | artic_v4.1 |
| SARS-CoV-2 | USA/PA-CDC-QDX36714295/2022\|ON574441.1\|2022-05-07 | SRR20781737 | Illumina | PAIRED | 248 | 19014x | artic_v4.1 |
| SARS-CoV-2 | USA/PA-CDC-QDX36606629/2022\|ON602567.1\|2022-05-02 | SRR20794843 | Illumina | PAIRED | 250 | 29625x | artic_v4.1 |
| SARS-CoV-2 | USA/MD-CDC-LC0477794/2022\|OM413029.1\|2022-01-12 | SRR20865142 | ONT | SINGLE | 723 | 799x | artic_v4.1 |
| SARS-CoV-2 | USA/CO-CDC-ASC210562243/2021\|OM290822.1\|2021-12-29 | SRR20870158 | NA | NA | 0 | NA | artic_v4.1 |
| SARS-CoV-2 | USA/NC-CDC-MMB12597453/2021\|OM414019.1\|2021-12-29 | SRR20890703 | Illumina | SINGLE | 260 | 11306x | artic_v4.1 |
| SARS-CoV-2 | USA/NC-CDC-MMB12603839/2021\|OM414032.1\|2021-12-29 | SRR20891254 | Illumina | SINGLE | 252 | 10500x | artic_v4.1 |
| SARS-CoV-2 | USA/CA-CDC-STM-AN8HZVPNB/2022\|OP147609.1\|2022-07-19 | SRR20909109 | Illumina | PAIRED | 151 | 2419x | artic_v4.1 |
| SARS-CoV-2 | USA/CA-CDC-QDX39078984/2022\|OP126264.1\|2022-07-13 | SRR20915991 | Illumina | PAIRED | 250 | 5494x | artic_v4.1 |
| SARS-CoV-2 | USA/NJ-CDC-LC0802035/2022\|OP167787.1\|2022-07-26 | SRR20955738 | ONT | SINGLE | 723 | 108x | artic_v4.1 |
| SARS-CoV-2 | USA/NM-CDC-QDX39214603/2022\|OP170782.1\|2022-07-17 | SRR20980219 | Illumina | PAIRED | 242 | 15284x | artic_v4.1 |
| SARS-CoV-2 | USA/CO-CDC-MMB14712265/2022\|ON068238.1\|2022-03-11 | SRR21087389 | Illumina | SINGLE | 196 | 7380x | artic_v4.1 |
| SARS-CoV-2 | USA/NM-CDC-2-6165322/2022\|ON998335.1\|2022-06-15 | SRR21131034 | Illumina | PAIRED | 106 | 11648x | artic_v4.1 |
| SARS-CoV-2 | USA/CO-CDC-2-6261958/2022\|OP292335.1\|2022-07-08 | SRR21230834 | Illumina | PAIRED | 99 | 10556x | artic_v4.1 |
| SARS-CoV-2 | USA/CA-CDC-LC0822310/2022\|OP241210.1\|2022-08-04 | SRR21242789 | ONT | SINGLE | 726 | 147x | artic_v4.1 |
| SARS-CoV-2 | USA/NC-CDC-MMB12210161/2021\|OM047982.1\|2021-12-16 | SRR21263251 | Illumina | SINGLE | 222 | 7159x | artic_v4.1 |
| SARS-CoV-2 | USA/NJ-CDC-ASC210850559/2022\|ON252304.1\|2022-04-04 | SRR21340670 | Illumina | PAIRED | 151 | 220454x | artic_v4.1 |
| SARS-CoV-2 | USA/FL-CDC-ASC210721481/2022\|ON250356.1\|2022-03-30 | SRR21341513 | Illumina | PAIRED | 127 | 183693x | artic_v4.1 |
| SARS-CoV-2 | USA/PR-CDC-ASC210721769/2022\|ON236379.1\|2022-03-31 | SRR21343215 | Illumina | PAIRED | 128 | 193326x | artic_v4.1 |
| SARS-CoV-2 | USA/OR-CDC-ASC210859697/2022\|OM862675.1\|2022-02-16 | SRR21344901 | Illumina | PAIRED | 151 | 190880x | artic_v4.1 |
| SARS-CoV-2 | USA/NY-CDC-ASC210737849/2022\|ON144422.1\|2022-03-22 | SRR21346832 | Illumina | PAIRED | 151 | 250535x | artic_v4.1 |
| SARS-CoV-2 | USA/MP-CDC-2-5839144/2022\|ON318629.1\|2022-03-10 | SRR21432734 | Illumina | PAIRED | 94 | 2081x | artic_v4.1 |
| SARS-CoV-2 | USA/MP-CDC-2-5838738/2022\|ON318500.1\|2022-03-06 | SRR21432827 | Illumina | PAIRED | 97 | 10836x | artic_v4.1 |
| SARS-CoV-2 | USA/MP-CDC-2-5876378/2022\|ON384212.1\|2022-03-25 | SRR21433337 | Illumina | PAIRED | 97 | 2623x | artic_v4.1 |
| SARS-CoV-2 | USA/VI-CDC-2-5419238/2021\|OM461145.1\|2021-12-29 | SRR21437987 | Illumina | PAIRED | 104 | 10559x | artic_v4.1 |
| SARS-CoV-2 | USA/VI-CDC-2-5419304/2021\|OM461240.1\|2021-12-31 | SRR21438201 | Illumina | PAIRED | 104 | 7481x | artic_v4.1 |
| SARS-CoV-2 | USA/VI-CDC-2-5419245/2021\|OM461234.1\|2021-12-31 | SRR21438206 | Illumina | PAIRED | 103 | 755x | artic_v4.1 |
| SARS-CoV-2 | USA/MP-CDC-2-5802637/2022\|ON260323.1\|2022-02-28 | SRR21439112 | Illumina | PAIRED | 105 | 8420x | artic_v4.1 |
| SARS-CoV-2 | USA/FL-CDC-STM-9M5J7ENZ9/2022\|ON389309.1\|2022-04-15 | SRR21652136 | Illumina | PAIRED | 139 | 4414x | artic_v4.1 |
| SARS-CoV-2 | USA/WA-PHL-018764/2022\|OQ982119.1\|2022-04-15 | SRR29129247 | Illumina | PAIRED | 76 | 16868x | artic_v4.1 |
| M. tuberculosis | NZ_CP048071.1 | SRR10828834 | NA | NA | 0 | NA |  |
| M. tuberculosis | NZ_CP069063.1 | SRR12801730 | ONT | SINGLE | 5816 | 197x |  |
| M. tuberculosis | NZ_CP069064.1 | SRR12801731 | Illumina | SINGLE | 222 | 353x |  |
| M. tuberculosis | CP069065.1 | SRR12801732 | ONT | SINGLE | 5004 | 182x |  |
| M. tuberculosis | NZ_CP069067.1 | SRR12801734 | Illumina | SINGLE | 266 | 49x |  |
| M. tuberculosis | CP069068.1 | SRR12801735 | ONT | SINGLE | 7400 | 133x |  |
| M. tuberculosis | NZ_CP069069.1 | SRR12801736 | ONT | SINGLE | 3539 | 254x |  |
| M. tuberculosis | NZ_CP069077.1 | SRR12801737 | ONT | SINGLE | 1209 | 181x |  |
| M. tuberculosis | CP069078.1 | SRR12801738 | ONT | SINGLE | 1479 | 45x |  |
| M. tuberculosis | NZ_CP089776.1 | SRR17234883 | Illumina | PAIRED | 125 | 214x |  |
| M. tuberculosis | NZ_CP089773.1 | SRR17234890 | NA | NA | 0 | NA |  |
| M. tuberculosis | NZ_CP089779.1 | SRR17234892 | Illumina | PAIRED | 125 | 115x |  |
| M. tuberculosis | CP044345.1 | SRR20995191 | Illumina | PAIRED | 158 | 94x |  |
| M. tuberculosis | NZ_CP117298.1 | SRR23325069 | Illumina | PAIRED | 225 | 172x |  |
| M. tuberculosis | CP097109.1 | SRR28776755 | Illumina | PAIRED | 149 | 38x |  |
| M. tuberculosis | NZ_CP154612.1 | SRR34323111 | ONT | SINGLE | 3389 | 349x |  |
| M. tuberculosis | CP003494.1 | SRR398632 | NA | NA | 0 | NA |  |
| M. tuberculosis | NZ_CP027035.1 | SRR6467888 | NA | NA | 0 | NA |  |
| M. tuberculosis | CP015773.2 | SRR6705904 | Illumina | PAIRED | 100 | 977x |  |
| M. tuberculosis | NZ_CP039851.1 | SRR7983753 | NA | NA | 0 | NA |  |
| M. tuberculosis | NZ_CP039850.1 | SRR7983755 | NA | NA | 0 | NA |  |

**Table S4: PanMAN sources and details**

| **Species** | **# genomes** | **Pangenome format** | **panmanutils summary output** | | | | | | | | | |
| --- | --- | --- | --- | --- | --- | --- | --- | --- | --- | --- | --- | --- |
|  |  |  | **Nodes** | **Substitutions** | **Insertions** | **Deletions** | **Inversions** | **Block Insertions** | **Block Deletions** | **Block Inversions** | **Block Duplications** | **Block Translocations** |
| *M. tuberculosis* | 400 | PanMAN, previously published data (Walia et al 2026) | 799 | 279,004 | 907,824 | 372,695 | 146,440 | 447,527 | 187,046 | 169 | 32,027 | 148,375 |
| RSV | 4,000 | PanMAN, previously published data | 7,999 | 149,267 | 37,071 | 28,657 | 93 | 6,114 | 7,448 | 0 | 21 | 3,248 |
| HIV | 20,000 | PanMAN, previously published data | 39,999 | 2,548,540 | 1,376,073 | 274,759 | 156 | 350,968 | 90,472 | 1 | 1,102 | 78,764 |
| SARS-CoV-2 | 20,000 | PanMAN built with TWILIGHT MSA and DIPPER guide tree | 39,999 | 219,232 | 139,424 | 52,255 | 0 | 1 | 0 | 0 | 0 | 0 |
| SARS-CoV-2 | 8,000,000 | PanMAN built with TWILIGHT MSA and DIPPER guide tree | 9,606,658 | 1,229,102,156 | 168,097,395 | 225,692,340 | 0 | 1 | 0 | 0 | 0 | 0 |
| Mitochondria (Vertebrata) | 15,654 | PanMAN built with TWILIGHT MSA and DIPPER guide tree | 31,207 | 8,757,319 | 731,233 | 524,439 | 0 | 1 | 0 | 0 | 0 | 0 |
| Mitochondria (Mammuthus spp.) | 72 | PanMAN built with TWILIGHT MSA and guide tree published by Martiniano et al. | 155 | 8,970 | 937 | 7,096 | 0 | 1 | 0 | 0 | 0 | 0 |

**Table S5: Wang et al. per sample mammoth placement**

|  | **Panmap** | | | **PathPhynder** | | | **Comparison** | |
| --- | --- | --- | --- | --- | --- | --- | --- | --- |
| **Sample** | **Placement** | **Node depth** | **Reads** | **Placement** | **Node depth** | **Reads** | **Same subtree** | **Node distance** |
| ar1_6 | node_18 | 3 | 3 | NA | NA | 0 | FALSE | NA |
| cr8_1 | NA | NA | 0 | NA | NA | 0 | NA | NA |
| cr6_2 | NA | NA | 0 | NA | NA | 0 | NA | NA |
| cr4_25 | NA | NA | 0 | NA | NA | 0 | NA | NA |
| tm9_8 | NA | NA | 0 | NA | NA | 0 | NA | NA |
| cr8_15 | NA | NA | 0 | NA | NA | 0 | NA | NA |
| cr2_7 | NA | NA | 0 | NA | NA | 0 | NA | NA |
| cr5_14 | NA | NA | 0 | NA | NA | 0 | NA | NA |
| cr2_2 | node_1 | 0 | 1 | NA | NA | 0 | FALSE | NA |
| cr2_32 | NA | NA | 0 | NA | NA | 0 | NA | NA |
| cr7_2 | NA | NA | 0 | node_17 | 2 | 1 | FALSE | NA |
| cr6_6 | NA | NA | 0 | NA | NA | 0 | NA | NA |
| tm3_5 | NA | NA | 0 | NA | NA | 0 | NA | NA |
| tm9_9 | NA | NA | 0 | NA | NA | 0 | NA | NA |
| cr4_11 | NA | NA | 0 | NA | NA | 0 | NA | NA |
| cr8_2 | NA | NA | 0 | NA | NA | 0 | NA | NA |
| cr6_19 | NA | NA | 0 | NA | NA | 0 | NA | NA |
| cr9_59 | NA | NA | 0 | NA | NA | 0 | NA | NA |
| cr8_10 | NA | NA | 0 | NA | NA | 0 | NA | NA |
| ar3_2 | NA | NA | 0 | NA | NA | 0 | NA | NA |
| cr9_56 | NA | NA | 0 | NA | NA | 0 | NA | NA |
| tm4_14 | NA | NA | 0 | NA | NA | 0 | NA | NA |
| tm5_2 | NA | NA | 0 | NA | NA | 0 | NA | NA |
| cr9_48 | NA | NA | 0 | NA | NA | 0 | NA | NA |
| cr5_5 | NA | NA | 0 | NA | NA | 0 | NA | NA |
| cr9_32 | NA | NA | 0 | NA | NA | 0 | NA | NA |
| tm6_12 | node_69 | 4 | 3 | NA | NA | 0 | FALSE | NA |
| cr2_8 | node_1 | 0 | 2 | NA | NA | 0 | FALSE | NA |
| cr1_23 | NA | NA | 0 | NA | NA | 0 | NA | NA |
| tm8_2 | NA | NA | 0 | NA | NA | 0 | NA | NA |
| cr3_21 | node_17 | 2 | 5 | NA | NA | 0 | FALSE | NA |
| tm3_6 | NA | NA | 0 | NA | NA | 0 | NA | NA |
| ar6_20 | NA | NA | 0 | NA | NA | 0 | NA | NA |
| cr9_39 | NA | NA | 0 | NA | NA | 0 | NA | NA |
| ar1_17 | node_1 | 0 | 2 | NA | NA | 0 | FALSE | NA |
| tm8_7 | NA | NA | 0 | NA | NA | 0 | NA | NA |
| cr2_15 | node_18 | 3 | 4 | NA | NA | 0 | FALSE | NA |
| tm3_8 | NA | NA | 0 | NA | NA | 0 | NA | NA |
| ar4_13 | node_17 | 2 | 7 | NA | NA | 0 | FALSE | NA |
| ar6_25 | NA | NA | 0 | NA | NA | 0 | NA | NA |
| tm4_17 | NA | NA | 0 | NA | NA | 0 | NA | NA |
| tm1_5 | node_1 | 0 | 1 | NA | NA | 0 | FALSE | NA |
| cr3_25 | node_2 | 1 | 3 | node_2 | 1 | 1 | FALSE | 0 |
| tm3_9 | NA | NA | 0 | NA | NA | 0 | NA | NA |
| tm6_21 | NA | NA | 0 | NA | NA | 0 | NA | NA |
| cr3_18 | node_1 | 0 | 1 | NA | NA | 0 | FALSE | NA |
| cr1_28 | NA | NA | 0 | NA | NA | 0 | NA | NA |
| tm7_2 | NA | NA | 0 | NA | NA | 0 | NA | NA |
| cr9_36 | NA | NA | 0 | NA | NA | 0 | NA | NA |
| cr9_42 | NA | NA | 0 | NA | NA | 0 | NA | NA |
| ar5_15 | DQ316067.1 | 3 | 6 | NA | NA | 0 | FALSE | NA |
| cr4_15 | NA | NA | 0 | NA | NA | 0 | NA | NA |
| cr2_14 | node_1 | 0 | 3 | NA | NA | 0 | FALSE | NA |
| cr1_13 | NA | NA | 0 | NA | NA | 0 | NA | NA |
| cr3_26 | node_17 | 2 | 8 | node_17 | 2 | 3 | FALSE | 0 |
| cr8_16 | NA | NA | 0 | NA | NA | 0 | NA | NA |
| ar4_33 | NA | NA | 0 | NA | NA | 0 | NA | NA |
| cr9_4 | NA | NA | 0 | NA | NA | 0 | NA | NA |
| ar3_14 | NA | NA | 0 | NA | NA | 0 | NA | NA |
| cr4_19 | NA | NA | 0 | NA | NA | 0 | NA | NA |
| tm4_4 | node_1 | 0 | 1 | NA | NA | 0 | FALSE | NA |
| ar1_3 | node_60 | 2 | 47 | node_60 | 2 | 15 | FALSE | 0 |
| cr4_22 | node_3 | 2 | 22 | node_2 | 1 | 7 | TRUE | 1 |
| ar5_11 | node_57 | 9 | 62 | node_17 | 2 | 16 | TRUE | 7 |
| ar6_19 | NA | NA | 0 | NA | NA | 0 | NA | NA |
| cr8_7 | NA | NA | 0 | NA | NA | 0 | NA | NA |
| cr4_6 | NA | NA | 0 | NA | NA | 0 | NA | NA |
| ar5_25 | node_2 | 1 | 1 | node_2 | 1 | 1 | FALSE | 0 |
| ar2_28 | NA | NA | 0 | NA | NA | 0 | NA | NA |
| cr8_33 | node_30 | 8 | 107 | node_17 | 2 | 26 | TRUE | 6 |
| ar6_6 | node_1 | 0 | 1 | NA | NA | 0 | FALSE | NA |
| ar4_24 | NA | NA | 0 | NA | NA | 0 | NA | NA |
| cr5_12 | NA | NA | 0 | NA | NA | 0 | NA | NA |
| cr6_4 | node_1 | 0 | 15 | NA | NA | 0 | FALSE | NA |
| cr8_35 | DQ316067.1 | 3 | 288 | MG334271.1 | 7 | 68 | FALSE | 6 |
| cr4_17 | NA | NA | 0 | NA | NA | 0 | NA | NA |
| cr3_24 | NA | NA | 0 | NA | NA | 0 | NA | NA |
| cr9_15 | NA | NA | 0 | NA | NA | 0 | NA | NA |
| ar3_24 | NA | NA | 0 | NA | NA | 0 | NA | NA |
| ar5_4 | NA | NA | 0 | NA | NA | 0 | NA | NA |
| tm7_16 | NA | NA | 0 | NA | NA | 0 | NA | NA |
| cr6_15 | NA | NA | 0 | NA | NA | 0 | NA | NA |
| cr6_20 | NA | NA | 0 | NA | NA | 0 | NA | NA |
| cr9_70 | NA | NA | 0 | NA | NA | 0 | NA | NA |
| cr2_11 | NA | NA | 0 | NA | NA | 0 | NA | NA |
| ar6_21 | NA | NA | 0 | NA | NA | 0 | NA | NA |
| tm7_1 | NA | NA | 0 | NA | NA | 0 | NA | NA |
| cr5_13 | node_1 | 0 | 17 | node_17 | 2 | 5 | TRUE | 2 |
| ar4_15 | node_21 | 5 | 32 | node_17 | 2 | 6 | TRUE | 3 |
| ar1_22 | NA | NA | 0 | NA | NA | 0 | NA | NA |
| ar4_22 | node_40 | 9 | 29 | node_17 | 2 | 6 | TRUE | 7 |
| ar1_9 | node_1 | 0 | 1 | NA | NA | 0 | FALSE | NA |
| cr1_11 | NA | NA | 0 | NA | NA | 0 | NA | NA |
| cr4_31 | node_1 | 0 | 1 | NA | NA | 0 | FALSE | NA |
| ar6_1 | node_17 | 2 | 4 | NA | NA | 0 | FALSE | NA |
| cr6_27 | NA | NA | 0 | NA | NA | 0 | NA | NA |
| tm3_2 | node_18 | 3 | 8 | node_17 | 2 | 1 | TRUE | 1 |
| ar5_29 | NA | NA | 0 | NA | NA | 0 | NA | NA |
| ar4_10 | node_2 | 1 | 7 | node_2 | 1 | 1 | FALSE | 0 |
| cr8_28 | node_1 | 0 | 2 | NA | NA | 0 | FALSE | NA |
| cr3_4 | node_32 | 5 | 2 | NA | NA | 0 | FALSE | NA |
| tm1_9 | NA | NA | 0 | NA | NA | 0 | NA | NA |
| cr8_40 | NA | NA | 0 | NA | NA | 0 | NA | NA |
| cr8_8 | NA | NA | 0 | NA | NA | 0 | NA | NA |
| cr4_26 | NA | NA | 0 | NA | NA | 0 | NA | NA |
| ar6_2 | node_17 | 2 | 3 | NA | NA | 0 | FALSE | NA |
| cr8_19 | NA | NA | 0 | NA | NA | 0 | NA | NA |
| cr9_17 | NA | NA | 0 | NA | NA | 0 | NA | NA |
| cr9_60 | NA | NA | 0 | NA | NA | 0 | NA | NA |
| cr7_6 | NA | NA | 0 | NA | NA | 0 | NA | NA |
| ar4_1 | node_1 | 0 | 1 | NA | NA | 0 | FALSE | NA |
| ar4_8 | NA | NA | 0 | NA | NA | 0 | NA | NA |
| cr3_5 | NA | NA | 0 | NA | NA | 0 | NA | NA |
| cr7_13 | NA | NA | 0 | NA | NA | 0 | NA | NA |
| cr9_20 | NA | NA | 0 | NA | NA | 0 | NA | NA |
| cr7_3 | NA | NA | 0 | NA | NA | 0 | NA | NA |
| cr5_21 | NA | NA | 0 | NA | NA | 0 | NA | NA |
| cr5_17 | node_60 | 2 | 4 | NA | NA | 0 | FALSE | NA |
| tm2_6 | NA | NA | 0 | NA | NA | 0 | NA | NA |
| cr6_22 | NA | NA | 0 | NA | NA | 0 | NA | NA |
| ar1_2 | node_62 | 3 | 44 | node_62 | 3 | 8 | FALSE | 0 |
| ar5_1 | node_17 | 2 | 5 | node_2 | 1 | 2 | TRUE | 1 |
| ar3_3 | node_1 | 0 | 1 | NA | NA | 0 | FALSE | NA |
| cr4_30 | node_26 | 4 | 7 | node_17 | 2 | 2 | TRUE | 2 |
| cr9_23 | node_1 | 0 | 1 | NA | NA | 0 | FALSE | NA |
| tm4_10 | NA | NA | 0 | NA | NA | 0 | NA | NA |
| cr1_29 | NA | NA | 0 | NA | NA | 0 | NA | NA |
| ar3_1 | KX027564.1 | 5 | 3 | KX027564.1 | 5 | 1 | FALSE | 0 |
| tm7_6 | NA | NA | 0 | NA | NA | 0 | NA | NA |
| ar4_25 | NA | NA | 0 | NA | NA | 0 | NA | NA |
| cr8_24 | NA | NA | 0 | NA | NA | 0 | NA | NA |
| cr7_17 | NA | NA | 0 | NA | NA | 0 | NA | NA |
| cr2_24 | NA | NA | 0 | NA | NA | 0 | NA | NA |
| ar3_20 | NA | NA | 0 | NA | NA | 0 | NA | NA |
| ar4_4 | node_1 | 0 | 2 | NA | NA | 0 | FALSE | NA |
| tm6_10 | node_60 | 2 | 3 | NA | NA | 0 | FALSE | NA |
| ar6_30 | NA | NA | 0 | NA | NA | 0 | NA | NA |
| cr9_28 | node_3 | 2 | 5 | NA | NA | 0 | FALSE | NA |
| cr5_27 | NA | NA | 0 | NA | NA | 0 | NA | NA |
| cr9_66 | NA | NA | 0 | NA | NA | 0 | NA | NA |
| cr6_1 | NA | NA | 0 | NA | NA | 0 | NA | NA |
| ar2_12 | NA | NA | 0 | NA | NA | 0 | NA | NA |
| cr1_20 | NA | NA | 0 | NA | NA | 0 | NA | NA |
| cr3_22 | NA | NA | 0 | NA | NA | 0 | NA | NA |
| cr4_14 | node_3 | 2 | 69 | node_3 | 2 | 13 | FALSE | 0 |
| ar1_19 | node_1 | 0 | 1 | NA | NA | 0 | FALSE | NA |
| cr8_34 | node_17 | 2 | 19 | node_2 | 1 | 3 | TRUE | 1 |
| ar3_4 | NA | NA | 0 | NA | NA | 0 | NA | NA |
| cr7_7 | NA | NA | 0 | NA | NA | 0 | NA | NA |
| ar1_15 | NA | NA | 0 | NA | NA | 0 | NA | NA |
| ar3_34 | NA | NA | 0 | NA | NA | 0 | NA | NA |
| ar4_27 | NA | NA | 0 | NA | NA | 0 | NA | NA |
| cr9_69 | NA | NA | 0 | NA | NA | 0 | NA | NA |
| cr3_3 | node_17 | 2 | 13 | node_17 | 2 | 6 | FALSE | 0 |
| cr4_12 | NA | NA | 0 | NA | NA | 0 | NA | NA |
| tm2_1 | node_2 | 1 | 6 | NA | NA | 0 | FALSE | NA |
| cr9_65 | NA | NA | 0 | NA | NA | 0 | NA | NA |
| cr9_73 | NA | NA | 0 | NA | NA | 0 | NA | NA |
| cr2_29 | node_1 | 0 | 10 | NA | NA | 0 | FALSE | NA |
| cr6_16 | NA | NA | 0 | NA | NA | 0 | NA | NA |
| ar3_31 | NA | NA | 0 | NA | NA | 0 | NA | NA |
| ar1_28 | node_17 | 2 | 7 | node_17 | 2 | 2 | FALSE | 0 |
| cr4_16 | NA | NA | 0 | NA | NA | 0 | NA | NA |
| ar2_29 | node_1 | 0 | 2 | NA | NA | 0 | FALSE | NA |
| ar1_13 | node_2 | 1 | 1 | node_2 | 1 | 2 | FALSE | 0 |
| cr7_12 | NA | NA | 0 | NA | NA | 0 | NA | NA |
| cr8_39 | EU153451.1 | 5 | 135 | EU153451.1 | 5 | 22 | FALSE | 0 |
| cr9_35 | NA | NA | 0 | NA | NA | 0 | NA | NA |
| tm4_7 | node_36 | 9 | 21 | node_17 | 2 | 4 | TRUE | 7 |
| cr5_30 | NA | NA | 0 | NA | NA | 0 | NA | NA |
| ar4_19 | node_1 | 0 | 7 | NA | NA | 0 | FALSE | NA |
| cr8_3 | node_1 | 0 | 1 | NA | NA | 0 | FALSE | NA |
| cr7_21 | NA | NA | 0 | NA | NA | 0 | NA | NA |
| tm7_13 | NA | NA | 0 | NA | NA | 0 | NA | NA |
| ar4_30 | EU153452.1 | 6 | 12 | node_2 | 1 | 2 | TRUE | 5 |
| cr6_29 | NA | NA | 0 | NA | NA | 0 | NA | NA |
| ar5_7 | node_2 | 1 | 5 | node_2 | 1 | 3 | FALSE | 0 |
| cr1_2 | NA | NA | 0 | NA | NA | 0 | NA | NA |
| cr8_37 | node_59 | 1 | 43 | node_62 | 3 | 11 | TRUE | 2 |
| tm6_22 | NA | NA | 0 | NA | NA | 0 | NA | NA |
| tm9_3 | NA | NA | 0 | NA | NA | 0 | NA | NA |
| ar6_10 | node_29 | 7 | 5 | node_2 | 1 | 1 | TRUE | 6 |
| ar3_18 | NA | NA | 0 | NA | NA | 0 | NA | NA |
| cr4_10 | NA | NA | 0 | NA | NA | 0 | NA | NA |
| cr9_13 | NA | NA | 0 | NA | NA | 0 | NA | NA |
| ar5_23 | node_3 | 2 | 7 | node_18 | 3 | 3 | FALSE | 3 |
| cr3_28 | NA | NA | 0 | NA | NA | 0 | NA | NA |
| cr5_26 | node_1 | 0 | 1 | NA | NA | 0 | FALSE | NA |
| cr5_28 | node_17 | 2 | 4 | node_17 | 2 | 1 | FALSE | 0 |
| cr9_46 | NA | NA | 0 | NA | NA | 0 | NA | NA |
| cr8_13 | NA | NA | 0 | NA | NA | 0 | NA | NA |
| cr5_2 | node_1 | 0 | 6 | node_2 | 1 | 2 | TRUE | 1 |
| ar6_16 | NA | NA | 0 | NA | NA | 0 | NA | NA |
| cr7_25 | NA | NA | 0 | NA | NA | 0 | NA | NA |
| ar5_30 | NA | NA | 0 | NA | NA | 0 | NA | NA |
| cr5_22 | node_2 | 1 | 4 | node_2 | 1 | 1 | FALSE | 0 |
| ar5_19 | node_17 | 2 | 12 | node_17 | 2 | 5 | FALSE | 0 |
| cr4_8 | NA | NA | 0 | NA | NA | 0 | NA | NA |
| cr9_49 | NA | NA | 0 | NA | NA | 0 | NA | NA |
| ar6_31 | NA | NA | 0 | NA | NA | 0 | NA | NA |
| ar6_9 | node_1 | 0 | 1 | NA | NA | 0 | FALSE | NA |
| ar5_17 | node_32 | 5 | 150 | KX027495.1 | 7 | 41 | FALSE | 6 |
| ar1_29 | NA | NA | 0 | MF770243.1 | 10 | 1 | FALSE | NA |
| ar4_3 | NA | NA | 0 | NA | NA | 0 | NA | NA |
| cr4_18 | NA | NA | 0 | NA | NA | 0 | NA | NA |
| cr7_26 | NA | NA | 0 | NA | NA | 0 | NA | NA |
| tm6_5 | NA | NA | 0 | NA | NA | 0 | NA | NA |
| tm6_9 | NA | NA | 0 | NA | NA | 0 | NA | NA |
| tm9_10 | NA | NA | 0 | NA | NA | 0 | NA | NA |
| tm3_4 | NA | NA | 0 | NA | NA | 0 | NA | NA |
| ar2_1 | node_1 | 0 | 3 | NA | NA | 0 | FALSE | NA |
| cr1_26 | NA | NA | 0 | NA | NA | 0 | NA | NA |
| ar4_7 | NA | NA | 0 | NA | NA | 0 | NA | NA |
| cr1_8 | NA | NA | 0 | NA | NA | 0 | NA | NA |
| tm9_6 | node_1 | 0 | 3 | NA | NA | 0 | FALSE | NA |
| cr5_1 | NA | NA | 0 | NA | NA | 0 | NA | NA |
| tm8_6 | NA | NA | 0 | NA | NA | 0 | NA | NA |
| cr8_14 | NA | NA | 0 | NA | NA | 0 | NA | NA |
| ar5_20 | KX176768.1 | 5 | 3 | node_2 | 1 | 1 | FALSE | 6 |
| ar4_11 | node_19 | 4 | 7 | node_2 | 1 | 3 | TRUE | 3 |
| cr9_24 | NA | NA | 0 | NA | NA | 0 | NA | NA |
| ar4_21 | node_2 | 1 | 7 | node_2 | 1 | 1 | FALSE | 0 |
| ar6_22 | NA | NA | 0 | NA | NA | 0 | NA | NA |
| cr4_4 | NA | NA | 0 | NA | NA | 0 | NA | NA |
| cr9_1 | NA | NA | 0 | NA | NA | 0 | NA | NA |
| tm5_1 | NA | NA | 0 | NA | NA | 0 | NA | NA |
| ar3_7 | node_1 | 0 | 1 | NA | NA | 0 | FALSE | NA |
| cr3_15 | NA | NA | 0 | NA | NA | 0 | NA | NA |
| cr2_10 | NA | NA | 0 | NA | NA | 0 | NA | NA |
| tm4_12 | NA | NA | 0 | NA | NA | 0 | NA | NA |
| ar2_17 | node_5 | 4 | 8 | node_2 | 1 | 1 | TRUE | 3 |
| ar6_12 | node_37 | 8 | 161 | node_32 | 5 | 36 | TRUE | 3 |
| ar2_11 | NA | NA | 0 | NA | NA | 0 | NA | NA |
| ar1_25 | node_1 | 0 | 1 | NA | NA | 0 | FALSE | NA |
| tm6_14 | NA | NA | 0 | NA | NA | 0 | NA | NA |
| tm8_9 | NA | NA | 0 | NA | NA | 0 | NA | NA |
| tm6_11 | NA | NA | 0 | NA | NA | 0 | NA | NA |
| cr9_19 | NA | NA | 0 | NA | NA | 0 | NA | NA |
| tm7_3 | node_1 | 0 | 1 | NA | NA | 0 | FALSE | NA |
| cr2_28 | NA | NA | 0 | NA | NA | 0 | NA | NA |
| ar3_16 | NA | NA | 0 | NA | NA | 0 | NA | NA |
| cr7_15 | NA | NA | 0 | NA | NA | 0 | NA | NA |
| cr4_21 | node_3 | 2 | 10 | NA | NA | 0 | FALSE | NA |
| cr1_18 | NA | NA | 0 | NA | NA | 0 | NA | NA |
| ar4_29 | NA | NA | 0 | NA | NA | 0 | NA | NA |
| ar3_10 | NA | NA | 0 | NA | NA | 0 | NA | NA |
| cr9_47 | NA | NA | 0 | NA | NA | 0 | NA | NA |
| cr1_27 | NA | NA | 0 | NA | NA | 0 | NA | NA |
| cr9_30 | NA | NA | 0 | NA | NA | 0 | NA | NA |
| cr2_9 | NA | NA | 0 | NA | NA | 0 | NA | NA |
| cr9_55 | NA | NA | 0 | NA | NA | 0 | NA | NA |
| tm8_1 | NA | NA | 0 | NA | NA | 0 | NA | NA |
| cr3_16 | NA | NA | 0 | NA | NA | 0 | NA | NA |
| ar5_18 | node_33 | 6 | 451 | node_32 | 5 | 142 | TRUE | 1 |
| cr9_10 | NA | NA | 0 | NA | NA | 0 | NA | NA |
| cr8_31 | MG334279.1 | 13 | 76 | node_17 | 2 | 19 | TRUE | 11 |
| cr8_18 | NA | NA | 0 | NA | NA | 0 | NA | NA |
| cr8_12 | NA | NA | 0 | NA | NA | 0 | NA | NA |
| cr8_38 | EU153451.1 | 5 | 55 | node_60 | 2 | 16 | TRUE | 3 |
| ar6_17 | NA | NA | 0 | NA | NA | 0 | NA | NA |
| ar6_24 | NA | NA | 0 | NA | NA | 0 | NA | NA |
| cr5_7 | NA | NA | 0 | NA | NA | 0 | NA | NA |
| cr2_22 | node_17 | 2 | 52 | node_17 | 2 | 11 | FALSE | 0 |
| cr5_31 | node_32 | 5 | 76 | EU155210.1 | 9 | 20 | TRUE | 4 |
| ar1_32 | NA | NA | 0 | NA | NA | 0 | NA | NA |
| ar2_3 | node_1 | 0 | 1 | NA | NA | 0 | FALSE | NA |
| cr9_25 | NA | NA | 0 | NA | NA | 0 | NA | NA |
| ar1_20 | NA | NA | 0 | NA | NA | 0 | NA | NA |
| cr1_5 | NA | NA | 0 | NA | NA | 0 | NA | NA |
| ar6_18 | NA | NA | 0 | NA | NA | 0 | NA | NA |
| cr3_23 | node_29 | 7 | 91 | node_17 | 2 | 15 | TRUE | 5 |
| cr2_23 | node_32 | 5 | 25 | node_32 | 5 | 4 | FALSE | 0 |
| ar3_25 | NA | NA | 0 | NA | NA | 0 | NA | NA |
| ar3_30 | NA | NA | 0 | NA | NA | 0 | NA | NA |
| cr8_5 | NA | NA | 0 | NA | NA | 0 | NA | NA |
| tm4_11 | NA | NA | 0 | NA | NA | 0 | NA | NA |
| cr3_11 | NA | NA | 0 | NA | NA | 0 | NA | NA |
| cr6_7 | node_28 | 6 | 11 | NA | NA | 0 | FALSE | NA |
| cr7_18 | NA | NA | 0 | NA | NA | 0 | NA | NA |
| ar6_14 | NA | NA | 0 | NA | NA | 0 | NA | NA |
| cr2_13 | node_1 | 0 | 2 | NA | NA | 0 | FALSE | NA |
| ar5_32 | NA | NA | 0 | NA | NA | 0 | NA | NA |
| ar1_21 | NA | NA | 0 | NA | NA | 0 | NA | NA |
| cr7_28 | NA | NA | 0 | NA | NA | 0 | NA | NA |
| cr5_6 | NA | NA | 0 | NA | NA | 0 | NA | NA |
| cr8_27 | NA | NA | 0 | NA | NA | 0 | NA | NA |
| cr7_32 | NA | NA | 0 | NA | NA | 0 | NA | NA |
| ar1_33 | NA | NA | 0 | NA | NA | 0 | NA | NA |
| cr1_10 | NA | NA | 0 | NA | NA | 0 | NA | NA |
| ar1_12 | node_1 | 0 | 2 | NA | NA | 0 | FALSE | NA |
| tm1_3 | NA | NA | 0 | NA | NA | 0 | NA | NA |
| cr1_1 | NA | NA | 0 | NA | NA | 0 | NA | NA |
| cr3_32 | node_17 | 2 | 1 | node_2 | 1 | 1 | TRUE | 1 |
| cr7_16 | NA | NA | 0 | NA | NA | 0 | NA | NA |
| ar3_27 | NA | NA | 0 | NA | NA | 0 | NA | NA |
| ar2_22 | node_1 | 0 | 4 | node_2 | 1 | 1 | TRUE | 1 |
| cr1_19 | NA | NA | 0 | NA | NA | 0 | NA | NA |
| ar4_5 | node_1 | 0 | 1 | NA | NA | 0 | FALSE | NA |
| tm1_2 | node_2 | 1 | 5 | NA | NA | 0 | FALSE | NA |
| ar4_12 | node_32 | 5 | 18 | node_32 | 5 | 6 | FALSE | 0 |
| ar5_5 | node_1 | 0 | 1 | NA | NA | 0 | FALSE | NA |
| cr8_30 | node_17 | 2 | 49 | node_17 | 2 | 14 | FALSE | 0 |
| cr4_1 | node_1 | 0 | 1 | NA | NA | 0 | FALSE | NA |
| tm2_4 | NA | NA | 0 | NA | NA | 0 | NA | NA |
| cr3_2 | node_1 | 0 | 1 | NA | NA | 0 | FALSE | NA |
| tm6_2 | node_17 | 2 | 8 | node_2 | 1 | 2 | TRUE | 1 |
| tm6_4 | node_17 | 2 | 30 | node_30 | 8 | 9 | TRUE | 6 |
| ar6_29 | NA | NA | 0 | NA | NA | 0 | NA | NA |
| ar2_6 | node_1 | 0 | 5 | NA | NA | 0 | FALSE | NA |
| cr9_57 | NA | NA | 0 | NA | NA | 0 | NA | NA |
| cr5_18 | NA | NA | 0 | NA | NA | 0 | NA | NA |
| ar3_29 | NA | NA | 0 | NA | NA | 0 | NA | NA |
| tm9_12 | NA | NA | 0 | NA | NA | 0 | NA | NA |
| cr8_32 | NA | NA | 0 | NA | NA | 0 | NA | NA |
| cr9_16 | NA | NA | 0 | NA | NA | 0 | NA | NA |
| cr4_5 | NA | NA | 0 | NA | NA | 0 | NA | NA |
| cr8_20 | NA | NA | 0 | NA | NA | 0 | NA | NA |
| ar2_23 | node_5 | 4 | 47 | node_5 | 4 | 9 | FALSE | 0 |
| ar1_14 | node_26 | 4 | 16 | node_2 | 1 | 3 | TRUE | 3 |
| ar2_27 | node_1 | 0 | 5 | NA | NA | 0 | FALSE | NA |
| tm6_1 | node_2 | 1 | 13 | node_2 | 1 | 3 | FALSE | 0 |
| ar3_8 | NA | NA | 0 | NA | NA | 0 | NA | NA |
| cr9_31 | NA | NA | 0 | NA | NA | 0 | NA | NA |
| tm7_14 | NA | NA | 0 | NA | NA | 0 | NA | NA |
| cr9_29 | NA | NA | 0 | NA | NA | 0 | NA | NA |
| tm7_4 | NA | NA | 0 | NA | NA | 0 | NA | NA |
| ar2_2 | NA | NA | 0 | NA | NA | 0 | NA | NA |
| ar3_17 | NA | NA | 0 | NA | NA | 0 | NA | NA |
| ar4_26 | NA | NA | 0 | NA | NA | 0 | NA | NA |
| ar3_21 | node_1 | 0 | 1 | NA | NA | 0 | FALSE | NA |
| ar2_32 | NA | NA | 0 | NA | NA | 0 | NA | NA |
| cr5_16 | NA | NA | 0 | NA | NA | 0 | NA | NA |
| ar6_27 | NA | NA | 0 | NA | NA | 0 | NA | NA |
| tm6_20 | NA | NA | 0 | NA | NA | 0 | NA | NA |
| ar3_12 | NA | NA | 0 | NA | NA | 0 | NA | NA |
| ar2_16 | NA | NA | 0 | NA | NA | 0 | NA | NA |
| ar2_5 | node_1 | 0 | 2 | NA | NA | 0 | FALSE | NA |
| cr8_9 | NA | NA | 0 | NA | NA | 0 | NA | NA |
| cr9_27 | node_1 | 0 | 1 | NA | NA | 0 | FALSE | NA |
| ar4_28 | NA | NA | 0 | NA | NA | 0 | NA | NA |
| cr9_34 | NA | NA | 0 | NA | NA | 0 | NA | NA |
| ar1_11 | node_2 | 1 | 2 | NA | NA | 0 | FALSE | NA |
| cr8_21 | NA | NA | 0 | NA | NA | 0 | NA | NA |
| cr8_11 | NA | NA | 0 | NA | NA | 0 | NA | NA |
| tm4_16 | NA | NA | 0 | NA | NA | 0 | NA | NA |
| tm6_24 | NA | NA | 0 | NA | NA | 0 | NA | NA |
| ar3_5 | node_1 | 0 | 3 | NA | NA | 0 | FALSE | NA |
| cr9_22 | NA | NA | 0 | NA | NA | 0 | NA | NA |
| cr3_9 | NA | NA | 0 | NA | NA | 0 | NA | NA |
| cr9_8 | NA | NA | 0 | NA | NA | 0 | NA | NA |
| cr9_58 | NA | NA | 0 | NA | NA | 0 | NA | NA |
| ar2_19 | NA | NA | 0 | NA | NA | 0 | NA | NA |
| ar4_32 | NA | NA | 0 | NA | NA | 0 | NA | NA |
| cr6_30 | NA | NA | 0 | NA | NA | 0 | NA | NA |
| cr2_17 | node_1 | 0 | 4 | NA | NA | 0 | FALSE | NA |
| cr6_23 | NA | NA | 0 | NA | NA | 0 | NA | NA |
| ar1_8 | NA | NA | 0 | NA | NA | 0 | NA | NA |
| tm2_2 | node_17 | 2 | 3 | NA | NA | 0 | FALSE | NA |
| cr8_25 | NA | NA | 0 | NA | NA | 0 | NA | NA |
| cr6_14 | node_17 | 2 | 12 | node_17 | 2 | 2 | FALSE | 0 |
| tm4_13 | node_60 | 2 | 10 | node_59 | 1 | 2 | TRUE | 1 |
| cr2_5 | NA | NA | 0 | NA | NA | 0 | NA | NA |
| cr7_10 | NA | NA | 0 | NA | NA | 0 | NA | NA |
| cr9_74 | NA | NA | 0 | NA | NA | 0 | NA | NA |
| ar5_3 | node_17 | 2 | 8 | node_2 | 1 | 2 | TRUE | 1 |
| tm7_7 | node_1 | 0 | 1 | NA | NA | 0 | FALSE | NA |
| cr1_6 | NA | NA | 0 | NA | NA | 0 | NA | NA |
| cr1_25 | NA | NA | 0 | NA | NA | 0 | NA | NA |
| tm8_8 | NA | NA | 0 | NA | NA | 0 | NA | NA |
| ar4_18 | node_26 | 4 | 14 | NA | NA | 0 | FALSE | NA |
| cr6_13 | node_2 | 1 | 4 | node_2 | 1 | 1 | FALSE | 0 |
| ar5_16 | NA | NA | 0 | NA | NA | 0 | NA | NA |
| ar4_14 | node_36 | 9 | 19 | node_17 | 2 | 3 | TRUE | 7 |
| ar5_2 | node_17 | 2 | 1 | node_17 | 2 | 1 | FALSE | 0 |
| cr5_8 | NA | NA | 0 | NA | NA | 0 | NA | NA |
| cr2_31 | NA | NA | 0 | NA | NA | 0 | NA | NA |
| ar2_15 | NA | NA | 0 | NA | NA | 0 | NA | NA |
| cr2_4 | node_1 | 0 | 5 | node_60 | 2 | 1 | TRUE | 2 |
| cr6_5 | NA | NA | 0 | NA | NA | 0 | NA | NA |
| cr1_17 | NA | NA | 0 | NA | NA | 0 | NA | NA |
| cr6_3 | node_1 | 0 | 1 | NA | NA | 0 | FALSE | NA |
| cr1_21 | NA | NA | 0 | NA | NA | 0 | NA | NA |
| cr3_17 | NA | NA | 0 | NA | NA | 0 | NA | NA |
| ar1_31 | NA | NA | 0 | NA | NA | 0 | NA | NA |
| cr9_51 | NA | NA | 0 | NA | NA | 0 | NA | NA |
| tm3_7 | node_1 | 0 | 1 | NA | NA | 0 | FALSE | NA |
| tm7_5 | node_59 | 1 | 4 | node_59 | 1 | 2 | FALSE | 0 |
| cr2_30 | node_1 | 0 | 1 | NA | NA | 0 | FALSE | NA |
| ar3_32 | NA | NA | 0 | NA | NA | 0 | NA | NA |
| ar4_2 | NA | NA | 0 | NA | NA | 0 | NA | NA |
| tm4_1 | node_18 | 3 | 14 | node_2 | 1 | 7 | TRUE | 2 |
| tm9_16 | NA | NA | 0 | NA | NA | 0 | NA | NA |
| cr1_12 | NA | NA | 0 | NA | NA | 0 | NA | NA |
| tm1_8 | NA | NA | 0 | NA | NA | 0 | NA | NA |
| cr1_16 | NA | NA | 0 | NA | NA | 0 | NA | NA |
| cr5_23 | node_18 | 3 | 27 | node_18 | 3 | 9 | FALSE | 0 |
| ar3_22 | node_1 | 0 | 1 | NA | NA | 0 | FALSE | NA |
| cr6_8 | NA | NA | 0 | NA | NA | 0 | NA | NA |
| cr8_6 | NA | NA | 0 | NA | NA | 0 | NA | NA |
| tm4_5 | node_26 | 4 | 13 | node_18 | 3 | 4 | TRUE | 1 |
| ar6_23 | NA | NA | 0 | NA | NA | 0 | NA | NA |
| ar1_7 | node_1 | 0 | 1 | NA | NA | 0 | FALSE | NA |
| ar3_33 | NA | NA | 0 | NA | NA | 0 | NA | NA |
| ar4_16 | NA | NA | 0 | NA | NA | 0 | NA | NA |
| cr4_13 | NA | NA | 0 | NA | NA | 0 | NA | NA |
| cr3_7 | NA | NA | 0 | NA | NA | 0 | NA | NA |
| cr8_17 | NA | NA | 0 | NA | NA | 0 | NA | NA |
| tm6_6 | node_1 | 0 | 3 | NA | NA | 0 | FALSE | NA |
| ar1_10 | node_17 | 2 | 3 | node_17 | 2 | 2 | FALSE | 0 |
| cr2_21 | node_40 | 9 | 28 | node_2 | 1 | 9 | TRUE | 8 |
| cr8_29 | node_26 | 4 | 27 | node_18 | 3 | 6 | TRUE | 1 |
| cr5_19 | NA | NA | 0 | NA | NA | 0 | NA | NA |
| tm9_13 | KX027567.1 | 5 | 3 | KX027567.1 | 5 | 2 | FALSE | 0 |
| ar4_17 | node_17 | 2 | 7 | node_17 | 2 | 2 | FALSE | 0 |
| tm1_7 | NA | NA | 0 | NA | NA | 0 | NA | NA |
| cr4_20 | node_3 | 2 | 14 | NA | NA | 0 | FALSE | NA |
| cr6_25 | NA | NA | 0 | NA | NA | 0 | NA | NA |
| tm1_4 | NA | NA | 0 | NA | NA | 0 | NA | NA |
| tm3_3 | NA | NA | 0 | NA | NA | 0 | NA | NA |
| ar5_26 | NA | NA | 0 | NA | NA | 0 | NA | NA |
| cr9_54 | NA | NA | 0 | NA | NA | 0 | NA | NA |
| cr3_1 | node_17 | 2 | 7 | node_17 | 2 | 1 | FALSE | 0 |
| ar5_31 | node_29 | 7 | 12 | node_18 | 3 | 4 | TRUE | 4 |
| cr7_8 | NA | NA | 0 | NA | NA | 0 | NA | NA |
| tm7_8 | node_33 | 6 | 7 | node_2 | 1 | 2 | TRUE | 5 |
| cr9_40 | NA | NA | 0 | NA | NA | 0 | NA | NA |
| tm4_8 | NA | NA | 0 | NA | NA | 0 | NA | NA |
| ar2_31 | NA | NA | 0 | NA | NA | 0 | NA | NA |
| cr8_26 | NA | NA | 0 | NA | NA | 0 | NA | NA |
| ar2_24 | node_1 | 0 | 1 | NA | NA | 0 | FALSE | NA |
| cr2_19 | NA | NA | 0 | NA | NA | 0 | NA | NA |
| ar5_24 | NA | NA | 0 | NA | NA | 0 | NA | NA |
| cr7_24 | NA | NA | 0 | NA | NA | 0 | NA | NA |
| tm6_15 | NA | NA | 0 | NA | NA | 0 | NA | NA |
| cr9_38 | NA | NA | 0 | NA | NA | 0 | NA | NA |
| cr9_61 | NA | NA | 0 | NA | NA | 0 | NA | NA |
| ar6_7 | node_18 | 3 | 7 | node_2 | 1 | 2 | TRUE | 2 |
| ar1_1 | node_60 | 2 | 8 | NA | NA | 0 | FALSE | NA |
| cr9_3 | NA | NA | 0 | NA | NA | 0 | NA | NA |
| ar1_34 | NA | NA | 0 | NA | NA | 0 | NA | NA |
| cr3_14 | NA | NA | 0 | NA | NA | 0 | NA | NA |
| ar5_8 | NA | NA | 0 | NA | NA | 0 | NA | NA |
| ar1_5 | NA | NA | 0 | NA | NA | 0 | NA | NA |
| ar2_14 | node_3 | 2 | 6 | node_3 | 2 | 2 | FALSE | 0 |
| cr9_12 | NA | NA | 0 | NA | NA | 0 | NA | NA |
| tm1_1 | node_1 | 0 | 5 | node_2 | 1 | 1 | TRUE | 1 |
| tm9_4 | node_26 | 4 | 24 | node_26 | 4 | 7 | FALSE | 0 |
| cr2_1 | node_1 | 0 | 1 | NA | NA | 0 | FALSE | NA |
| cr4_2 | NA | NA | 0 | NA | NA | 0 | NA | NA |
| cr9_11 | NA | NA | 0 | NA | NA | 0 | NA | NA |
| cr1_24 | NA | NA | 0 | NA | NA | 0 | NA | NA |
| cr3_6 | NA | NA | 0 | NA | NA | 0 | NA | NA |
| ar2_18 | node_2 | 1 | 1 | node_2 | 1 | 1 | FALSE | 0 |
| cr1_9 | NA | NA | 0 | NA | NA | 0 | NA | NA |
| cr5_9 | EU153458.1 | 8 | 15 | node_17 | 2 | 3 | TRUE | 6 |
| cr1_7 | NA | NA | 0 | NA | NA | 0 | NA | NA |
| ar5_14 | node_26 | 4 | 42 | node_54 | 6 | 9 | TRUE | 2 |
| cr2_6 | node_72 | 6 | 10 | NA | NA | 0 | FALSE | NA |
| cr6_17 | NA | NA | 0 | NA | NA | 0 | NA | NA |
| cr6_31 | NA | NA | 0 | NA | NA | 0 | NA | NA |
| cr7_30 | NA | NA | 0 | NA | NA | 0 | NA | NA |
| tm6_18 | NA | NA | 0 | NA | NA | 0 | NA | NA |
| ar2_25 | node_2 | 1 | 12 | node_2 | 1 | 3 | FALSE | 0 |
| ar6_11 | NA | NA | 0 | NA | NA | 0 | NA | NA |
| ar1_26 | node_2 | 1 | 2 | node_2 | 1 | 1 | FALSE | 0 |
| cr6_9 | node_2 | 1 | 3 | node_2 | 1 | 2 | FALSE | 0 |
| ar5_12 | MG334273.1 | 10 | 4 | MG334273.1 | 10 | 1 | FALSE | 0 |
| cr6_26 | NA | NA | 0 | NA | NA | 0 | NA | NA |
| ar2_30 | node_1 | 0 | 3 | NA | NA | 0 | FALSE | NA |
| cr3_30 | node_1 | 0 | 1 | NA | NA | 0 | FALSE | NA |
| cr4_32 | NA | NA | 0 | NA | NA | 0 | NA | NA |
| ar2_4 | node_2 | 1 | 1 | NA | NA | 0 | FALSE | NA |
| cr5_29 | NA | NA | 0 | NA | NA | 0 | NA | NA |
| ar2_26 | node_1 | 0 | 2 | NA | NA | 0 | FALSE | NA |
| ar3_11 | NA | NA | 0 | NA | NA | 0 | NA | NA |
| ar4_20 | NA | NA | 0 | NA | NA | 0 | NA | NA |
| cr9_14 | NA | NA | 0 | NA | NA | 0 | NA | NA |
| cr6_18 | NA | NA | 0 | NA | NA | 0 | NA | NA |
| cr1_30 | NA | NA | 0 | NA | NA | 0 | NA | NA |
| cr9_71 | NA | NA | 0 | NA | NA | 0 | NA | NA |
| cr9_72 | NA | NA | 0 | NA | NA | 0 | NA | NA |
| cr6_11 | NA | NA | 0 | NA | NA | 0 | NA | NA |
| cr9_9 | NA | NA | 0 | NA | NA | 0 | NA | NA |
| cr1_32 | NA | NA | 0 | NA | NA | 0 | NA | NA |
| cr9_50 | NA | NA | 0 | NA | NA | 0 | NA | NA |
| ar5_9 | node_1 | 0 | 2 | node_2 | 1 | 2 | TRUE | 1 |
| cr6_12 | node_2 | 1 | 9 | NA | NA | 0 | FALSE | NA |
| cr4_24 | NA | NA | 0 | NA | NA | 0 | NA | NA |
| cr2_20 | node_1 | 0 | 5 | NA | NA | 0 | FALSE | NA |
| cr5_10 | node_17 | 2 | 12 | node_2 | 1 | 4 | TRUE | 1 |
| cr5_25 | node_1 | 0 | 1 | NA | NA | 0 | FALSE | NA |
| tm6_16 | NA | NA | 0 | NA | NA | 0 | NA | NA |
| tm7_10 | NA | NA | 0 | NA | NA | 0 | NA | NA |
| cr7_4 | NA | NA | 0 | NA | NA | 0 | NA | NA |
| tm9_2 | node_17 | 2 | 11 | node_2 | 1 | 2 | TRUE | 1 |
| cr5_24 | NA | NA | 0 | NA | NA | 0 | NA | NA |
| ar4_31 | MF770243.1 | 10 | 26 | node_17 | 2 | 10 | TRUE | 8 |
| tm2_9 | NA | NA | 0 | NA | NA | 0 | NA | NA |
| ar3_19 | NA | NA | 0 | NA | NA | 0 | NA | NA |
| tm3_1 | node_1 | 0 | 4 | NA | NA | 0 | FALSE | NA |
| tm7_12 | NA | NA | 0 | NA | NA | 0 | NA | NA |
| ar2_20 | node_3 | 2 | 48 | node_3 | 2 | 10 | FALSE | 0 |
| cr1_15 | NA | NA | 0 | NA | NA | 0 | NA | NA |
| cr1_31 | NA | NA | 0 | NA | NA | 0 | NA | NA |
| tm6_13 | NA | NA | 0 | NA | NA | 0 | NA | NA |
| ar1_27 | node_1 | 0 | 1 | NA | NA | 0 | FALSE | NA |
| cr7_29 | NA | NA | 0 | NA | NA | 0 | NA | NA |
| ar2_13 | NA | NA | 0 | NA | NA | 0 | NA | NA |
| cr3_8 | NA | NA | 0 | NA | NA | 0 | NA | NA |
| cr1_14 | NA | NA | 0 | NA | NA | 0 | NA | NA |
| tm9_1 | node_1 | 0 | 3 | NA | NA | 0 | FALSE | NA |
| cr7_11 | NA | NA | 0 | NA | NA | 0 | NA | NA |
| cr9_63 | NA | NA | 0 | NA | NA | 0 | NA | NA |
| tm6_3 | node_32 | 5 | 14 | NA | NA | 0 | FALSE | NA |
| cr4_7 | NA | NA | 0 | NA | NA | 0 | NA | NA |
| cr6_32 | NA | NA | 0 | NA | NA | 0 | NA | NA |
| ar4_9 | node_17 | 2 | 5 | NA | NA | 0 | FALSE | NA |
| tm2_5 | NA | NA | 0 | NA | NA | 0 | NA | NA |
| tm4_9 | NA | NA | 0 | NA | NA | 0 | NA | NA |
| cr6_28 | NA | NA | 0 | NA | NA | 0 | NA | NA |
| cr9_45 | NA | NA | 0 | NA | NA | 0 | NA | NA |
| ar6_26 | NA | NA | 0 | NA | NA | 0 | NA | NA |
| cr7_5 | NA | NA | 0 | NA | NA | 0 | NA | NA |
| cr9_44 | NA | NA | 0 | NA | NA | 0 | NA | NA |
| ar3_23 | NA | NA | 0 | NA | NA | 0 | NA | NA |
| ar1_30 | node_1 | 0 | 3 | NA | NA | 0 | FALSE | NA |
| ar3_26 | NA | NA | 0 | NA | NA | 0 | NA | NA |
| cr5_15 | NA | NA | 0 | NA | NA | 0 | NA | NA |
| cr9_5 | NA | NA | 0 | NA | NA | 0 | NA | NA |
| ar4_34 | NA | NA | 0 | NA | NA | 0 | NA | NA |
| tm7_15 | NA | NA | 0 | NA | NA | 0 | NA | NA |
| cr3_20 | node_17 | 2 | 44 | node_17 | 2 | 17 | FALSE | 0 |
| cr4_9 | NA | NA | 0 | NA | NA | 0 | NA | NA |
| cr7_31 | NA | NA | 0 | NA | NA | 0 | NA | NA |
| ar3_13 | NA | NA | 0 | NA | NA | 0 | NA | NA |
| cr2_25 | NA | NA | 0 | NA | NA | 0 | NA | NA |
| ar4_36 | NA | NA | 0 | NA | NA | 0 | NA | NA |
| cr9_2 | NA | NA | 0 | NA | NA | 0 | NA | NA |
| tm1_6 | NA | NA | 0 | NA | NA | 0 | NA | NA |
| tm4_6 | node_32 | 5 | 49 | node_17 | 2 | 10 | TRUE | 3 |
| cr9_41 | NA | NA | 0 | NA | NA | 0 | NA | NA |
| ar4_6 | NA | NA | 0 | NA | NA | 0 | NA | NA |
| tm8_4 | NA | NA | 0 | NA | NA | 0 | NA | NA |
| ar5_22 | node_1 | 0 | 1 | NA | NA | 0 | FALSE | NA |
| cr3_13 | node_1 | 0 | 1 | NA | NA | 0 | FALSE | NA |
| cr4_3 | NA | NA | 0 | NA | NA | 0 | NA | NA |
| ar6_3 | node_17 | 2 | 8 | node_17 | 2 | 3 | FALSE | 0 |
| ar1_18 | NA | NA | 0 | NA | NA | 0 | NA | NA |
| ar6_32 | NA | NA | 0 | NA | NA | 0 | NA | NA |
| ar1_24 | NA | NA | 0 | NA | NA | 0 | NA | NA |
| cr9_53 | NA | NA | 0 | NA | NA | 0 | NA | NA |
| tm9_5 | node_2 | 1 | 6 | NA | NA | 0 | FALSE | NA |
| ar5_27 | NA | NA | 0 | NA | NA | 0 | NA | NA |
| ar1_4 | NA | NA | 0 | NA | NA | 0 | NA | NA |
| cr2_16 | NA | NA | 0 | NA | NA | 0 | NA | NA |
| ar5_6 | node_17 | 2 | 1 | NA | NA | 0 | FALSE | NA |
| cr3_19 | node_63 | 4 | 8 | node_63 | 4 | 1 | FALSE | 0 |
| cr7_1 | NA | NA | 0 | NA | NA | 0 | NA | NA |
| cr9_7 | NA | NA | 0 | NA | NA | 0 | NA | NA |
| ar6_28 | NA | NA | 0 | NA | NA | 0 | NA | NA |
| tm4_3 | node_17 | 2 | 8 | node_17 | 2 | 4 | FALSE | 0 |
| test | NA | NA | 0 | NA | NA | 0 | NA | NA |
| cr5_32 | NA | NA | 0 | NA | NA | 0 | NA | NA |
| tm8_3 | NA | NA | 0 | NA | NA | 0 | NA | NA |
| ar3_9 | NA | NA | 0 | NA | NA | 0 | NA | NA |
| cr8_23 | NA | NA | 0 | NA | NA | 0 | NA | NA |
| cr3_29 | node_28 | 6 | 13 | node_17 | 2 | 2 | TRUE | 4 |
| cr8_4 | NA | NA | 0 | NA | NA | 0 | NA | NA |
| cr9_26 | NA | NA | 0 | NA | NA | 0 | NA | NA |
| ar3_28 | NA | NA | 0 | NA | NA | 0 | NA | NA |
| tm9_7 | NA | NA | 0 | NA | NA | 0 | NA | NA |
| tm9_15 | NA | NA | 0 | NA | NA | 0 | NA | NA |
| cr3_27 | node_17 | 2 | 13 | NA | NA | 0 | FALSE | NA |
| cr5_20 | node_18 | 3 | 16 | node_18 | 3 | 3 | FALSE | 0 |
| tm6_23 | NA | NA | 0 | NA | NA | 0 | NA | NA |
| cr4_23 | NA | NA | 0 | NA | NA | 0 | NA | NA |
| cr7_27 | NA | NA | 0 | NA | NA | 0 | NA | NA |
| ar1_16 | NA | NA | 0 | NA | NA | 0 | NA | NA |
| ar6_4 | node_32 | 5 | 10 | node_32 | 5 | 1 | FALSE | 0 |
| ar1_23 | NA | NA | 0 | NA | NA | 0 | NA | NA |
| cr9_67 | NA | NA | 0 | NA | NA | 0 | NA | NA |
| tm8_16 | NA | NA | 0 | NA | NA | 0 | NA | NA |
| cr7_23 | NA | NA | 0 | NA | NA | 0 | NA | NA |
| tm2_3 | node_1 | 0 | 1 | NA | NA | 0 | FALSE | NA |
| tm7_11 | node_1 | 0 | 1 | NA | NA | 0 | FALSE | NA |
| ar3_15 | NA | NA | 0 | NA | NA | 0 | NA | NA |
| ar5_21 | node_1 | 0 | 1 | NA | NA | 0 | FALSE | NA |
| cr9_62 | NA | NA | 0 | NA | NA | 0 | NA | NA |
| cr4_27 | NA | NA | 0 | NA | NA | 0 | NA | NA |
| cr9_6 | NA | NA | 0 | NA | NA | 0 | NA | NA |
| tm6_17 | NA | NA | 0 | NA | NA | 0 | NA | NA |
| ar2_9 | node_1 | 0 | 1 | NA | NA | 0 | FALSE | NA |
| cr4_28 | NA | NA | 0 | NA | NA | 0 | NA | NA |
| cr9_21 | node_1 | 0 | 2 | NA | NA | 0 | FALSE | NA |
| cr1_4 | NA | NA | 0 | NA | NA | 0 | NA | NA |
| cr9_33 | NA | NA | 0 | NA | NA | 0 | NA | NA |
| ar6_5 | node_32 | 5 | 5 | node_32 | 5 | 1 | FALSE | 0 |
| tm2_8 | NA | NA | 0 | NA | NA | 0 | NA | NA |
| cr7_20 | NA | NA | 0 | NA | NA | 0 | NA | NA |
| tm7_17 | NA | NA | 0 | NA | NA | 0 | NA | NA |
| cr9_75 | NA | NA | 0 | NA | NA | 0 | NA | NA |
| cr4_29 | NA | NA | 0 | NA | NA | 0 | NA | NA |
| cr6_21 | NA | NA | 0 | NA | NA | 0 | NA | NA |
| cr7_22 | NA | NA | 0 | NA | NA | 0 | NA | NA |
| ar5_10 | node_17 | 2 | 36 | node_2 | 1 | 7 | TRUE | 1 |
| cr9_68 | NA | NA | 0 | NA | NA | 0 | NA | NA |
| cr6_24 | NA | NA | 0 | NA | NA | 0 | NA | NA |
| ar2_10 | node_1 | 0 | 1 | NA | NA | 0 | FALSE | NA |
| cr9_18 | NA | NA | 0 | NA | NA | 0 | NA | NA |
| cr5_4 | node_1 | 0 | 5 | NA | NA | 0 | FALSE | NA |
| cr3_31 | node_2 | 1 | 14 | node_2 | 1 | 1 | FALSE | 0 |
| cr3_10 | NA | NA | 0 | NA | NA | 0 | NA | NA |
| tm8_5 | node_1 | 0 | 2 | NA | NA | 0 | FALSE | NA |
| cr2_18 | NA | NA | 0 | NA | NA | 0 | NA | NA |
| cr7_19 | NA | NA | 0 | NA | NA | 0 | NA | NA |
| cr9_64 | NA | NA | 0 | NA | NA | 0 | NA | NA |
| cr5_3 | node_1 | 0 | 1 | NA | NA | 0 | FALSE | NA |
| cr8_22 | NA | NA | 0 | NA | NA | 0 | NA | NA |
| ar2_8 | NA | NA | 0 | NA | NA | 0 | NA | NA |
| ar6_8 | NA | NA | 0 | NA | NA | 0 | NA | NA |
| tm6_7 | NA | NA | 0 | NA | NA | 0 | NA | NA |
| cr5_11 | node_59 | 1 | 285 | node_59 | 1 | 69 | FALSE | 0 |
| tm5_8 | NA | NA | 0 | NA | NA | 0 | NA | NA |
| cr2_26 | NA | NA | 0 | NA | NA | 0 | NA | NA |
| ar5_28 | node_2 | 1 | 3 | NA | NA | 0 | FALSE | NA |
| cr1_22 | NA | NA | 0 | NA | NA | 0 | NA | NA |
| tm7_9 | node_32 | 5 | 10 | node_2 | 1 | 4 | TRUE | 4 |
| ar4_23 | node_24 | 8 | 55 | node_30 | 8 | 6 | FALSE | 10 |
| ar2_7 | node_18 | 3 | 2 | node_2 | 1 | 1 | TRUE | 2 |
| cr9_43 | NA | NA | 0 | NA | NA | 0 | NA | NA |
| cr7_9 | NA | NA | 0 | NA | NA | 0 | NA | NA |
| tm9_11 | NA | NA | 0 | NA | NA | 0 | NA | NA |
| tm2_7 | NA | NA | 0 | NA | NA | 0 | NA | NA |
| cr8_36 | AP008987.1 | 9 | 54 | node_63 | 4 | 12 | FALSE | 13 |
| ar5_13 | node_26 | 4 | 10 | node_2 | 1 | 3 | TRUE | 3 |
| cr3_12 | node_1 | 0 | 2 | NA | NA | 0 | FALSE | NA |
| cr6_10 | node_1 | 0 | 1 | NA | NA | 0 | FALSE | NA |
| cr2_27 | NA | NA | 0 | NA | NA | 0 | NA | NA |
| cr9_37 | NA | NA | 0 | NA | NA | 0 | NA | NA |
| cr9_52 | NA | NA | 0 | NA | NA | 0 | NA | NA |
| ar6_13 | NA | NA | 0 | NA | NA | 0 | NA | NA |
| cr7_14 | NA | NA | 0 | NA | NA | 0 | NA | NA |
| tm6_8 | NA | NA | 0 | NA | NA | 0 | NA | NA |
| tm9_14 | node_1 | 0 | 3 | NA | NA | 0 | FALSE | NA |
| ar2_21 | node_2 | 1 | 3 | node_2 | 1 | 1 | FALSE | 0 |
| tm6_19 | NA | NA | 0 | NA | NA | 0 | NA | NA |
| cr1_3 | NA | NA | 0 | NA | NA | 0 | NA | NA |
| cr2_3 | node_1 | 0 | 2 | NA | NA | 0 | FALSE | NA |
| tm5_3 | NA | NA | 0 | NA | NA | 0 | NA | NA |
| ar3_6 | NA | NA | 0 | NA | NA | 0 | NA | NA |
| ar6_15 | NA | NA | 0 | NA | NA | 0 | NA | NA |
| cr2_12 | NA | NA | 0 | NA | NA | 0 | NA | NA |

**Table S6: Panmap assigned reads validation**

| **Panmap assigned reads** | **Reads alignable to DQ188829.2** | **BLAST hits to Elephantidae mito** | **BLAST hits exclusively to Elephantidae mito genome** |
| --- | --- | --- | --- |
| 4,098 | 3,803 | 3,786 | 3,522 |
| **Competitive mapping assigned reads** | **Assigned by Panmap and competitive mapping** |  |  |
| 846 | 788 |  |  |
| **Average sequence divergence** | |  |  |
| **Panmap** | **Competitive mapping** |  |  |
| 7.12E-03 | 9.44E-03 |  |  |
